# Discovery of a synthetic small molecule targeting the central regulator of *Salmonella* pathogenicity

**DOI:** 10.1101/2024.04.29.591313

**Authors:** Abdelhakim Boudrioua, Joe D. Joiner, Iwan Grin, Thales Kronenberger, Vadim S. Korotkov, Wieland Steinchen, Alexander Kohler, Sophie Schminke, Julia-Christina Schulte, Michael Pietsch, Arun Naini, Simon Kalverkamp, Sven-Kevin Hotop, Travis Coyle, Claudio Piselli, Murray Coles, Katharina Rox, Matthias Marschal, Gert Bange, Antje Flieger, Antti Poso, Mark Brönstrup, Marcus D. Hartmann, Samuel Wagner

## Abstract

The enteric pathogen *Salmonella enterica* serovar Typhimurium relies on the activity of effector proteins to invade, replicate, and disseminate into host epithelial cells and other tissues, thereby causing disease. Secretion and injection of effector proteins into host cells is mediated by dedicated secretion systems, which hence represent major virulence determinants. Here, we report the identification of a synthetic small molecule with drug-like properties, C26, which suppresses the secretion of effector proteins, and consequently hinders bacterial invasion of eukaryotic cells. C26 binds to and inhibits HilD, the transcriptional regulator of the major secretion systems. While sharing the same binding pocket as the previously described long-chain fatty acid ligands, C26 inhibits HilD with a unique binding mode and a distinct mechanism. We provide evidence for target engagement within infected eukaryotic cells and present analogs with improved potency and suitability as scaffolds to develop anti-virulence agents against *Salmonella* infections in humans and animals.

## Main

The threat that antibiotic resistance poses to global public health is well recognized. In 2019, an estimated 1.27 million deaths were attributed to antibiotic resistant bacterial infections worldwide^1^. A promising strategy to circumvent antibiotic resistance is the development of drugs targeting virulence factors that are essential for bacterial pathogenesis but not for bacterial growth and viability^2–4^. In contrast to antibiotics that directly inhibit growth or kill the bacteria, non-lethal anti-virulence agents are thought to exert a reduced selective pressure for the development of resistant strains^5^, and preserve the commensal microbiota.

Non-typhoidal salmonellae (NTS) like *Salmonella enterica* subspecies *enterica* serovar Typhimurium are enteric pathogens causing inflammatory diarrhea that can develop into invasive non-typhoidal *Salmonella* (iNTS) infections once the bacteria invade the intestinal epithelium^6,7^. *S.* Typhimurium invasion of epithelial cells is mediated by secretion systems that are encoded on horizontally-acquired *Salmonella* pathogenicity islands (SPIs), and through which effector proteins are exported^8^. The first step in the pathogenesis of *S.* Typhimurium is adhesion to the host epithelial cells. A giant non-fimbrial adhesin is secreted through the SPI-4-encoded type I secretion system (T1SS) to initiate adhesion^9,10^. A type III secretion system encoded in SPI-1 (T3SS-1) is concomitantly assembled and enables the engulfment, followed by the internalization of bacteria into host epithelial cells^9^. Once inside the epithelial cells or inside macrophages, bacteria survive and replicate inside the *Salmonella*-containing vacuole (SCV)^11^, owing to a second T3SS encoded on SPI-2 (T3SS-2)^12^. Within macrophages, *S.* Typhimurium can disseminate in the bloodstream leading to a life-threatening systemic infection^13^.

The sequential activation of the different secretion systems requires a finely tuned regulation of the expression of SPI-encoded genes to coordinate the adhesion and injection of virulence factors in response to environmental signals. *S.* Typhimurium possesses virulence-associated signal transduction systems, which sustain a feed-forward regulatory loop formed by the three transcriptional regulators HilD, HilC, and RtsA^14^. These three regulators positively modulate each other’s expression, by binding to the promoter regions of their respective encoding genes^15,16^, and activity by forming homo- or heterodimers^17^. HilD is the main regulator through which the upstream signals feed into the regulatory network^14,18^. HilD positively regulates the transcriptional regulator HilA, known to be an activator of T3SS-1^19,20^ and T1SS^11^. HilD is also involved in the regulation of SPI-2 through the transcriptional activation of *ssrAB*^21^, which code for a two-component system. HilD is therefore considered the central regulator of *S.* Typhimurium invasion-related pathogenicity. A HilD-deficient strain is unable to activate the virulence genes encoded on SPIs^22^, to invade the caecal tissue, and to elicit inflammation in a mouse model of *S*. Typhimurium gastrointestinal infection^23^. The importance of the HilD-regulated SPI-1 and SPI-2 has also been demonstrated in a chicken infection model. Oral infection of 1-day old chickens with SPI-1- or SPI-2-deficient mutants resulted in a strong reduction of intestinal and systemic salmonellosis^24^.

Considering the critical role of T3SSs at different stages of *S.* Typhimurium pathogenicity, several anti-virulence agents targeting T3SS structural proteins^25–28^ and regulatory proteins^29–36^ have been identified. However, none of these inhibitors are actively being developed to treat *Salmonella* infections. In this study, we combined a virtual and phenotypic screen to identify inhibitors of T3SS-1 with drug-like properties. We identified compound C26 as a small molecule inhibiting protein secretion through T1SS, T3SS-1, and T3SS-2. Analysis of the mode of action revealed that C26 leads to a downregulation of all invasion-associated SPIs by targeting the transcriptional regulator HilD. As a result, treating bacteria with C26 impeded the invasion into host cells. We finally conducted a structure-activity relationship (SAR) analysis and uncovered analogs with improved potency.

### Identification of T3SS-1 inhibitors

To identify inhibitors of *S.* Typhimurium T3SS-1, we first set up an assay to monitor the secretion of the effector protein SipA (Fig. 1a). We then used molecular docking to computationally screen an Enamine library of ∼470,000 commercially available compounds against the major export apparatus protein SctV (InvA), resulting in the selection of 49 compounds (Fig. 1b). SipA secretion was monitored as reported previously^37^ to assess the inhibitory activity of the 49 compounds. The most potent compound, named C26, was a drug-like small molecule with a molecular weight of 397.3 Da (Fig. 1c and Supplementary Tab. 1).

**Figure 1.**
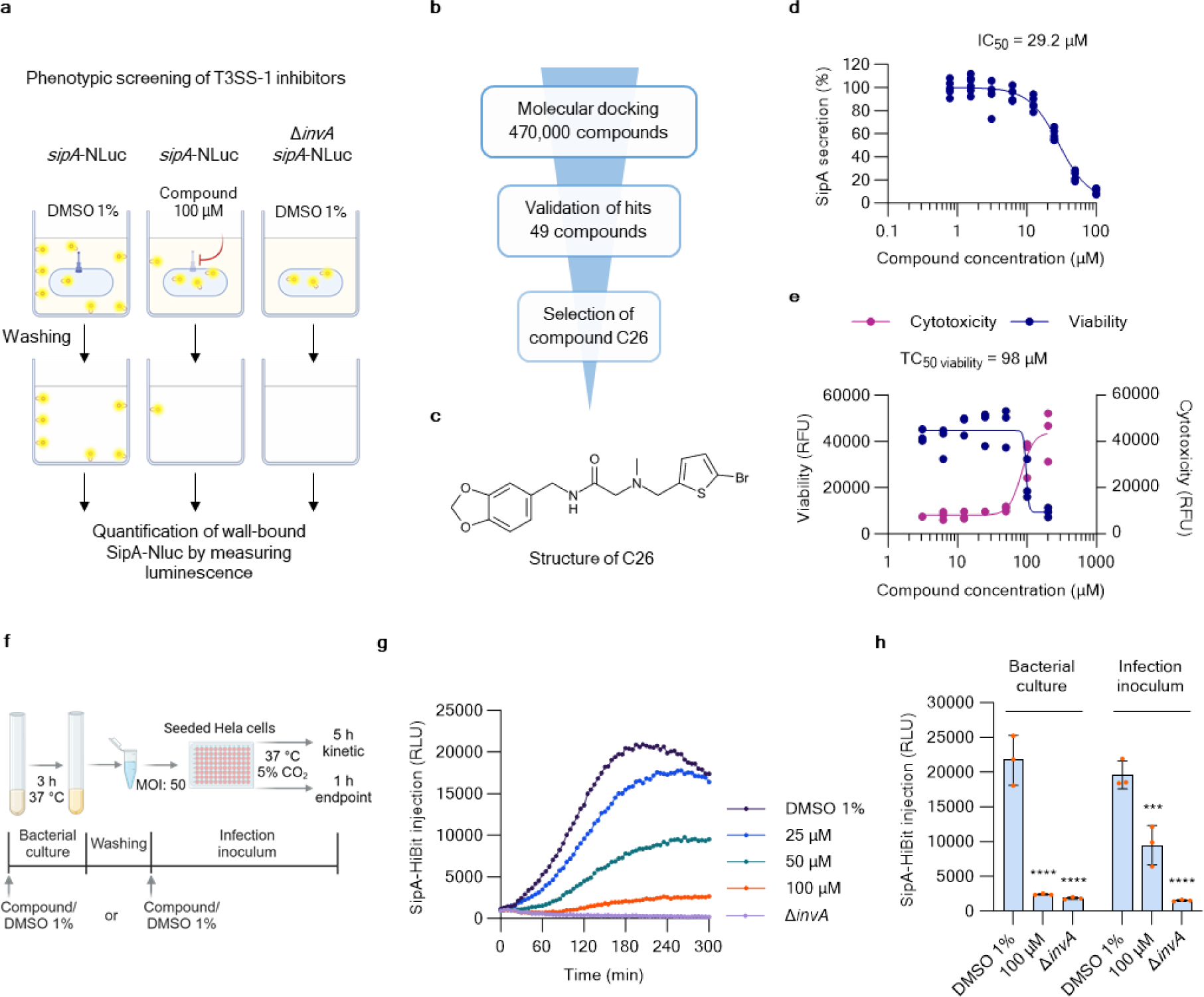
Identification of T3SS-1 inhibitors. **a)** NanoLuc Luciferase assay to screen T3SS-1 inhibitors in 384-well plates. Compounds were screened at a final concentration of 100 µM (n = 3 replicates). Cultures with strains *sipA*-NLuc and Δ*invA*, *sipA*-NLuc with 1% (v/v) DMSO were used as positive and negative controls, respectively. Figure created with BioRender. **b)** Screening workflow applied to identify T3SS-1 inhibitors. **c)** Chemical structure of compound C26. **d)** Dose-response curve of SipA secretion with increasing concentrations of C26. 100% relative SipA secretion corresponds to the luminescence intensity of the WT strain grown in the presence of 1% (v/v) DMSO. The relative SipA secretion of the Δ*invA* mutant was considered as the bottom. (n = 2 technical replicates within n = 3 biological replicates). **e)** *in vitro* toxicity in HeLa cells exposed to 100 µM C26 for 18 h using the ApoTox-Glo assay. Fluorescence was measured as a readout for viability 400_Ex_/505_Em_ and cytotoxicity 485_Ex_/520_Em_ (n = 3 biological replicates). **f)** Description of the experimental plan used to monitor SipA-HiBiT injection into HeLa cells. Figure created with BioRender. **g)** Kinetic of SipA-HiBiT injection intro HeLa cells when the compound or DMSO 1% was added to the bacterial culture. Multiplicity of infection (MOI): 50. Representative replicate from 3 independent experiments. **h)** Effect of C26 at 100 µM on the injection of SipA-HiBiT into HeLa cells when added to the bacterial culture or only to the infection inoculum. Endpoint measurement 1 h post infection. MOI:50. *** p < 0.001; **** p < 0.0001 (Bonferroni’s multiple comparisons test. n = 3 biological replicates).

C26 inhibited the secretion of SipA with an average half maximal inhibitory concentration (IC_50_) of 29.2 µM (Confidence interval (CI): 27.1 - 31.4) (Fig. 1d), and did not impair the growth of *S.* Typhimurium (Extended data Fig. 1). Mammalian cell toxicity was assessed *in vitro* on HeLa cells using the ApoTox-Glo assay. After an exposure of 18 h, the compound exhibited an average half maximal toxic concentration (TC_50 viability_) of 98 µM (Fig. 1e). The toxicity was further assessed in mice at an initial dose of 3 mg/kg in a maximum tolerated dose (MTD) experiment. Neither mortality nor significant adverse effects were observed after oral administration of C26 at 3, 10, and at the highest tested dose of 30 mg/kg in three separate rounds (Supplementary Tab. 2). The body weight gain in all tested animals was normal after 72 h of treatment and no significant abnormalities were observed at termination in all groups (Supplementary Tab. 3).

To better characterize the T3SS-1 inhibitory activity of C26, a split NanoLuc (HiBiT/LgBiT) system was used to quantify the levels of injected SipA into HeLa cells, as previously described^37,38^. SipA was fused to HiBiT, while LgBiT was stably expressed in the cytoplasm of the HeLa cell line. Only if SipA-HiBiT is injected into the HeLa cells, a functional luciferase can be reconstituted by the interaction between LgBiT and HiBiT. Therefore, luminescence intensity inside the host cells can be used as a proxy for the translocation efficiency of SipA-HiBiT. When C26 was added to the bacterial culture (Fig. 1f), then removed from the medium by centrifugation of the inoculum before infecting HeLa cells, we observed a dose-dependent decrease of SipA injection into HeLa cells over a 5 h infection time (Fig. 1g). Similarly, the endpoint measurement of SipA injection at 1 h post infection in the same conditions showed a strong effect of C26 (100 µM) with 11% residual SipA injection (Fig. 1h). Notably, when bacteria were treated exclusively during the infection of host cells (infection inoculum), SipA injection was reduced to an average of 48% of the untreated bacteria (Fig. 1h). This observation suggests a fast inhibition of SipA injection by the compound without the need for prior treatment of growing bacteria.

### C26 impedes host cell invasion by disrupting the expression of invasion genes

We analyzed the expression and secretion of the T3SS-1-secreted proteins SipA, SctE, and SctP by Western blotting (Fig. 2a and 2b). SipA, SctE, and SctP secretion into the culture supernatant was reduced in a dose-dependent manner when bacteria were grown in the presence of C26 (Fig 2a). As expected, no secretion was detected in Δ*invA* and Δ*hilD* mutants, which are deficient in T3SS-1 secretion and expression, respectively. Contrary to the Δ*invA* phenotype, C26 blocked the expression of SipA, SctE, and SctP in whole cells, matching the phenotype of the Δ*hilD* mutant (Fig. 2b). These data suggested that C26 may interfere with the regulation of SPI-1. We therefore performed a transcriptome analysis by RNA-seq on bacteria treated with C26 (100 µM) under SPI-1-inducing conditions (Fig. 2c). C26 led to the downregulation of genes encoded on several SPIs (Fig. 2d), with stronger downregulation observed for genes encoded on SPI-1 and SPI-4 (Fig. 2e and Extended data Fig. 2a). The expression of *hilD*, *hilC*, and *RtsA* was downregulated by a Log_2_ fold-change (FC) of 1.95, 3.02, and 3.82, respectively (Fig. 2e and 2f), and consequently, the expression of *hilA* was also reduced by a Log_2_ FC of 3.94. Based on these results, C26 was assumed to, besides SPI-1, affect the HilA-regulated SPI-4 encoding the T1SS, and to a lesser extent the SPI-2 encoding T3SS-2, which are necessary for invasion and survival inside the host cell, respectively.

**Figure 2.**
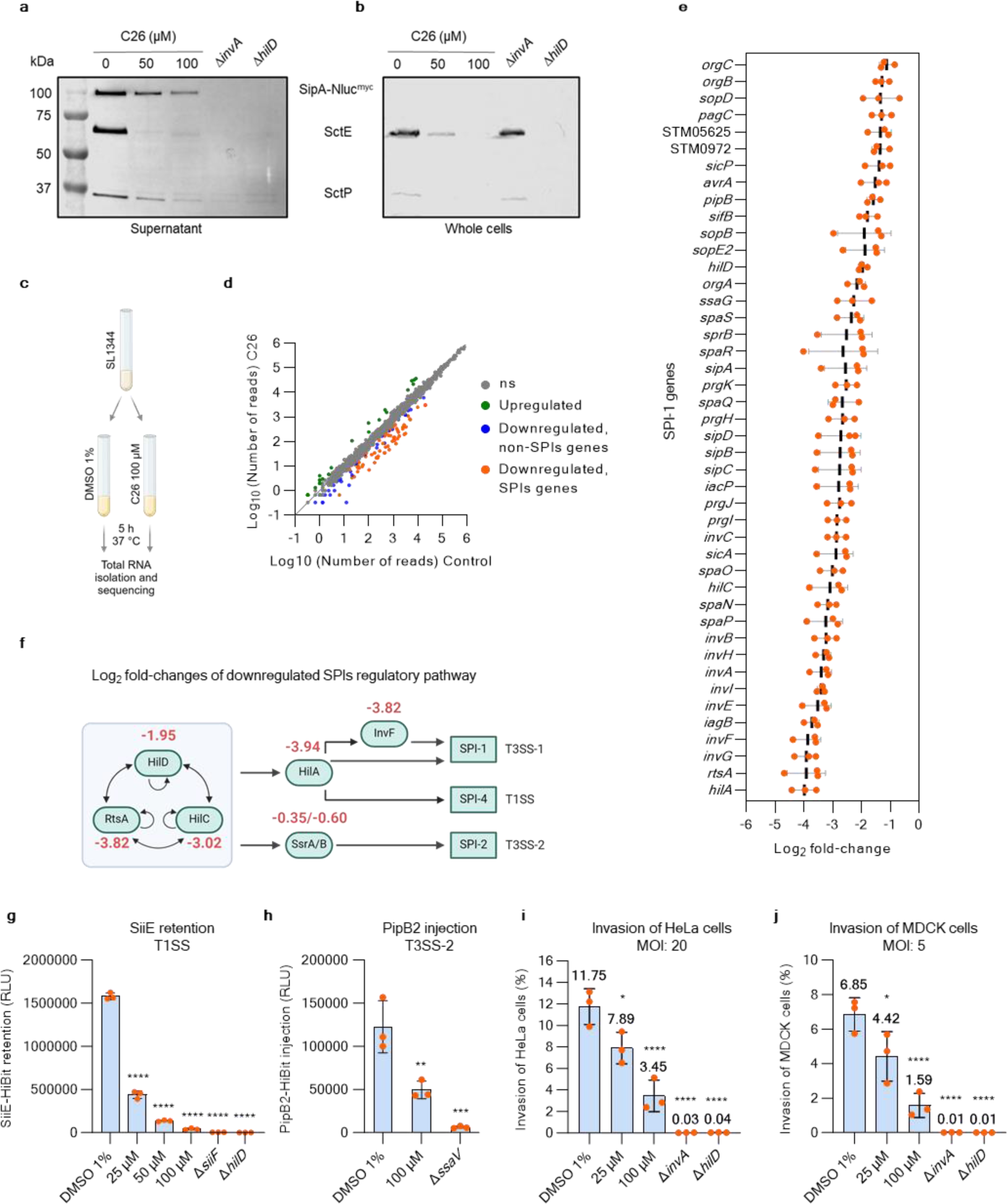
C26 targets the regulatory pathway of SPIs and reduces invasion into host cells. **a, b)** Abundance of T3SS-1 effector proteins in the supernatant **(c)** and in whole cells **(d)** monitored by Western blotting. Mouse anti-*myc* (1:1000), mouse anti-SctE (1:1000), and mouse anti-SctP (1:1000) antibodies were used to quantify SipA, SctE, and SctP, respectively. **c)** Experimental plan of the transcriptome analysis by RNA-seq. Created with BioRender. **d)** Scatter plot representing the level of gene expression when bacteria were grown in the presence of C26 (100 µM) or DMSO (1% (v/v)) as a control condition. ns, not significant. Green: Upregulated genes. Orange: Downregulated genes encoded in SPIs. Blue: Downregulated genes that are not encoded in SPIs. Grey: Below statistical cut-off. **e)** Log_2_ fold-changes in the expression of SPI-1-encoded genes in the presence of C26 (100 µM) (n = 3 biological replicates). **f)** Feed-forward regulatory loop HilD/HilC/RtsA regulating SPIs-encoded genes. The Log_2_ fold-changes in gene expression in the presence of C26 (100 µM) are indicated in red. Figure created with BioRender. **g)** Activity of C26 on the cell surface retention of SiiE-HiBit. Δ*siiF* and Δ*hilD* mutants were used as controls for lack of SiiE secretion and expression, respectively. **** p < 0.0001 (Bonferroni’s multiple comparisons test). n = 3 biological replicates. **h)** Activity of C26 on T3SS-2 as quantified by measuring the injection of PipB2-HiBit inside the host cells. The Δ*ssaV* mutant was used as a control for the lack of T3SS-2 activity. MOI: 100. ** p < 0.01; *** p < 0.001 (Bonferroni’s multiple comparisons test). n = 3 biological replicates. **i, j)** Invasion of HeLa cells, MOI: 20 **(i)** and MDCK cells, MOI: 5 **(j)** by *S.* Typhimurium in the presence of C26. Δ*invA* and Δ*hilD* mutants were used as negative controls. * p < 0.05; **** p < 0.0001 (Bonferroni’s multiple comparisons test). n = 3 biological replicates.

The effect of C26 on the SPI-4 encoded T1SS was quantified using the secretion of the giant adhesin SiiE fused to HiBiT (SiiE-HiBiT) as a readout. The deleted *siiF*, encoding for the ABC-transporter component of the T1SS, and Δ*hilD* mutants served as controls for impeded secretion and expression of SPI-4-encoded genes, respectively. We used the Nano-Glo HiBiT extracellular detection system to quantify the SiiE-HiBiT surface retention. When bacteria were grown in the presence of C26, the amount of SiiE-HiBiT retained on the bacterial cell surface was reduced in a dose-dependent manner (Fig. 2g). Additionally, we showed by Western blotting analysis that C26 led to a dose-dependent reduction of the expression of SiiF (Extended data Fig. 2b), confirming the inhibitory activity of C26 on the SPI-4-encoded T1SS.

The activity of C26 on T3SS-2 was investigated by monitoring the secretion of PipB2-HiBiT by bacteria localized inside the LgBiT-expressing HeLa cells. After allowing bacteria to invade the HeLa cells for 1 h, followed by a gentamycin treatment to kill the non-invading bacteria, the compound was added to the infection medium. Luminescence was measured 16 h after infection and its value corresponds to the amount of PipB2-HiBiT secreted by the bacteria localized inside the host cell. In this experimental setup, 100 µM C26 reduced PipB2-HiBiT secretion to 40% of the control (1% (v/v) DMSO) (Fig. 2h), suggesting that the compound interferes with the activity of the T3SS-2 when *Salmonella* is inside the eukaryotic host cell.

Finally, we investigated how C26 affected the invasion of *S*. Typhimurium into HeLa cells (Fig. 2i) and MDCK cells (Fig. 2j). Under standard conditions, an average of 11.8% and 6.9% of the original inoculum invaded HeLa cells and MDCK cells, respectively. In the presence of 100 µM C26, the counts of *S.* Typhimurium inside HeLa cells and MDCK cells decreased to 3.5% and 1.6% of the original inoculum, respectively. The effect of C26 on invasion, however, did not reach the level of Δ*invA* and Δ*hilD* mutants, for which invasion was abolished. All together, our results provide evidence that C26 hinders the invasion of host cells by targeting the regulation of the gene expression of SPI-1 and SPI-4, and also affects the expression of SPI-2.

### C26 targets the transcriptional regulator HilD

To decipher how C26 downregulates the invasion-associated SPIs, a cell-based assay was developed to monitor the activity of HilD, HilC, and RtsA. The endogenous *hilA* promoter (P*hilA*) was fused with a reporter gene encoding superfolder green fluorescent protein (sfGFP). We first tested the effect of C26 on P*hilA* activation in different knockouts of *hilD*, *hilC*, and *rtsA* (Extended data Fig. 3a). In Δ*hilD* strains, P*hilA* activation was at the background noise. The deletion of *hilC*, *rtsA*, or both, did not affect the activation of P*hilA*, and C26 remained as active as in the WT strain. These results are in accordance with previous observations on the minor role of HilC and RtsA in the activation of P*hilA* in SPI-1-inducing conditions^18^. P*hilA* activation levels can therefore be used as a proxy of HilD activity. Several HilD inhibitors have been described. Among them, the fatty acids (FAs) oleic acid, palmitoleic acid, cis-2-hexadecenoic acid (c2-HDA)^29,32^, and the bile acid chenodeoxycholic acid (CDCA)^35^(Fig. 3a). We used the P*hilA* activation assay to compare their HilD inhibitory activity with that of C26 (Fig. 3b). Here, C26 exhibited an IC_50_ of 16.9 µM (CI: 14.1 - 20.3 µM). The IC_50_ of oleic acid and palmitoleic acid were 25.1 µM and 25.8 µM, respectively. c2-HDA exhibited a strong inhibition of HilD with an IC_50_ of 0.21 µM. No activity of CDCA was observed at the highest tested concentration of 100 µM. These data are in line with the reported activities of the FAs^39^ and CDCA^35^, and therefore confirm the reliability of the assay to quantify HilD transcriptional activity.

**Figure 3.**
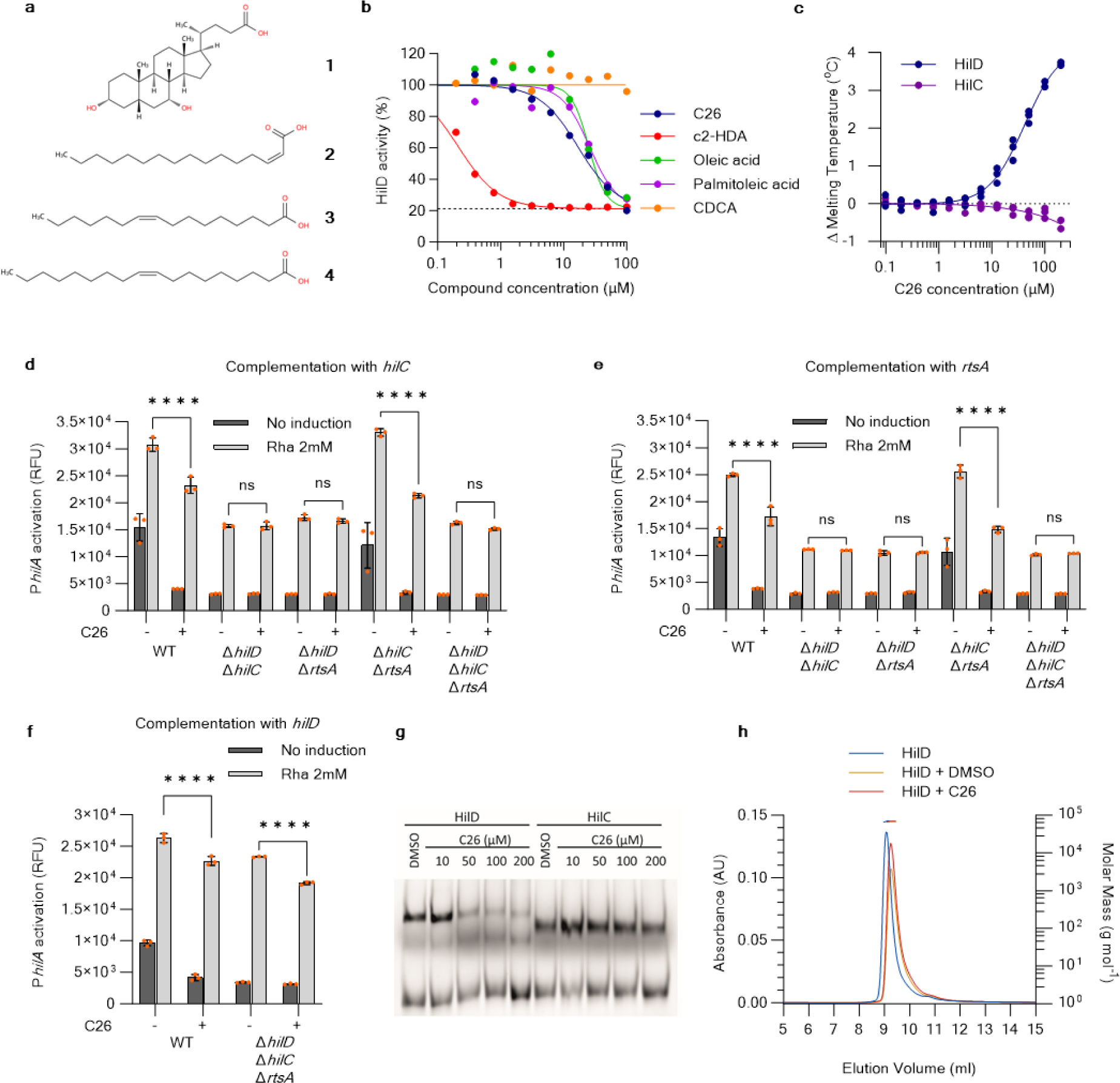
C26 targets the transcriptional regulator HilD. **a)** Structure of the HilD inhibitors chenodeoxycholic acid (**1**), cis-2-hexadecenoic acid (**2**), palmitoleic acid (**3**), and oleic acid (**4**). **b)** Cell-based HilD activity assay. Dose-response curve of P*hilA*-sfGFP expression with increasing concentrations of C26 and other known HilD inhibitors. Fluorescence measured at 485_Ex_/510_Em_ (n = 3 biological replicates). The dashed line corresponds to the baseline Δ*hilD*. **c)** Changes in the calculated melting temperature of HilD and HilC upon incubation with increasing concentrations of C26, as determined from the fluorescence at 350 nm by NanoDSF, (n = 3 separate experiments). **d, e)** Effect of C26 (100 µM) on P*hilA* activation in different background strains complemented with *hilC* **(d)** or *rtsA* **(e)**. ns, not significant; * p < 0.05; **** p < 0.0001 (Bonferroni’s multiple comparisons test) (n = 3 biological replicates). **f)** Effect of C26 (100 µM) on strains overexpressing *hilD*. ns, not significant; * p < 0.05; **** p < 0.0001 (Bonferroni’s multiple comparisons test) (n = 3 biological replicates). **g)** Electrophoretic mobility shift assay (EMSA) showing the effect of C26 on the binding of purified HilD and HilC to the promoter of *hilA*. **h)** SEC-MALS analysis of HilD in presence of DMSO (1%) or C26 (100 µM). Left x-axis shows UV absorbance measured at 280 nm. Right x-axis shows the calculated molecular weight values from light scattering, highlighted by horizontal dashes.

To assess the affinity of C26 to HilD and HilC *in vitro*, we used nano-Differential scanning fluorimetry (nanoDSF), a technique monitoring protein unfolding from intrinsic fluorescence. C26 binding resulted in a dose-dependent increase in the melting temperature of HilD with an apparent K_D_ of 30.2 μM, while no effect on the thermal stability of HilC was observed (Fig. 3c).

Next, we investigated the selectivity of C26 for HilD using the cell-based P*hilA* activation assay. In the absence of HilD, the pool of HilC and RtsA is insufficient to activate the expression of *hilA*. Therefore, assessing the sensitivity of HilC and RtsA to C26 requires P*hilA* to be activated in a HilD-independent manner. To create such a condition, we used a plasmid-encoded *hilC* or *rtsA* under the control of a rhamnose-inducible promoter. Induction of *hilC* (Fig. 3d) or *rtsA* (Fig. 3e) with rhamnose at 2 mM in Δ*hilD* Δ*hilC*, Δ*hilD* Δ*rtsA*, and Δ*hilD* Δ*hilC* Δ*rtsA* knockout mutants resulted in a P*hilA* activation level close to that of the WT strain. Under these conditions, in which either *hilC* and/or *rtsA* are the sole activators of P*hilA*, C26 did not lead to a reduction of P*hilA* activation, as opposed to the WT and the Δ*hilC* Δ*rtsA* backgrounds where P*hilA* activation was reduced in the presence of C26. These data suggested that the compound does not impair the activity of HilC and RtsA.

Supposing that HilD is the target of C26, its overexpression would titrate C26 activity and enable P*hilA* activation, thus resulting in a resistance mechanism by target overexpression. In the absence of C26, inducing the expression of a plasmid-encoded *hilD* resulted in an increase of P*hilA* activation both in the WT background and in the Δ*hilD* Δ*hilC* Δ*rtsA* knock-out strain (Fig. 3f). In the presence of C26, P*hilA* activation decreased significantly by 14% and 18% in WT and Δ*hilD* Δ*hilC* Δ*rtsA* background strains, respectively, indicating an activity against HilD. The remaining high activation level of P*hilA* in the presence of C26 could be explained by the relatively high concentration of induced HilD when compared to the standard pool in the WT strain.

To understand the mechanism of HilD inhibition, we assessed whether C26 impaired the DNA-binding activity of HilD using an electrophoretic mobility shift assay (EMSA). HilC, which does not interact with C26, was used as a negative control. (Fig. 3g). Recombinant HilD and HilC both bind to a fragment of the *hilA* promoter, encompassing the common A1 binding site^15,16^. In the presence of C26, we observed a dose-dependent inhibition of the DNA-binding activity of HilD, while no effect on HilC activity was observed.

CDCA and oleic acid have been shown to inhibit the binding of HilD to DNA by disrupting HilD homodimerization^35,39^. We used multi-angle light scattering coupled to size-exclusion chromatography (SEC-MALS) to investigate whether C26 had a similar effect. However, incubation with equimolar amounts of C26 had no effect on the oligomerization state of HilD, which eluted as a dimer (Fig. 3h and Supplementary Tab. 4). We confirmed this result by performing BS^3^ (bis(sulfosuccinimidyl)suberate) cross-linking of HilD after incubation with C26 or oleic acid (Extended data Fig. 3b). Decreased levels of the cross-linked HilD dimer were observed in the presence of oleic acid at 50 µM and 100 µM, while C26 did not affect the levels of cross-linking at the highest tested concentrations of 200 µM. Taken together, our data suggest that C26 inhibits HilD binding to P*hilA* without disrupting its dimerization, a mechanism distinct from that of other inhibitors of HilD. We further investigated the effect of C26 on the formation of HilD-HilE heterodimers using a microscale thermophoresis (MST) dimerization assay that we previously described^39^. In contrast to oleic acid, which disrupted the binding of HilE to HilD, no effect on heterodimerization was observed for C26 (Extended data Fig. 3c).

### Structural analysis and druggability of HilD

To assess the druggability of HilD, we first performed a bioinformatics analysis. Homologs of *Salmonella* HilD from γ-proteobacteria and a pool of representative homologous sequences, belonging to the AraC/XylS family, were retrieved from NCBI/GenBank using BLAST (accession date 01/08/2022) and from the full draft genomes of the Integrated Microbial Genomes and Microbiomes database (Fig. S4). No similar sequences were found in vertebrate genomes. Phylogenetic analyses of the AraC/XylS family showed that HilC and HilD share the highest similarity (Extended data Fig. 4). The lack of binding of C26 to HilC (Fig. 3c) therefore strengthens the assumption that the compound selectively binds to HilD.

**Figure 4.**
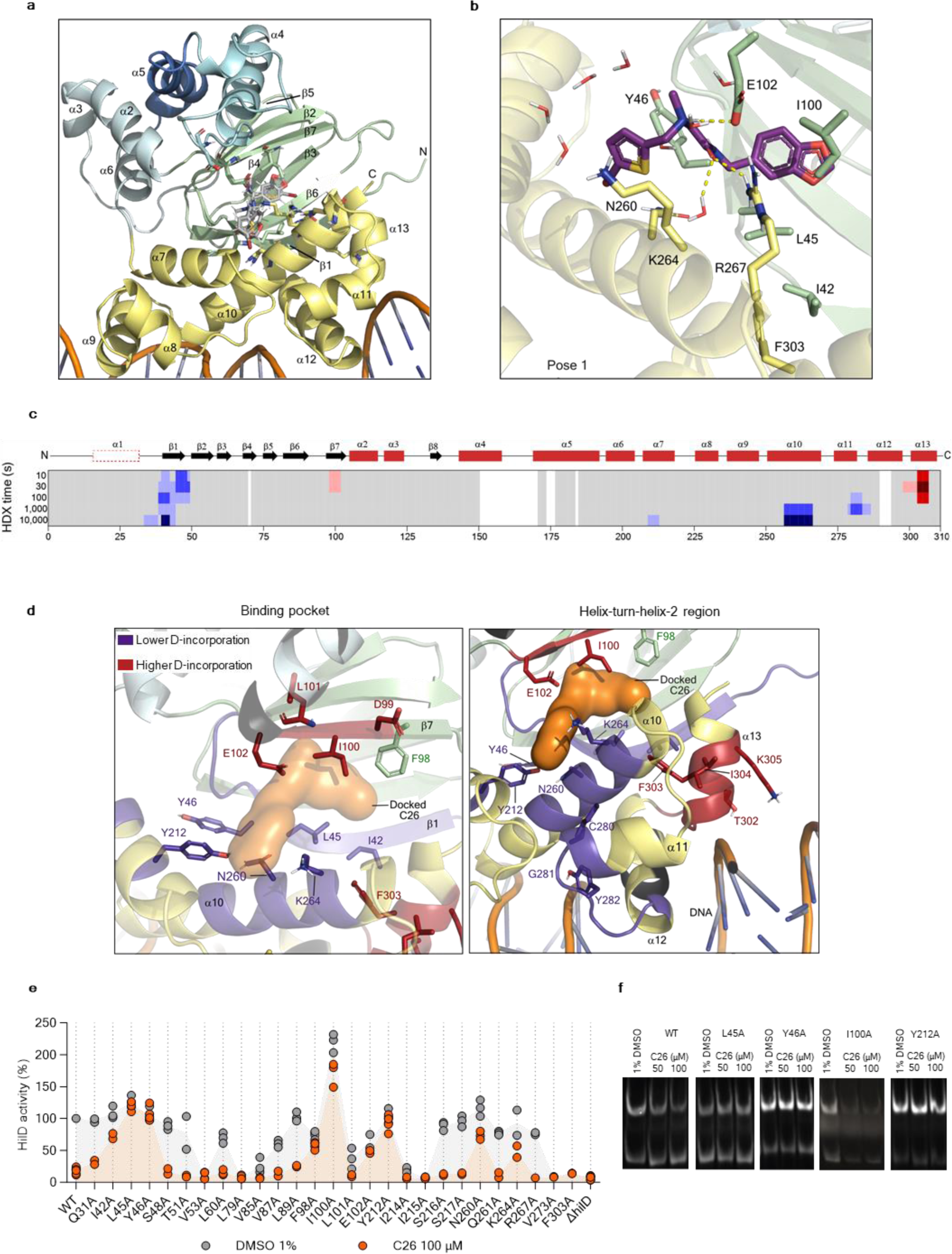
Druggability of HilD and structural characterization of HilD-C26 complex. **a)** HilD model with dsDNA generated by AlphaFold. DNA binding domain is highlighted in yellow, bound to a generic dsDNA fragment, while the beta-sheets of the cupin barrel are depicted in green, dimerization interfaces are displayed in tones of blue and numbered accordingly. **b)** Structural representation of the proposed binding mode from molecular modelling of pose 1 within the predicted binding pocket, generated by clustering the MD trajectory by the ligand RMSD variation (see extended methods). **c)** The difference in HDX between C26-bound and apo HilD projected on its amino acid sequence. Different tones of blue or red reflect, respectively, a decrease or an increase in HDX in presence of C26 (100 µM). The HilD secondary structure is schematically depicted above (red rectangles, α-helices; black arrows, β-strands). **d)** Mapping of the regions exhibiting a lower (blue) or higher (red) deuterium incorporation in presence of C26 (100 µM) as identified by HDX-MS. Zoom on the predicted binding pocket (left), and on the predicted DNA-binding HTH-2 region (right). Relevant residues are shown as sticks. Binding pocket volume is depicted in an orange surface. **e)** Cell-based assay to monitor the sensitivity of HilD mutants to C26. Bacteria were treated with either 1% DMSO (grey dots), or 100 µM C26 (orange dots). HilD activity of the mutants was calculated relative to HilD wild-type (WT) grown in DMSO 1% (n = 3 biological replicates). **f)** EMSAs showing the binding of 600 nM HilD_WT_, HilD_L45A_, HilD_Y46A_, HilD_I100A_, and HilD_Y212A_ to P*hilA*, upon incubation with the indicated concentrations of C26.

To gain a deeper understanding of the interaction between C26 and HilD, we used AlphaFold2 to generate a structural model for the HilD core (residues 37-308), which was simulated with and without a short DNA fragment (see methods). HilD consists of an N-terminal domain with a cupin barrel and an all α-helical dimerization interface, and a C-terminal DNA binding domain with two helix-turn-helix (HTH) motifs (Fig. 4a). We previously identified a pocket involving the cupin barrel and α-helices 7 and 10 as the oleic acid binding site^39^. Assuming the same pocket to bind C26, we identified potential key amino acid positions involved in HilD-ligand interaction using simulations of protonated C26 within this pocket (Fig. 4b). Our docking calculations suggested two different potential binding modes for C26: pose 1 (Fig. 4b and Extended data Fig. 5), with the bromothiophene establishing a chalcone interaction with the backbone of N260 and a cation-π interaction with K264, and pose 2 (Extended data Fig. 5), horizontally inverted with this moiety accommodated within the cupin barrel near residues L45 and I100. We performed longer simulations for both poses to derive their relevant protein-ligand interactions and determine the potential binding energy for each pose. Interestingly, simulations of pose 1 were most stable within HilD’s pocket, as observed by their small variation of the root mean square deviation (RMSD), and low predicted binding energy values (Extended data Fig. 6 and Fig. 7), which together would support this as the preferred binding mode. We could further gain experimental support for this conclusion by NMR spectroscopy via saturation transfer difference (STD) experiments on the HilD-C26 complex. Intensities in STD experiments are sensitive to the proximity of the ligand to the saturated groups in the protein, here the methyl groups of aliphatic residues. The high concentration of these groups surrounding the cupin barrel binding pocket of HilD allowed the discrimination of the binding poses. We applied the CORCEMA algorithm^40–42^ to predict STD intensities for frames of both MD trajectories and calculate an R-factor for each frame reflecting the fit to the experimental data. The trajectory starting from pose 1 has a significantly higher density of frames with lower R-factors than that starting from pose 2 (Extended data Fig. 8).

**Figure 5.**
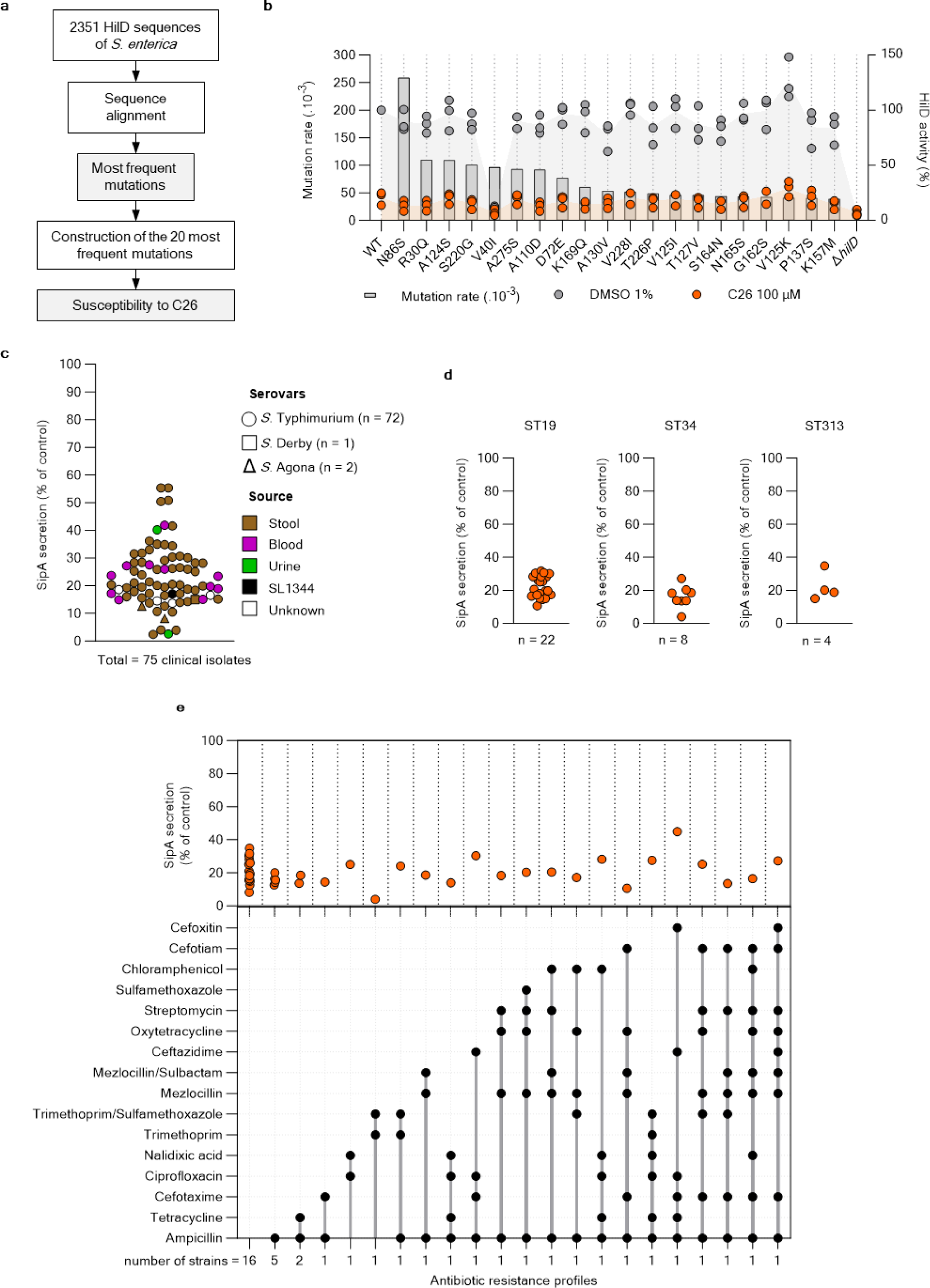
Spectrum of activity. **a)** Applied workflow for the identification of the most frequent amino acid substitutions in HilD among *S. enterica*. **b)** Mutation rates of the 20 most frequent substitutions in 2,351 sequences of HilD, and their consequence on sensitivity to C26 (100 µM). Mutation rates are shown in grey bars (left y-axis). HilD activity (right y-axis) was quantified as in Fig. 4e. Bacteria were treated with either 1% DMSO (black dots), or 100 µM C26 (orange dots). **c)** Activity of C26 (100 µM) on clinical strains of *S. enterica* isolated from human stool (brown), blood (purple), and urine (green) samples. 100% corresponds to SipA secretion in bacteria treated with 1% DMSO. **d)** Activity of C26 (100 µM) on clinical strains of *S. enterica* isolates clustered by sequence type (ST). 100% corresponds to SipA secretion in bacteria treated with 1% DMSO. **e)** Combination matrix (bottom) of the phenotypic antibiotic resistance profiles of *S*. Typhimurium clinical isolates (total = 42), and their corresponding sensitivity to C26 (100 µM) as monitored by quantification of SipA secretion (top). 100% corresponds to SipA secretion in bacteria treated with 1% DMSO. Each orange dot corresponds to a single clinical isolate.

**Figure 6.**
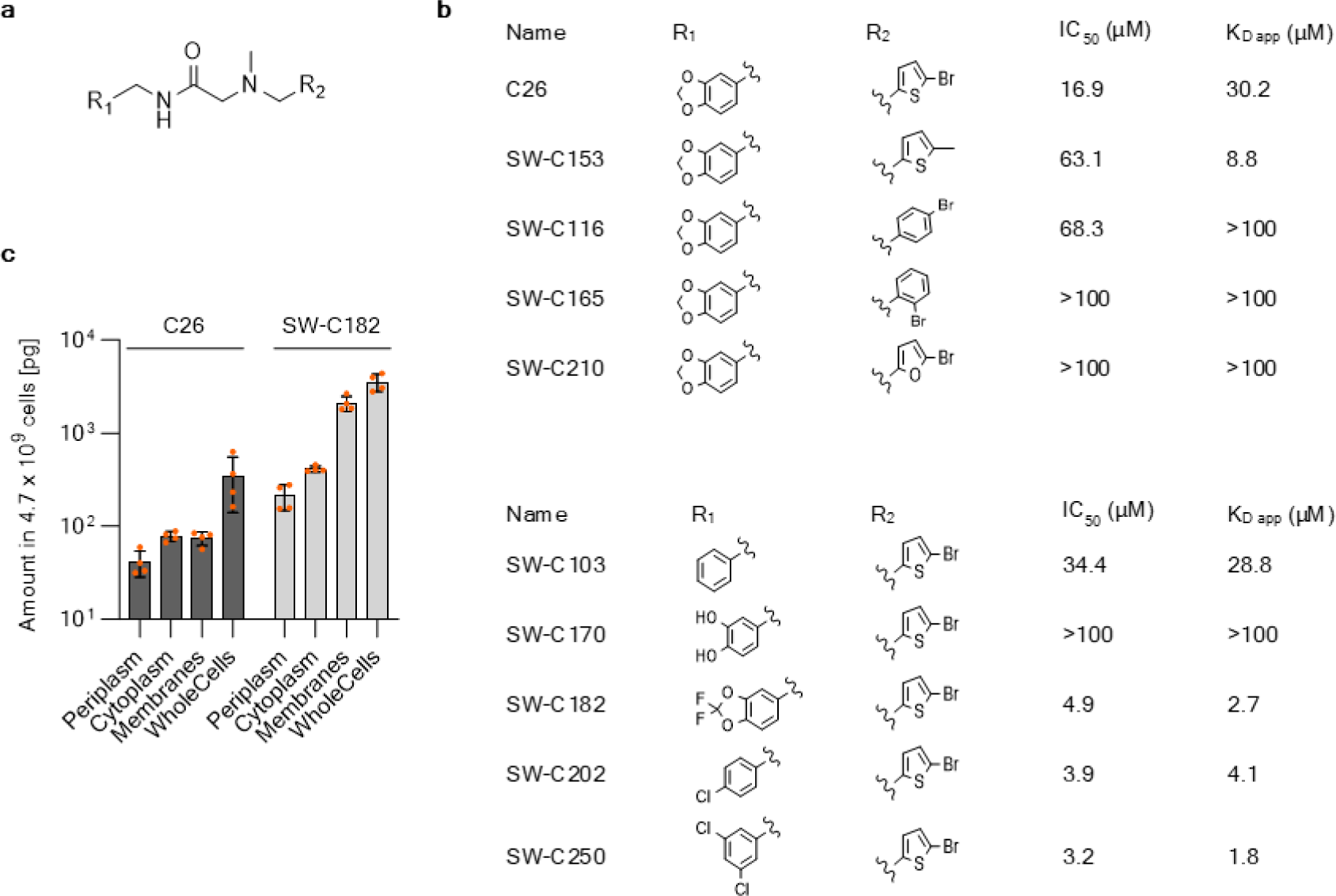
Structure-activity relationship analysis. **a** and **b)** Structures and activities of C26 and analogs. **c)** Quantification of C26 and SW-C182 in subcellular compartments of *S*. Typhimurium. Whole cell is the amount found in unfractionated bacteria (n = 3 biological replicates).

### Structural characterization of the HilD-C26 complex

To probe this binding model further, we performed hydrogen-deuterium exchange mass spectrometry (HDX-MS) on purified HilD alone (with DMSO as a mock) or in the presence of C26 (100 µM). Differences in deuterium exchange upon incubation with C26 are highlighted along the sequence of HilD (Fig. 4c and Extended data Fig. 9) and mapped onto the HilD model to highlight the binding pocket and the helix-turn-helix-2 region (α11 and α12) (Fig. 4d). Decreased HDX was observed for residues 35-50, 209-212, and 256-265, all of which are located in the predicted binding pocket in agreement with our computational model (Fig. 4b). These HDX changes mirror those observed for oleic acid^39^, supporting the assumption that both compounds bind to the same binding pocket. In contrast to oleic acid binding, areas of decreased HDX induced by C26 in the DNA-binding domain of HilD were restricted to a short stretch (residues 279-285, α11). Interestingly, an HDX increase was apparent for helix α13 (residues 297-305) in the presence of C26, while oleic acid exclusively reduced HDX in helices α11-13 and the interconnecting linkers^39^. It is likely that binding of C26 to the pocket enclosed by α10-α7-β1-β7 causes a conformational change in helices α11 and α13, resulting in the loss of affinity to P*hilA*. We propose this mechanism to be the mode of action of C26 rather than interference with HilD dimerization as shown for oleic acid (Fig. 3h and Extended data Fig. 3c).

Next, we developed a plasmid-based system for the fast introduction of point mutations to assess their effect on HilD activity and sensitivity to C26. An alanine scan was performed on amino acid positions located in the predicted binding pocket. F303A inactivated HilD, which confirms the importance of the helix α13 for DNA binding, and therefore supports the proposed mode of action of C26. L45A, Y46A, E102A, and Y212A resulted in a full loss of sensitivity to C26 (100 µM), while I42A, F98A, I100A, N260A, and K264A resulted in a partial loss of sensitivity (Fig. 4e). We then assessed the effect of C26 on the DNA binding of HilD_L45A_, HilD_Y46A_, HilD_I100A_, and HilD_Y212A_ by EMSA, and observed that each of the mutants bound to the *hilA* promoter with comparable affinity to HilD_WT_ (Supplementary Fig. 1). C26 did not exert any effect on the DNA-binding ability of HilD_L45A_, HilD_Y46A_, and HilD_Y212A_ (Fig. 4f), confirming the resistance of these mutants to C26. HilD_I100A_ binding to P*hilA* was hindered by the compound, suggesting that its reduced sensitivity to C26 in the cell-based assay (Fig. 4e) may be attributed to its 2-fold higher transcriptional regulatory activity as compared to HilD_WT_. Combining data generated by MD simulations, biophysical methods, and cell-based and *in vitro* assays, we were able to map the binding pocket, identify the amino acid residues important for C26 action, and confirm the proposed mode of action of the compound.

### Spectrum of activity

Having identified amino acid substitutions leading to resistance of HilD to C26, we searched for the presence of these mutations among 2351 HilD sequences from NCBI (acquired on 06/10/2023) (Fig. 5a). A sequence alignment was first performed to identify the most frequent substitutions as compared to the reference HilD sequence of SL1344 (Supplementary Tab. 5). Using the plasmid-based system to monitor HilD activity, we individually introduced the 20 most frequent substitutions into the sequence of HilD and quantified their sensitivity to C26 (Fig. 5b). Except for V40I, which resulted in a non-functional HilD, none of the other 19 substitutions affected HilD transcriptional activity. Importantly, C26 exhibited an inhibitory activity on all the tested variants at a level similar to that of the WT reference sequence.

Finally, we aimed to evaluate the spectrum of activity of C26 among clinical isolates of *S. enterica*. We acquired 37 strains of *S.* Typhimurium isolated from patients between 2010 and 2020 at the university hospital of Tübingen, Germany. An additional set of 70 representative clinical *S. enterica* strains, covering different sequence types (Supplementary Fig. 2), and antibiotic resistance profiles (Supplementary Tab. 6), were selected from the strain collection of the National Reference Centre for *Salmonella* at the Robert Koch Institute, Germany (total = 107 clinical isolates). We used a plasmid-encoded *sipA*-NLuc to assess the secretion levels of SipA through T3SS-1 in the 107 clinical isolates. 32 strains exhibited SipA secretion levels lower than the predefined cut-off of 5% of the reference strain SL1344 and were therefore excluded from further analyses (Extended data Fig. 10). The remaining 75 strains, regardless of their isolation source (stool: n = 58, blood: n = 11, or urine: n = 2), were all sensitive to C26 (100 µM) (Fig. 5c). Notably, the compound exhibited an activity against *S.* Agona (n = 2) and *S.* Derby (n = 1). We then clustered the characterized strains according to their clinical multilocus sequence types (MLST). Strains belonging to most frequent STs in Germany ST19 (n = 22) and ST34 (n = 8), and other STs such as ST313 (n = 4), were all sensitive to C26 without a reduced sensitivity pattern (Fig. 5d). Finally, we clustered the *S*. Typhimurium clinical isolates according to their phenotypic antibiotic resistance profiles (Fig. 5e). The latter had no influence on the inhibitory activity of the compound, strengthening the assumption that anti-virulence agents are characterized by a reduced risk of cross-resistance with direct-acting antibiotics. We conclude that C26 has an advantageous activity spectrum among *S.* Typhimurium clinical isolates, regardless of their source of isolation, sequence type, and antibiotic resistance profile.

### Structure-activity relationship analysis

To gauge the potential for further optimization of C26 potency, we performed an initial structure-activity relationship (SAR) analysis. The 5-bromothiophen-2-yl unit turned out to be favorable, since its replacement by isosteres such as 2-methylthiophene (SW-C153), 4-bromophenyl (SW-C116), 2-bromophenyl (SW-C165), and 2-bromofurane (SW-C210) led to less active compounds in the P*hilA* activation assay with IC_50_’s of 63.1, 68.3, >100, and >100 µM, respectively (Fig. 6a and 6b). The replacement of the benzodioxol moiety allowed a fine-tuning of activities. Compound SW-C103 with a simple phenyl residue, was slightly less potent (IC_50_ = 34.4 µM), while the removal of the methylene bridge in the heterocycle to give the catechol (SW-C170) led to a complete loss of activity. Importantly, the 4-chloro- and 3,5-dichlorophenyl compounds SW-C202 and SW-C250 were considerably more potent than C26, with IC_50_’s of 3.9 and 3.2 µM, respectively. This improved potency was due to an increased affinity of SW-C202 and SW-C250 to HilD, as reflected by apparent K_D_ values obtained by nanoDSF of 4.1 µM and 1.8 µM, respectively.

In order to probe whether cellular uptake also contributes to differences in cellular bioactivities, we adapted our method for the quantification of intracellular uptake to *S*. Typhimurium^43^. The overall uptake of compound SW-C182 was 10-fold higher than that of C26 (Fig. 6c and Supplementary Tab. 7). In the cytosol, the target compartment of the inhibitors, uptake differed by 5.3-fold. This implies that the activity of C26 and its analogs are driven by both cellular uptake as well as binding to HilD. In summary, a first round of hit-to-lead led to a 10-fold improved, low µM cellular activity, demonstrating the optimization potential of the C26 series.

## Discussion

The slow pace at which direct-acting antibiotics targeting Gram-negative pathogens are being discovered and developed requires the exploration of different approaches. Instead of selecting drug targets that are essential for bacterial survival, we pursued an anti-virulence strategy, targeting the invasion-mediating pathogenicity factors of *S. enterica*. We designed a combined *in silico* and phenotypic screening assay to identify T3SS-1 inhibitors and discovered a small-molecule targeting the transcriptional regulator HilD. Our hit compound (C26) is characterized by a straightforward chemical synthesis, and a favorable, rule-of-five compliant druglikeness. By combining molecular dynamics simulations with experimental evidence, we showed that C26 binds to the HilD homodimer to inhibit its binding to P*hilA*, identified the binding pocket, and suggested a mechanism of HilD inhibition.

Several natural compounds have been identified as HilD inhibitors. Plant-derived compounds like the cyclic diarylheptanoid myricanol^31^, and the flavonoid fisetin^36^ bind to HilD and inhibit its DNA binding activity. The bile acid CDCA has been shown to interfere with HilD dimerization and DNA binding^35^. Similarly, long chain fatty acids (LCFAs) have been shown to bind to HilD and block its dimerization^32,39^. We showed that C26 shares the same binding pocket as CDCA and LCFAs, however, its binding mode is suggested to be different since the inhibition occurs without interfering with the dimerization of the protein.

LCFAs are characterized by a broad spectrum of targets among the AraC-like regulators including HilC and RtsA^32,39,44,45^, and ToxT and Rns to modulate the virulence of *Vibrio cholerae*^46^ and *Escherichia coli*^47^, respectively. In contrast, our hit compound is, to our knowledge, the first described synthetic molecule to selectively bind and inhibit HilD activity. From a drug development perspective, the latter property is advantageous considering the hurdles associated with polypharmacology^48^.

Point mutations resulting in resistance to C26 were not found among the 20 most frequent HilD variants that were revealed to be fully sensitive to the compound. Additionally, C26 exhibited a good spectrum of activity among *S. enterica* clinical isolates, including strains from ST19, which is associated with gastroenteritis^49^, and ST313, which is the dominant sequence types in sub-Saharan Africa causing systemic infections^50,51^.

The current hit compound shows promising activity and drug-like properties. Our results further demonstrate that C26 is able to engage with its target, HilD, in *Salmonella* within infected host cells. Thus, it can readily cross four biological membranes, which is a critical prerequisite for anti-infective drugs targeting Gram-negative intracellular pathogens. Using this compound as a scaffold to perform structure-activity relationship analysis will be necessary to identify more potent analogs that can serve as drug candidates. Synthetic small molecules targeting HilD could be valuable options for the treatment of *S. enterica* gastrointestinal infections and the prevention of invasive *S. enterica* infections in humans. Considering that secretion systems are essential for the systemic dissemination of NTS in chicken^24^, it is also conceivable to use HilD inhibitors to prevent invasive *S. enterica* infections in poultry.

## Methods

### Strains and growth conditions

All the strains used in this study are listed in Supplementary Tab. 8. Except clinical isolates (Supplementary Tab. 6), all *Salmonella* strains were derived from *Salmonella enterica* serovar Typhimurium SL1344^52^. Plasmids used in this study are listed in Supplementary Tab. 9. Primers used for cloning are listed in Supplementary Tab. 10. *S*. Typhimurium strains were cultured with low aeration at 37 °C in Luria Bertani (LB) medium supplemented with 0.3 M NaCl and the appropriate antibiotic when required.

### Virtual screening of T3SS-1 inhibitors

Virtual screening against T3SS was performed based on the InvA (SctV) C-terminal structure against a commercially available library of ligands. The InvA C-terminal domain as a dimer is available with an excellent resolution^53^ (PDB ID 2X49, resolution 1.50 Å). Potential binding sites were determined using SiteMap, which predicted four potential druggable pockets (DrugScore > 1.0). We proceeded with site 2 (DScore: 1.014), encompassing the region of F388, M505, K512, R544, and M546, which is near the dimerization interface. Ligands were docked within a grid around 12 Å from the centroid of the predicted binding site pocket, as mentioned above.

For this virtual screening step, system preparation and docking calculations were performed using the Schrödinger Drug Discovery suite for molecular modelling (version 2014.1) with standard settings. All ligands were retrieved from Enamine Advanced screening collection (accessed on December 2014 containing 468,436 unique compounds – which is limited by chemical properties: MW≤350 Da, cLogP≤3, and rotB≤7) prepared using LigPrep^54^ to generate the 3D conformation, adjust the protonation state to physiological pH (7.4), and calculate the partial atomic charges with the OPLS2005 force field^55^, generating a total of 1,604,573 states. Docking studies with the prepared ligands were performed using Glide (Glide V7.7)^56,57^ with the Virtual Screening Workflow pipeline that starts docking the total ligand library with high throughput screening (HTVS) precision and just proceeds with the top 10% of the best scored ligands for Single Precision (SP) and, then their top 10% to extra precision (XP). The final 1,000 ligands underwent MM/GBSA calculations to predict their binding energy. Ligands for testing were selected based on their predicted binding energy and visually inspected for hydrogen bond interactions.

### Sequence similarity search and phylogenetic tree

*Salmonella’s* HilD homologues and a pool of representative homologues sequences containing the AraC domain were retrieved from gamma-proteobacteria species. Sequences were retrieved from NCBI/GenBank using the Blast tool (with scoring matrix BLOSUM45 for distant similar sequences) and from the full draft genomes of the Integrated Microbial Genomes and Microbiomes database^58^ (with an e-value cut-off of 10^−5^) creating a dataset. No similar sequences were found in vertebrate genomes. Sequences renaming and editing were performed with in-house Perl scripts. Sequences with less than 30% global similarity were excluded from further analyses. The full dataset was clustered by similarity (99%) using CD-Hit^59^ and a set of representative sequences were selected for global alignment using Muscle^60^. Maximum likelihood phylogenetic tree was generated using PhyML 3.0^61^, with posterior probability values (aBayes) as branch statistical support. The substitution model VT was selected for calculations, by ProtTest3^62^, based on the highest Bayesian Information Criterion values. All other parameters, with the exception of the equilibrium frequencies, were estimated from the dataset. Dendrogram figures were generated using FigTree v1.4.4 (https://github.com/rambaut/figtree/releases).

### Phenotypic screen of T3SS-1 inhibitors

For compound screening, 50 µl of overnight cultures of *Salmonella* with an approximate OD_600_ of 2 were diluted in fresh LB medium to an OD_600_ of 0.05 and added to 384-well plates (Nunc MaxiSorp, white) containing 5 nmol screening compound per well, for a final compound concentration of 100 µM. The plates were incubated for 5 h at 37 °C with shaking at 180 rpm, upon which the bacteria were removed. The plates were washed with phosphate buffer saline (PBS: 137 mM NaCl, 2.7 mM KCl, 10 mM Na_2_HPO_4_, and 1.8 mM KH_2_PO_4_, pH 7.4) using a Tecan HydroSpeed plate washer. Residual PBS after washing was removed, and 50 µl NanoLuc working solution (NanoGlo, Promega) was added to the wells. Luminescence was then measured using a Tecan Sparc Multimode reader. Cultures with *sipA-*NLuc and Δ*invA sipA-*NLuc strains with 1% (v/v) DMSO were used as positive and negative controls, respectively.

### Chemical synthesis

#### Starting Materials

Starting materials were purchased from commercial suppliers (Sigma-Aldrich, TCI, BLDpharm, abcr, Carbolution, Thermo Scientific, Alfa Aesar, Acros Organics) and used without further purification. The compounds SW-C153, SW-C116 and SW-C103 are commercially available and were purchased from commercial suppliers.

#### Accurate Mass method

High resolution masses were obtained using a Maxis II TM HD mass spectrometer (Bruker Daltonics, Bremen, Germany).

#### Flash Column Chromatography

Purification on reverse phase was done with a Pure C-850 FlashPrep system (Büchi) using FlashPure EcoFlex C18 cartridges (Büchi). A gradient of water and acetonitrile was used as an eluent. Dryloads were prepared with silica gel C18, 0.035-0.07, 400-220 mesh (Carl Roth).

Normal phase purification was carried out with a Pure C-810 Flash system (Büchi) using FlashPure cartridges (Büchi). A gradient of cyclohexane and ethyl acetate or dichloromethane and methanol was used as an eluent. Dryloads were prepared with silica gel, 60 Å, 230-400 mesh, 40-63 µm (Merck).

Conventional column chromatography was carried out with silica gel, 60 Å, 230-400 mesh, 40-63 µm (Merck) using the eluents described in the synthesis procedures.

#### High-Performance Liquid Chromatography (HPLC)

HPLC was carried out with a Dionex UltiMate 3000 system (Thermo Scientific) using a Luna® 5 µm C18(2) 100 Å, LC column 250 × 21.2 mm, AXIA^TM^ packed (phenomenex). As an eluent, water and acetonitrile were used without or with 0.1% formic acid.

NMR spectra of synthesized compounds are provided in Supplementary information.

### Synthesis of compound C26

The synthesis of the compound C26 and some of its analogs was performed according to the following scheme 1.

**Scheme 1.**
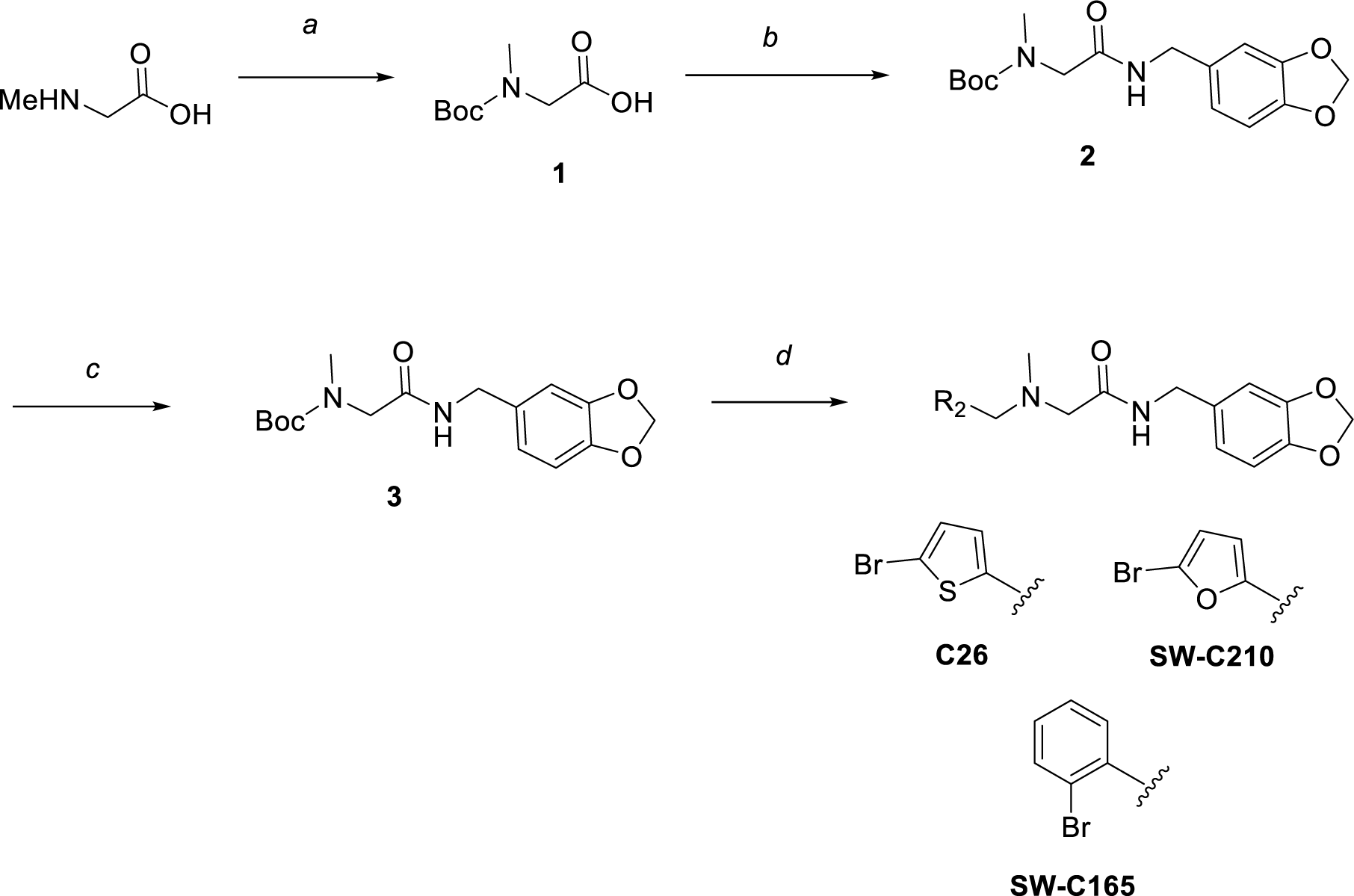
Reagents and conditions. (*a*) Boc_2_O, KOH, H_2_O/dioxane, r. t., overnight 57%; (*b*) iBuOCOCl, Et_3_N, THF, 0 °C to r. t., 2 h, *then* piperonylamine, r. t., 2 h, 84%; (*c*) TFA, CH_2_Cl_2_, r. t., overnight, 72%; (*d*) R_2_CHO, HOAc, THF, r. t., 10 min, *then* NaBH(OAc)_3_, r. t., overnight.

#### *N*-(*tert*-Butoxycarbonyl)-*N*-methylglycine (1)

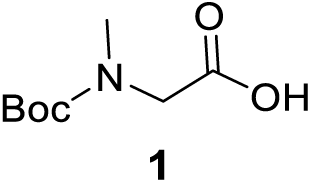

Sarcosine (1.00 g, 11.22 mmol, 1 eq.) was dissolved in a mixture of H_2_O (11 mL) and 1,4-dioxane (25 mL) and the solution was cooled to 0 °C. KOH (2.73 g, 50.51 mmol, 4.5 eq.), dissolved in H_2_O (4 mL), and di-*tert*-butyl dicarbonate (2.94 g, 13.46 mmol, 1.2 eq.) was added and the solution was stirred overnight at r. t.. Then 1,4-dioxane was rotary evaporated and the aqueous residue was acidified with 1 N HCl to pH = 3. The solution was extracted with EtOAc (3 × 40 mL) and combined organic phased were washed with brine, dried over sodium sulfate, filtered and concentrated under reduced pressure to obtain compound **1** as a brownish oil (1.2 g, 6.33 mmol, 57%). The crude product was used for the next step without further purification. The experimental NMR data correspond with those from the literature.

**^1^H NMR** (500 MHz, CDCl_3_): δ 4.02 (s, 1H), 3.95 (s, 1H), 2.94 (s, 3H), 1.46 (d, *J* = 18.7 Hz, 9H).

**LCMS (ESI):** m/z 190 (M + H^+^).

#### *tert*-Butyl (2-((Benzo[d][1,3]dioxol-5-ylmethyl)amino)-2-oxoethyl)(methyl)carbamate (2)

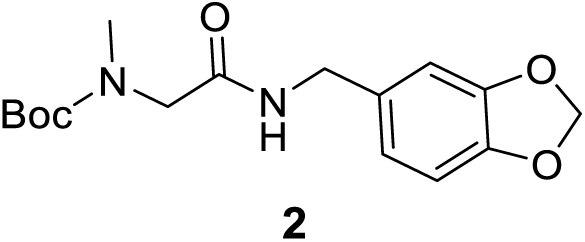

To an ice-cooled and stirring solution of *N*-Boc-sarcosine **1** (1.20 g, 6.33 mmol, 1 eq.) in dry THF (10 mL) was added triethylamine (2.54 mL, 1.36 g, 18.99 mmol, 3 eq.) and isobutyl chloroformate (0.95 g, 6.96 mmol, 1.2 eq). The solution was allowed to come to r.t. and was stirred for 2 h at r.t. under nitrogen atmosphere. Piperonylamine (0.76 mL, 0.95 g, 6.33 mmol, 1 eq.) was added and the reaction mixture was stirred for 2 h under the same conditions. The solution was quenched with an aqueous, saturated NaHCO_3_ solution (50 mL) and extracted with CH_2_Cl_2_ (3 × 40 mL). Combined organic phases were washed with brine (40 mL), dried over sodium sulfate, filtered and concentrated under reduced pressure. The crude product was purified by column chromatography (2 % MeOH in CH_2_Cl_2_) to afford compound **2** (1.71 g, 5.32 mmol, 84%).

**TLC:** R_f_ = 0.38 (CH_2_Cl_2_/ MeOH 50:1).

**^1^H NMR** (500 MHz, CDCl_3_) δ 6.76 – 6.74 (m, 2H), 6.72 (dd, *J* = 7.9, 1.6 Hz, 1H), 5.94 (s, 2H), 4.37 (d, *J* = 5.7 Hz, 2H), 3.88 (s, 2H), 2.94 (s, 3H), 1.43 (s, 9H).

**LCMS (ESI)**: m/z 345 (M + Na).

#### *N*-(Benzo[d][1,3]dioxol-5-ylmethyl)-2-(methylamino)acetamide (3)

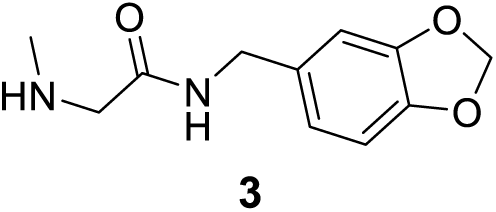

Compound **2** (629 mg, 1.95 mmol, 1 eq) was dissolved in CH_2_Cl_2_ (10 mL). TFA (5 mL) was added and the reaction mixture was stirred overnight at r.t. The solution was quenched with an aqueous, saturated NaHCO_3_ solution (50 mL) and extracted with CH_2_Cl_2_ (3 × 40 mL). Combined organic phases were washed with brine (40 mL), dried over sodium sulfate, filtered and concentrated under reduced pressure to afford the compound **3** (311 mg, 1.40 mmol, 72%). The crude product was used for the next step without further purification.

**^1^H NMR** (500 MHz, CDCl_3_): δ 6.78 – 6.75 (m, 3H), 5.94 (s, 2H), 4.38 (d, *J* = 5.9 Hz, 2H), 3.93 (d, *J* = 6.7 Hz, 2H), 2.46 (s, 3H).

**LCMS (ESI):** m/z 223 (M + H^+^).

#### *N*-(benzo[d][1,3]dioxol-5-ylmethyl)-2-(((5-bromothiophen-2-yl)methyl)(methyl)amino)acetamide (C26)

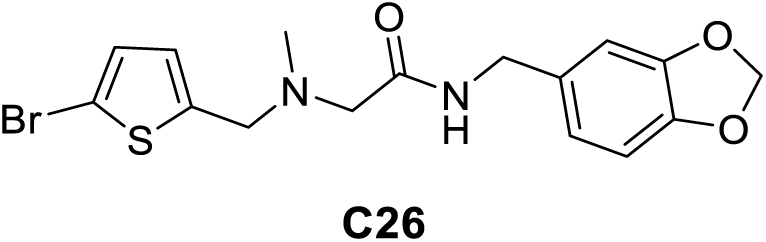

To a stirring solution of compound **3** (202 mg, 0.90 mmol, 1 eq) in dry THF (6 mL) was added 5-bromothiophene-2-carbaldehyde (83 µL, 146 mg, 0.76 mmol, 0.84 eq.) and acetic acid (104 µL, 109 mg, 1.80 µmol, 2 eq.). After 10 minutes, sodium triacetoxyborohydride (288 mg, 1.35 mmol, 1.5 eq) was added and the solution was stirred for 18 h at r. t. under nitrogen atmosphere. The reaction mixture was concentrated under reduced pressure, the crude product was purified by RP flash chromatography, and product-containing fractions were lyophilized. The residue was taken up in MeOH and purified by HPLC to afford compound **C26** (58 mg, 0.14 µmol, 20%).

**^1^H NMR** (500 MHz, DMSO-d_6_) δ 8.19 (t, *J* = 5.8 Hz, 1H), 7.06 (d, *J* = 3.7 Hz, 1H), 6.86 – 6.83 (m, 3H), 6.73 (dd, *J* = 8.1, 1.4 Hz, 1H), 5.97 (s, 2H), 4.20 (d, *J* = 6.1 Hz, 2H), 3.79 (s, 2H), 3.05 (s, 2H), 2.25 (s, 3H).

**^13^C NMR** (126 MHz, DMSO-d_6_) δ 169.1, 147.23, 147.21, 146.0, 133.4, 129.8, 127.0, 120.4, 110.5, 107.9, 100.8, 59.1, 55.5, 42.0, 41.6.

**HRMS (ESI)** calculated for C_16_H_18_BrN_2_O_3_S (M (^79^ Br) + H^+^): 397.0222, found: 397.0214; calculated for C_16_H_17_BrN_2_O_3_S (M (^81^ Br) + H^+^): 399.0201, found: 399.0194.

#### N-(benzo[d][1,3]dioxol-5-ylmethyl)-2-((2-bromobenzyl)(methyl)amino)acetamide (SW-C165)

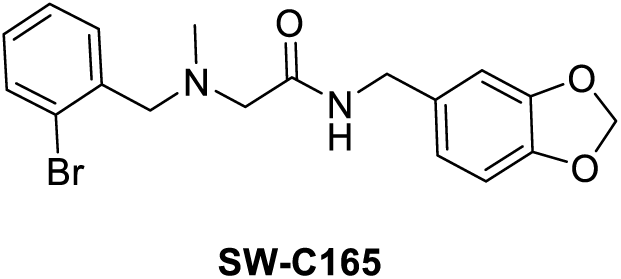

To the compound **3** (35 mg, 0.157 mmol, 1.0 eq.) in THF (2 mL) was added 2-bromobenzaldehyde (29 mg, 0.157 mmol, 1.0 eq.) and acetic acid (19 µL, 0.314 mmol, 2.0 eq.) at r. t. under nitrogen. The mixture was stirred for 5 min and sodium triacetoxyborohydride (50 mg, 0.236 mmol, 1.5 eq.) was added in one portion and stirred overnight at r. t. The reaction mixture was quenched with saturated NaHCO_3_ solution (30 mL) and extracted with Et_2_O (2 × 25 mL). The combined organic layers were dried over Na_2_SO_4_, filtered and concentrated under reduced pressure. The crude compound was purified by column chromatography (PE: EtOAc, 2:1) to obtain the compound **SW-C165** (37 mg, 61%) as colorless oil.

**TLC analysis:** 1:1, EtOAc: PE *R*_f_ : 0.5 (Stain: UV/KMnO_4_).

**^1^H NMR** (400 MHz, CDCl_3_): δ = 7.56 (bs, 1H), 7.53 (d, *J* = 8.4 Hz, 1H), 7.22-7.27 (m, 2H), 7.10-7.18 (m, 1H), 6.73 (d, *J* = 7.9 Hz, 1H), 6.64-6.70 (m, 2H), 5.94 (s, 2H), 4.30 (d, *J* = 6.1 Hz, 2H), 3.65 (s, 2H), 3.12 (s, 2H), 2.31 (s, 3H).

**^13^C NMR** (101 MHz, CDCl_3_): δ = 170.7, 148.0, 147.0, 137.2, 133.4, 132.3, 131.6, 129.4, 127.5, 125.2, 121.0, 108.4, 108.4, 101.1, 62.3, 60.5, 43.5, 42.9.

**HRMS (ESI)** calculated for C_18_H_20_BrN_2_O_3_ (M + H^+^): 391.0657, found: 391.0656.

#### N-(benzo[d][1,3]dioxol-5-ylmethyl)-2-(((5-bromofuran-2-yl)methyl)(methyl)amino)acetamide (SW-C210)

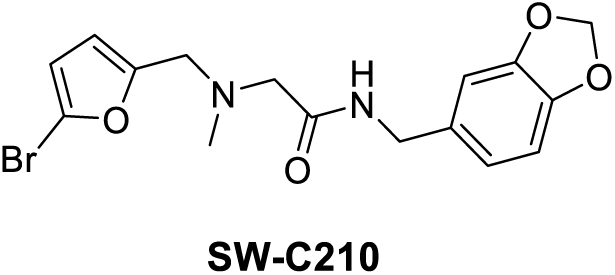

The compound **3** (151.6 mg, 0.68 mmol, 1 eq) was dissolved in THF (10 mL). 5-Bromo-2-furaldehyde (119 mg, 0.68 mmol, 1 eq.), acetic acid (78 µL, 80 mg, 1.36 mmol, 2 eq.) and sodium triacetoxyborohydride (216 mg, 1.02 mmol, 1.5 eq.) were added and the yellowish reaction mixture was stirred for 16 h at r. t. under nitrogen atmosphere. The reaction was quenched with saturated aqueous NaHCO_3_ solution (30 mL) and the aqueous phase was extracted with diethyl ether (3 × 50 mL). Combined organic phases were dried over sodium sulfate, filtered and concentrated under reduced pressure. The crude product was purified by column chromatography (EtOAc : PE 50 : 50 to 100 : 0) and HPLC afterwards. Product fractions were lyophilised yielding compound **SW-C210** as a white solid (72 mg, 0.18 mmol, 28 %).

**TLC:** R_f_ = 0.41 (PE/ EtOAc 1 : 1).

**^1^H NMR** (500 MHz, MeOH-d_4_) δ 6.79 (dd, *J* = 1.3, 0.6 Hz, 1H), 6.76 (dd, *J* = 1.9, 1.1 Hz, 2H), 6.35 (q, *J* = 3.3 Hz, 2H), 5.91 (s, 2H), 4.30 (s, 2H), 3.73 (s, 2H), 3.21 (s, 2H), 2.39 (s, 3H).

**^13^C NMR** (126 MHz, MeOH-d_4_) δ 171.9, 154.2, 149.2, 148.3, 133.6, 122.8, 122.0, 113.8, 113.2, 109.1, 109.1, 102.3, 60.1, 54.2, 43.6, 42.8.

**HRMS (ESI)** calculated for C_16_H_18_BrN_2_O_4_ (M (^79^ Br) + H^+^): 381.0450, found: 381.0453; calculated for C_16_H_17_BrN_2_O_4_ (M (^81^ Br) + H^+^): 383.0429, found: 383.0438. Further C26-analogs were obtained according to Scheme 2.

**Scheme 2.**
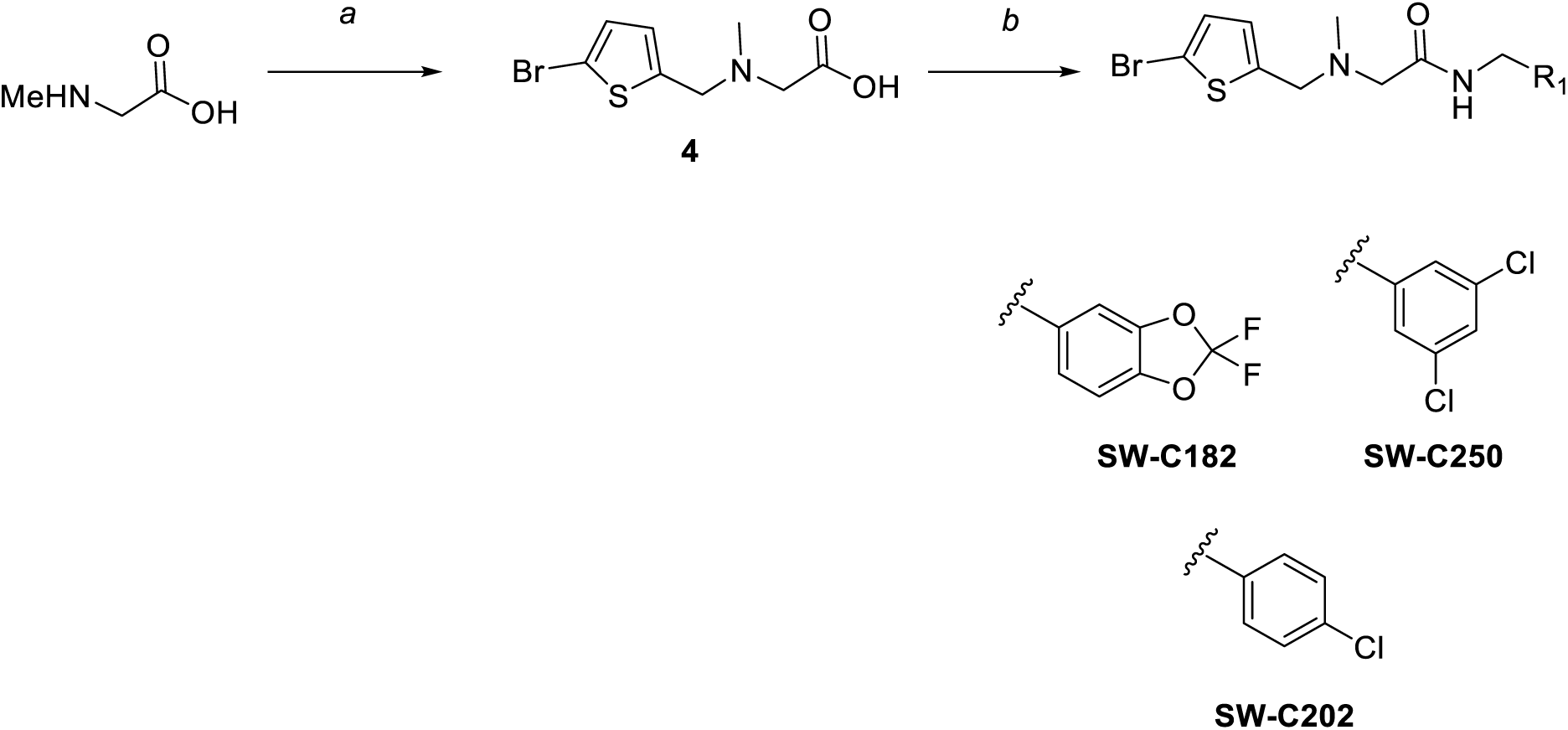
Reagents and conditions. (*a*) 5-bromothiophene-2-carbaldehyde, HOAc, THF, r. t., 10 min, *then* NaBH(OAc)_3_, r. t., overnight, 97%; (*b*) iBuOCOCl, Et_3_N, THF, 0 °C to r. t., 2 h, *then* R_1_CH_2_NH_2_, r. t., 2 h; *or* R_1_CH_2_NH_2_, HATU, DIPEA, DMF.

#### *N*-((5-bromothiophen-2-yl)methyl)-*N*-methylglycine (4)

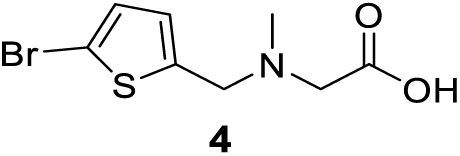

To a solution of sarcosine (200 mg, 2.24 mmol, 2 eq.) in dry MeOH (5 mL) was added triethylamine (313 µL, 227 mg, 2.24 mmol, 2 eq.) and 5-bromothiophene-2-carboxaldehyde (121 µL, 214 mg, 1.12 mmol, 1 eq.). The reaction mixture was stirred for 14 h at r. t. under argon atmosphere, then NaBH_4_ (169 mg, 4.48 mmol, 4 eq.) was added and the solution was stirred for another 16 h at the same conditions. All volatile components were removed under reduced pressure and the crude product was purified by RP flash chromatography (C18; MeCN : H_2_O) to afford the compound **4** as a white solid (286 mg, 1.08 mmol, 97 %).

**^1^H NMR** (500 MHz, MeOH-d_4_) δ 6.92 (d, *J* = 3.7 Hz, 1H), 6.77 (dt, *J* = 3.7, 0.8 Hz, 1H), 3.82 (d, *J* = 0.6 Hz, 2H), 3.03 (s, 2H), 2.32 (s, 3H).

**^13^C NMR** (126 MHz, MeOH-d_4_) δ 178.1, 145.0, 130.5, 128.2, 112.1, 61.2, 56.4, 42.4, 35.9.

**HRMS (ESI)** calculated for C_8_H_11_BrNO_2_S (M (^79^ Br) + H^+^): 263.9694, found: 263.9704; calculated for C_8_H_10_BrNO_2_S (M (^81^ Br) + H^+^): 265.9673, found: 265.9683.

#### 2-(((5-Bromothiophen-2-yl)methyl)(methyl)amino)-N-((2,2-difluorobenzo[d][1,3]dioxol-5-yl)methyl)acetamide (SW-C182)

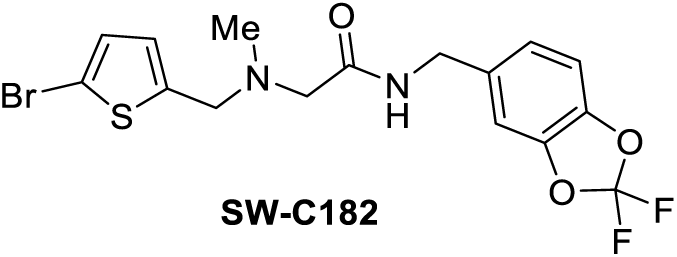

To the compound **39** (30 mg, 0.113 mmol, 1.0 eq.) in THF (2 mL) was added TEA (80 µL, 0.567 mmol, 5.0 eq.) under nitrogen atmosphere at 0 °C. Isobutylchloroformate (23 µL, 0.175 mmol, 1.50 eq.) was added drop wise at 0 °C. Stirred for 2 h at r. t. and (2,2-difluorobenzo[d][1,3]dioxol-5-yl)methanamine (21 mg, 0.113 mmol, 1.0 eq.) was added at r. t. in a single portion and stirred at r. t. for an additional 2 h. Reaction mixture was quenched with saturated NaHCO_3_ solution (20 mL) and extracted with Et_2_O (2 × 20 mL). Combined organic layers were dried over anhydrous Na_2_SO_4_, filtered and concentrated under vacuum. Crude compound was purified using column chromatography (3:2, EtOAc:PE) to obtain compound **SW-C182** (10 mg, 20%) as colorless oil.

**TLC analysis:** 2:1, EtOAc: PE, R_f_: 0.40 (Stain: KMnO_4_/UV).

**^1^H NMR** (400 MHz, CDCl_3_): δ = 6.97-7.05 (m, 3H), 6.89 (d, *J* = 3.6 Hz, 1H), 6.71 (bs, 1H), 4.44 (d, *J* = 5.6 Hz, 2H), 3.80 (bs, 2H), 3.20 (bs, 2H), 2.40 (s, 3H).

**^13^C NMR** (101 MHz, CDCl_3_): δ = 170.0, 144.1, 143.2, 134.7, 131.8, 129.8, 129.3, 123.0, 109.6, 109.3, 56.9, 42.9, 31.1.

**HRMS (ESI)** calculated for C_16_H_16_BrF_2_N_2_O_3_S (M + H^+^): 433.0033, found: 433.0029.

#### 2-(((5-Bromothiophen-2-yl)methyl)(methyl)amino)-N-(3,5-dichlorobenzyl)acetamide (SW-C250)

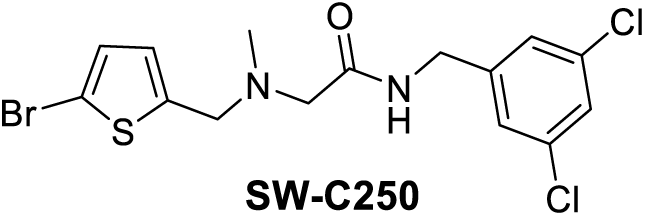

To a stirring solution of compound **4** (100 mg, 379 µmol, 1 eq.) in dry DMF (6 mL) was added DIPEA (198 µL, 147 mg, 1140 µmol, 3 eq) and HATU (173 mg, 454 µmol, 1.2 eq.). After 10 minutes, 3,4-dichlorobenzylamine (61 µL, 80 mg, 454 µmol, 1.2 eq.) was added and the solution was stirred for 20 h at r. t. under argon atmosphere. The mixture was diluted with water (10 mL) and extracted with CH_2_Cl_2_ (3×30 mL). Phases were separated and combined organic phases were washed with brine (20 mL) and dried over sodium sulfate. The solution was filtered, washed with CH_2_Cl_2_ and concentrated under reduced pressure. The residue was purified by C18 flash (MeCN : H_2_O) and then by preparative HPLC (MeCN : H_2_O+0.1% HCOOH). Product containing fractions were lyophilised to yield compound **SW-C250** (28.5 mg, 18%).

**^1^H NMR** (500 MHz, CD_3_OD) δ 7.33 (t, *J* = 2.0 Hz, 1H), 7.27 (d, *J* = 2.0 Hz, 2H), 6.94 (d, *J* = 3.7 Hz, 1H), 6.79 (dt, *J* = 3.7, 0.9 Hz, 1H), 4.39 (s, 2H), 3.79 (d, *J* = 0.9 Hz, 2H), 3.10 (s, 2H), 2.36 (s, 3H).

**^13^C NMR** (126 MHz, CD_3_OD) δ 173.64, 145.03, 144.45, 136.29, 130.88, 128.43, 128.26, 127.33, 112.85, 60.33, 57.43, 43.48, 42.91.

**HRMS (ESI)** calculated for C_15_H_16_BrCl_2_N_2_OS (M (^79^ Br) + H^+^): 420.9544, found: 420.9548; calculated for C_15_H_16_BrCl_2_N_2_OS (M (^81^ Br) + H^+^): 422.9523, found: 422.9558.

#### 2-(((5-Bromothiophen-2-yl)methyl)(methyl)amino)-N-(4-chlorobenzyl)acetamide (SW-C202)

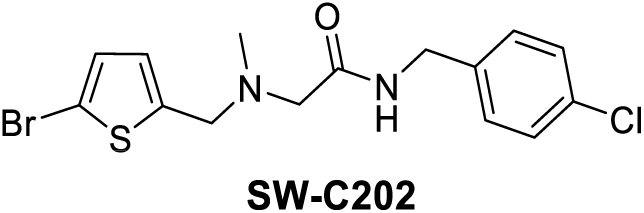

To a stirring solution of compound **4** (200 mg, 0.75 mmol, 1 eq.) in dry DMF (6 mL) was added DIPEA (395 µL, 293 mg, 2.27 mmol, 3 eq) and HATU (345 mg, 0.90 mmol, 2 eq.). After 10 minutes, 4-chlorobenzylamine (110 µL, 128 mg, 0.90 mmol, 1.2 eq.) was added and the solution was stirred for 3 h at r. t. under nitrogen atmosphere. The reaction mixture was diluted with water (10 mL) and extracted with CH_2_Cl_2_ (3×30 mL). Combined organic phases were washed with brine (20 mL), dried over sodium sulfate, filtered, washed with CH_2_Cl_2_ and concentrated under reduced pressure. The residue was taken up in DMSO and purified by HPLC twice. Product containing fractions were lyophilised to yield compound **SW-C202** (50 mg, 128 µmol, 17 %) as a yellow oil.

**^1^H NMR** (500 MHz, DMSO-d_6_) δ 8.41 (s, 1H), 7.39 – 7.36 (m, 2H), 7.29 – 7.26 (m, 2H), 7.10 (d, *J* = 3.3 Hz, 1H), 6.90 (s, br, 1H), 4.29 (d, *J* = 6.1 Hz, 2H), 3.90 (s, 2H), 3.19 (s, 2H), 2.34 (s, 3H).

**^13^C NMR** (176 MHz, DMSO-d_6_) δ 158.1, 140.4, 138.5, 131.3, 130.8, 129.9, 129.1, 128.8, 128.2, 127.9, 55.2, 42.9, 35.8.

**HRMS (ESI)** calculated for C_15_H_17_BrClN_2_OS (M (^79^ Br) + H^+^): 386.9933, found: 386.9954; calculated for C_15_H_16_BrClN_2_OS (M (^81^ Br) + H^+^): 388.9913, found: 388.9905.

2-(((5-Bromothiophen-2-yl)methyl)(methyl)amino)-*N*-(3,4-dihydroxybenzyl)acetamide (**SW-C170**) was synthesized according to the Scheme 3.

**Scheme 3.**
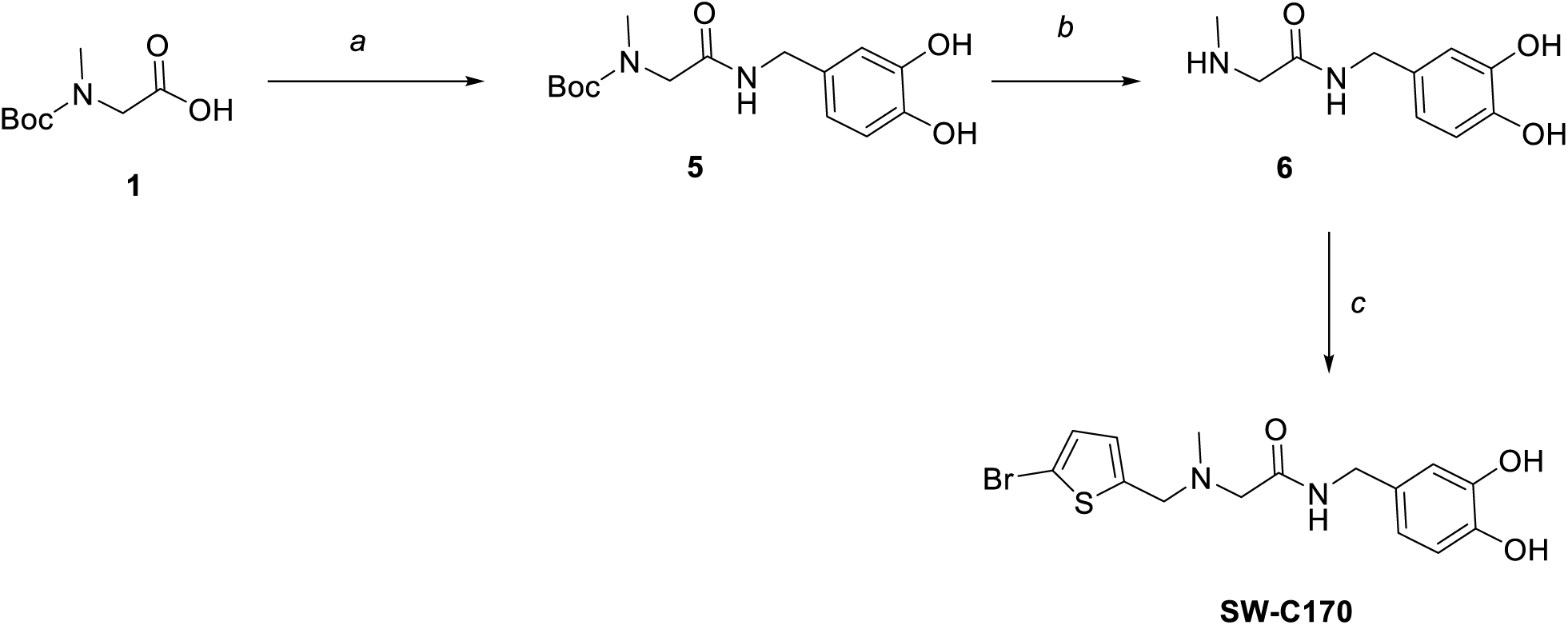
Reagents and conditions. (*a*) iBuOCOCl, Et_3_N, THF, 0 °C to r. t., 2 h, *then* (HO)_2_C_6_H_3_CH_2_NH_2_, r. t., 2 h, 42%; (*b*) TFA, CH_2_Cl_2_, r. t., 5 h, (*c*) 2-bromo-5-(bromomethyl)thiophene, Et_3_N, THF, 16 h, 68% over two steps.

#### *tert*-Butyl (2-((3,4-dihydroxybenzyl)amino)-2-oxoethyl)(methyl)carbamate

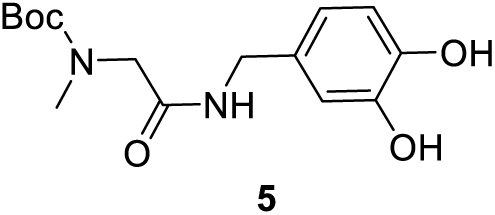

To the Boc-sarcosine **1** (0.270 g, 1.440 mmol, 1.0 eq.) in THF (10 mL) was added triethylamine (0.80 mL, 5.750 mmol, 4.0 eq.) under nitrogen atmosphere at 0 °C. Isobutylchloroformate (0.210 mL, 1.580 mmol, 1.10 eq.) was added drop wise at 0 °C. Stirred for 2 h min at r. t. and 3,4-dihydroxybenzylamine (0.200 g, 1.440 mmol, 1.0 eq.) was added at r. t. in a single portion and stirred at r.t. for 16 h. Reaction mixture was quenched with saturated NaHCO_3_ solution (20 mL) and extracted with EtOAc (2 × 20 mL). Combined organic layers are dried over anhydrous Na_2_SO_4_, filtered and concentrated under reduced pressure. The crude compound was purified using column chromatography (1:20, MeOH: CH_2_Cl_2_) to obtain compound **5** (0.189 g, 42%) as colorless liquid.

**TLC analysis:** 20:1, CH_2_Cl_2_: MeOH, R_f_ : 0.30 (Stain: Ninhydrin/UV).

**^1^H NMR** (400 MHz, CD_3_OD): δ = 6.59–7.03 (m, 3H), 4.30 (s, 2H), 3.96 (m, 2H), 2.95 (s, 3H), 1.36 (m, 9H).

**^13^C NMR** (101 MHz, CD_3_OD): δ = 171.4, 157.1, 146.4, 145.7, 120.4, 120.1, 116.2, 81.6, 53.4, 43.8, 36.3, 28.5

**HRMS (ESI)** calculated for C_15_H_22_N_2_O_5_Na (M + Na^+^): 333.1426, found: 333.1426.

#### *N*-(3,4-Dihydroxybenzyl)-2-(methylamino)acetamide

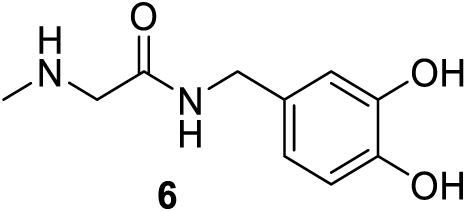

To the compound **5** (50 mg, 0.160 mmol, 1.0 eq.) in CH_2_Cl_2_ (2 mL) was added TFA (0.7 mL) at r. t. and stirred for 5 h. The reaction mixture was concentrated under reduced pressure to afford crude **6** that was used without further characterization.

#### 2-(((5-Bromothiophen-2-yl)methyl)(methyl)amino)-N-(3,4-dihydroxybenzyl)acetamide (SW-C170)

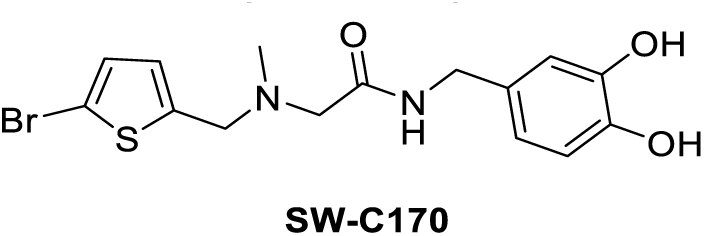

2-Bromo-5-(bromomethyl)thiophene (40 mg, 0.17 mmol, 1.0 eq.) was dissolved in dry THF (0.5 mL) and triethylamine (242 µL, 1.68 mmol, 10.0 eq.) was added and the solution was cooled to 0 °C. Compound **6** (54 mg, 0.16 mmol, 1.00 eq.) was added and the mixture was stirred 5 min at 0 °C and at r. t. for 16 h. THF was removed under reduced pressure. Crude mixture was washed with sat. NaHCO_3_ solution (10 mL) and extracted with ethyl acetate (3 × 10 mL). The combined organic layers were dried over anhydrous Na_2_SO_4_, filtrated and evaporated under reduced pressure. Crude product was purified by column chromatography (MeOH : CH_2_Cl_2_, 1 : 20) to obtain product **SW-C170** (44 mg, 68%) as yellow oil.

**TLC analysis:** 1:20, MeOH: CH_2_Cl_2_, R_f_ : 0.8 (Stain: KMnO_4_/UV).

**^1^H NMR** (400 MHz, CD_3_OD): δ = 6.92 (d, *J* = 3.7 Hz, 1H), 6.73 (dd, *J* = 8.2, 6.0 Hz, 3H), 6.61 (dd, *J* = 8.1, 2.0 Hz, 1H), 4.26 (s, 2H), 3.76 (s, 2H), 3.07 (s, 2H), 2.32 (s, 3H).

**^13^C NMR** (101 MHz, CD_3_OD): δ = 172.7, 146.4, 145.7, 144.8, 131.3, 130.7, 128.2, 120.2, 116.3, 116.0, 112.6, 60.4, 57.2, 43.6, 43.1.

### *In vitro* cell toxicity assay

*In vitro* toxicity was assessed in HeLa cells (ATCC® CCL-2) using the ApoTox-Glo^TM^ Triplex Assay (Promega, USA) according to the manufacturer’s instructions. HeLa cells (10^4^ cells per well) were seeded into a 96-well plate and incubated for 24 h at 37 °C, 5% (v/v) CO_2_. HeLa cells were then treated with the compound at different concentrations or 1% (v/v) DMSO in Dulbecco’s Modified Eagle Medium (DMEM) without phenol red for 18 h. Fluorescence intensity was measured with TECAN Spark® microplate reader. Cytotoxicity (RFU) and viability (RFU) values were plotted to calculate the TC_50_ using GraphPad Prism.

### Toxicology studies

C26 was formulated in 5% Tween 20 (v/v), 50% lecithin (from soy bean: 40 mg in 800 μl distilled water), 45% PBS at 0.6, 2, and 6 mg/ml for oral administration. A dosing volume of 5 ml/kg was applied. C26 was administered orally to groups of 3 male ICR mice (23 ± 3 g) at an initiating dose of 3 mg/kg in a Maximum Tolerated Dose (MTD) setup. The animals received an initial dose of 3 mg/kg. If the animals survived for 72 hours, the 10 mg/kg group was tested. If the animals survived for 72 hours, the 30 mg/kg group was tested. Experiments were performed by Pharmacology Discovery Services Taiwan, Ltd., in general accordance with the “Guide for the Care and Use of Laboratory Animals: Eighth Edition” (National Academies Press, Washington, D.C., 2011). The animal care and protocol were reviewed and approved by the IACUC at Pharmacology Discovery Services Taiwan, Ltd.

### SipA injection assay

SipA injection assay was performed using the split-Nanoluc (HiBiT/LgBiT) system as described previously^37,38^. In brief, NanoLuc is split into two parts: LgBiT, comprising 10 of the 11 β-strands of the luciferase, and HiBiT, a short peptide with high affinity to LgBiT, contributing the missing β-strand to make a functional luciferase. *S*. Typhimurium strains expressing SipA-HiBiT were grown in LB supplemented with 0.3 M NaCl, and either the compound at different concentrations or 1% (v/v) DMSO. Cultures were incubated at 37 °C with shaking at 180 rpm until an OD_600_ of 0.9 was reached. Bacteria were then pelleted and washed twice with HBSS. The bacterial suspension was diluted to obtain a MOI of 50, and then used to infect the LgBiT-expressing HeLa cell. Luminescence was measured using a Tecan Sparc Multimode reader.

### Plasmid-based SipA secretion assay

The plasmid pT10-SipA-NLuc was electroporated into electrocompetent cells of *S*. enterica clinical isolates. Bacteria at an initial OD_600_ of 0.02 were grown for 5 h in 1 ml LB supplemented with 0.3 M NaCl, kanamycin 50 µg/ml, and either the compound at 100 µM or 1% (v/v) DMSO. Cultures were pelleted, and then 25 µl of the supernatant was transferred to a 384-well plate to measure luminescence as previously described.

### SiiE cell surface retention assay

HiBiT was inserted chromosomally into SiiE at position K5411, using the suicide plasmid pMIB8021 (pSB890-*siiE*::K5411HiBiT). Bacteria were grown for 5 h in SPI-4-inducing conditions (LB supplemented with 0.3 M NaCl). 0.5 OD units were harvested at 10000 × *g*, 2 min, 4 °C. Cell pellets were washed twice with cold PBS and then resuspended to a final concentration of 0.5 OD units. 25 µl from the bacterial suspension were transferred into 384-well plate. The Nano-Glo HiBiT Extracellular Buffer and corresponding substrate were prepared according to manufacturer’s instructions. 25 µl of the Nano-Glo HiBiT Extracellular buffer-substrate mix were added to each sample and then incubated at room temperature for 10 min. Luminescence was measured with a Tecan Spark microplate reader.

### Invasion assay into HeLa cells

In a white 24-well plate (NUNC), 10^5^ HeLa cells were seeded in 350 μl DMEM (Gibco), 24 h before the infection. Bacteria were grown in LB supplemented with 0.3 M NaCl, and either the compound at different concentrations or 1% (v/v) DMSO, at 37 °C with shaking at 180 rpm until an OD_600_ of 0.9 was reached. Bacteria were then pelleted and washed twice with HBSS. The bacterial suspension was diluted to obtain a MOI of 20 to infect HeLa cells. The plate was centrifuged for 5 min at 300 × g to synchronize the infection and then incubated for 25 min, 37 °C, 5% (v/v) CO_2_, to allow *Salmonella* invasion into HeLa cells. To quantify the invasiveness, the cells were first washed 3 times with 500 µl prewarmed PBS. The remaining extracellular bacteria were killed with 500 µl DMEM supplemented with 100 µg/ml gentamicin. HeLa cells were incubated for 1 h, then washed three times with prewarmed PBS and lysed with 500 µl of 0.5% (v/v) SDS in PBS for 5 min at 37 °C on a shaking platform. The lysate was serial diluted in PBS-T (PBS with 0.05% (v/v) Tween 20) to determine the CFUs by plating on LB medium supplemented with streptomycin 50 µg/ml.

### Invasion assay into MDCK cells

MDCK (NBL-2) cells were seeded in a 24-well plate at a density of 10^5^ cells per well in 1 ml MEM (Gibco) and grown for 5-6 days, to allow polarization. Cells were washed with fresh medium every two days. After 5 days, each well contained approximately 1.8 × 10^6^ MDCK cells. Bacteria were grown as for the invasion assay of HeLa cells, and diluted to obtain a MOI of 5. The plate was centrifuged for 3 min at 300 × *g* to synchronize the infection, then incubated for 25 min, 37 °C, 5% (v/v) CO_2_, to allow *Salmonella* invasion into MDCK cells. The quantification of invasiveness was carried out as described above for HeLa cells.

### PipB2 secretion assay

PipB2 injection assay was performed using the split-Nanoluc (HiBiT/LgBiT) system as described for SipA. In a white 96 well plate (NUNC) with optical bottom, 10^4^ HeLa LgBiT cells were seeded in 100 μl DMEM, 24 h before the infection. The *pipB2*-HiBiT strain was grown in LB-NaCl at 37 °C at 180 rpm until an OD_600_ of 0.9. Bacteria were washed twice with HBSS (Serva) and then used to infect pre-seeded HeLa cells on a 96-well plate with an MOI of 100. The plate was centrifuged at 300 *x g* for 3 min and then incubated for 1h to allow invasion. Afterwards, HeLa cells were washed with DMEM and incubated for 1h with DMEM supplemented with 100 μg/ml gentamicin to eliminate non-invading bacteria. The medium was replaced by DMEM with 16 μg/ml gentamicin, supplemented with either DMSO (1%) or C26 (100 µM). Cells were incubated at 37 °C with 5% (v/v) CO_2_ for 14h, then washed twice with 1x PBS before luminescence measurements were carried out following manufacturer’s instructions.

### Western blotting analysis

Western blot analysis to quantify secreted and non-secreted proteins was carried out as previously described^63^. After a 5h culture in SPI-1-inducing conditions, cultures were centrifuged at 10,000 × g for 2 min at 4 °C to separate cell pellets for the quantification of target proteins in whole cells, and the supernatant for the quantification of secreted proteins. The supernatants were first filtered with a 0.22 µm pore size filter. Sodium deoxycholic acid was then added to a final concentration of 0.1% (w/v) followed by a protein precipitation with 10% trichloroacetic acid (v/v) for 30 min at 4 °C. The samples were pelleted by centrifugation at 20,000 *x g* for 20 min at 4 °C to retrieve precipitated proteins which were finally washed with acetone before resuspension in SDS PAGE loading buffer. Whole cells samples were directly resuspended in SDS PAGE loading buffer.

### Transcriptome analysis

*Salmonella* was grown for 5 h at 37 °C in LB medium in presence of 100 µM C26 or 1% (v/v) DMSO as a control. After growth, bacteria were harvested by centrifugation, and total RNA was isolated using Qiagen RNeasy mini kit according to manufacturer’s protocol. RNA sequencing was performed on a HiSeq2500 with a 2 × 125 bp paired end read protocol.

Illumina Casava software was used to de-multiplex the sequenced reads providing individual raw fastq sample files. Raw fastq files was pre-filtered using the chastity filter to remove reads that contain a “Y” flag. FastQC ((http://www.bioinformatics.babraham.ac.uk/projects/fastqc/), version v0.11.4) was used to determine quality of the resulting fastq files. Subsequently, an adapter trimming/removal step was conducted with Cutadapt ((https://pypi.python.org/pypi/cutadapt/), version 1.8.3). This process used FastQC output (see step before) to identify reads that showed a match to some typical overrepresented (Illumina) sequences/adapters. TopHat2 ((https://ccb.jhu.edu/software/tophat/index.shtml), version v2.0.12) was used as aligner to map the remaining quality controlled reads to the Salmonella genome. Read counting to features (e.g., genes or exons) in the genome was performed with HTSeq-count (http://www-huber.embl.de/users/anders/HTSeq/doc/count.html), version 0.6.0.). Counting was performed using “union” mode on the feature “gene”. The stranded option was also set to “–stranded=no” to indicate to count features on both strands. Further, the -r parameter from HTSeq was set to “pos”, as this is the default of the output of Tophat’s ‘accepted_hits.bam’ mapping file. For differential expression analysis the raw read count table resulting from HTSeq counting is used and fed into the R package DESeq2 (version 1.10.1). Graphs were also produced in the R language (R version 3.2.1) mainly using the R package ggplot2 (version 2.2.0). Reports were produced using the R package rmarkdown (version 1.3).

### Cell-based assay monitoring HilD activity

A cell-based reporter gene assay was used to quantify HilD transcriptional activity. Strain P*_hilA_*-sfGFP and the isogenic Δ*hilD* P*_hilA_*-sfGFP were used for this purpose. Cultures were performed in 1 ml LB supplemented with 0.3 M NaCl. Compounds at different concentrations or 1% (v/v) DMSO were added to the corresponding tubes. Cultures were incubated at 37 °C, 180 rpm, for 5 h. Cells were pelleted, resuspended in 100 µl PBS and then transferred into a 96-well black clear bottom plate (Thermo scientific, USA). Fluorescence intensity was measured with Tecan Spark microplate reader with an excitation wavelength of 485 nm and emission wavelength of 510 nm. Dose-response curves and IC_50_ values were calculated using CDD Vault®.

### Alanine scan

A plasmid-based assay in a ΔSPI-1 background strain was developed to facilitate the introduction of point mutations in *hilD*. The Rha cassette (*rhaS*, *rhaR*, and P*rha*) was first deleted from the pT10 backbone^64^. The fragment P*hilD*-*hilD*-P*hilA*-sfGFP was then inserted upstream of the terminator *rrnB*. *hilD* was then deleted from the resulting plasmid to serve as a negative control. Site-directed mutagenesis was performed using using KOD polymerase (Novagen). The cell-based fluorescence assay was performed as described above.

### Molecular modelling for HilD’s binding site prediction and C26’s binding mode suggestion

Molecular modelling of the HilD target and complete protocol for MD simulations are described in the supporting information Methods

#### Protein Structure Prediction and Binding site prediction

The structural model of the N-terminal truncated Salty HilD (UniProt ID: P0CL08, starting at Ser37) was retrieved from the AlphaFold Protein Structure Database^65^. All structure models can be found in the supplementary material. System preparation and docking calculations were performed using the Schrödinger Drug Discovery suite for molecular modelling (version 2022.1). Protein−ligand complex was prepared with the Protein Preparation Wizard to fix protonation states of amino acids, add hydrogens, and fix missing side-chain atoms, where we selected the most likely ionization state as proposed by the software, and the structures were minimized. Currently, DNA-binding interactions associated with the carboxy-terminal domain (CTD) of other AraC-like proteins’ CTD have been inferred from static models based on similar MarA and Rob proteins^66–69^. However, there are no structural studies focused on the full length HilD protein regarding how the amino-terminal domain (NTD) and CTD interact with each other, and how potential ligands interfere with this geometry. In this sense, for each system, namely monomer (M), monomer with DNA (MDNA) systems were generated. HilD+DNA was generated using the coordinates from the CTD with bound DNA modelled based on the MarA-DNA structure (PDB ID: 1BL0^68^, resolution: 2.3 Å) followed by energy minimization. Potential binding pockets were predicted using SiteMap^70^.

#### Molecular Docking

All ligands for docking were drawn using Maestro and prepared using LigPrep^54^ to generate the 3D conformation, adjust the protonation state to physiological pH (7.4), and calculate the partial atomic charges with the OPLS4 force field^55^. Docking studies with the prepared ligands were performed using Glide (Glide V7.7)^56,57^ with the flexible modality of induced-fit docking with extra precision (XP), followed by a side-chain minimization step using Prime. Ligands were docked within a grid around 12 Å from the centroid of the predicted binding site pocket, as determined using SiteMap.

#### Molecular dynamics simulation

MD simulations were carried out using Desmond^32^ with the OPLS4 force-field^55^. The simulated system encompassed the protein-ligand complexes, a predefined water model (TIP3P^71^) as a solvent, and counterions. The system was treated in a cubic box with periodic boundary conditions specifying the box’s shape and size as 13 Å distance from the box edges to any atom of the protein. In all simulations, we used a time step of 1 fs, the short-range coulombic interactions were treated using a cut-off value of 9.0 Å using the short-range method, while the Smooth Particle Mesh Ewald method (PME) handled long-range coulombic interactions^72^. Initially, the system’s relaxation was performed using Steepest Descent and the limited-memory Broyden-Fletcher-Goldfarb-Shanno algorithms in a hybrid manner, according to the established protocol available in the Desmond standard settings. During the equilibration step, the simulation was performed under the NPT ensemble for 5 ns implementing the Berendsen thermostat and barostat methods^73^. A constant temperature of 310 K was kept throughout the simulation using the Nose-Hoover thermostat algorithm^74^ and Martyna-Tobias-Klein Barostat^75^ algorithm to maintain 1 atm of pressure. After minimization and relaxation of the system, we continued with the production step of at least 2 µs, with frames being recorded/saved every 1,000 ps. Five independent replicas were produced for each compound, resulting in a total of ∼10 µs simulation/ligand. Trajectories and interaction data are available on the Zenodo repository^76^. The representative structures were selected by inspecting changes in the Root-mean-square deviation (RMSD), meaning for figures a representative frame was selected at random at points of the trajectory where the RMSD were not fluctuating, after equilibration. Extended data Fig. 6 represents the variation of the RMSD values along with the simulation, for both template crystal structures and simulations with docking pose. Additionally, the changes in the Root-mean-square fluctuation (RMSF), normalized by residue for the protein backbone, are displayed in Extended data Fig. 7.

##### MM-GBSA binding energy calculations

Molecular mechanics with generalized Born and surface area (MM-GBSA) predicts the binding free energy of protein-ligand complexes and the ranking of ligands based on the free energy could be correlated to the experimental binding affinities, especially in a congeneric series. Every 50^th^ frame from the simulations was considered for the calculations. These were used as input files for the MM-GBSA calculations with thermal_mmgbsa.py script from the Schrödinger package, using Prime ^77^. Calculated free-binding energies are represented by the MM/GBSA and normalized by the number of heavy atoms (HAC), according to the following formula: ligand efficiency = (Binding energy) / (1 + ln(HAC)) and is expressed in kcal/mol.HAC, where HAC is the Heavy Atom Count. Trajectory distances between specific secondary structure elements were calculated using their centers of mass with the Maestro script trj_asl_distance.py (Schrödinger LLC), using the carbon alpha coordinate of specific amino acids as a reference. Energy distribution is depicted in Extended data Extended data Fig. 5.

### Recombinant protein expression and purification

The *hilC* gene was inserted into the pET-21a(+) vector, with an N-terminal His_6_ tag followed by a TEV protease cleavage site. The *hilD* gene was cloned into the pET-24a(+) vector, with an N-terminal His_6_-SUMO fusion. HilD mutants were cloned into the *hilD* construct by site-directed mutagenesis. Proteins were expressed in *E. coli* C41(DE3)^78^ cells using lysogeny broth (LB) medium. An overnight culture was inoculated into LB medium, grown at 37 °C until an OD_600_ of 0.6-0.8 was reached and induced by the addition of 0.5 mM isopropyl β-D-1-thiogalactopyranoside (IPTG). Cells were incubated with shaking overnight at 25 °C, collected by centrifugation (11,800 × *g*, 4 °C), and resuspended in buffer A (50 mM NaH_2_PO_4_, pH 7.0, 300 mM NaCl, 10 mM imidazole) supplemented with DNase and one cOmplete™ EDTA-free protease inhibitor cocktail tablet (Roche #11 873 580 001). Cells were lysed using a French press (2x, 16,000 psi) and cell debris removed by centrifugation (95,000 × *g*, 1 h, 4 °C). The supernatant was filtered (0.40 μm) and loaded to a Ni-NTA column. Bound proteins were washed first with 20% (v/v) buffer B (50 mM NaH_2_PO_4_, pH 7.0, 300 mM NaCl, 250 mM imidazole), and then eluted with 100% (v/v) of it.. The eluted SUMO-HilD protein was supplemented with SUMO protease (250 μg) to cleave the His_6_-SUMO tag and dialyzed overnight at room temperature against buffer A. For HilC, the eluted protein was supplemented with TEV protease (1 mg) and dialyzed overnight at 6 °C against Buffer C (50 mM NaH_2_PO_4_, pH 7.0, 400 mM NaCl). The dialysed protein was reapplied to the Ni-NTA column, equilibrated with buffer A, and the column was washed with 25% (v/v) buffer B to elute the cleaved protein. In the case of HilC, a higher NaCl concentration of 500 mM was used for both Ni-NTA purification steps. Proteins were then concentrated using Amicon® Ultra Centrifugal Filters and loaded to a size exclusion chromatography column (Superdex™ 75 26/60) equilibrated with SEC Buffer (50 mM NaH_2_PO_4_, pH 7.0, 200 mM NaCl). Eluted fractions containing purified protein were combined, concentrated, and stored in aliquots at −80 °C. Protein purity was assessed by SDS-PAGE and protein concentration was determined from UV absorbance at 280 nm, measured using a NanoPhotometer® NP80 (IMPLEN).

### Electrophoretic mobility shift assays (EMSAs)

EMSAs were performed similarly to as described previously^39^ using a 62 base pair dsDNA fragment of the *hilA* promoter, encompassing the A1 binding site^79^. Double stranded DNA fragments were generated by melting the complementary primers PhilA_A1_f / PhilA_A1_r (Supplementary Tab. 10) together in TE Buffer (10 mM Tris pH 8.0, 1 mM EDTA) at 95 °C for 10 min before slowly cooling to room temperature. The forward primer was modified with a 5’-Cy5 fluorescent dye for detection. 600 nM of protein was incubated with 50 nM of labelled DNA in EMSA buffer (20 mM Tris, pH 8.0, 100 mM KCl, 100 μM EDTA, 3% glycerol). C26 was diluted first in DMSO and subsequently 1:100 into the protein-DNA sample. Samples were incubated at 37 °C for 15 min, supplemented with diluted DNA loading dye, and separated on a 1.5 mm thick, 6% (w/v) TBE polyacrylamide gel at 6 °C at a constant voltage of 100 V. Gels were imaged using a ChemiDoc^TM^MP imaging system (Bio-Rad).

### Nanoscale differential scanning fluorimetry (nanoDSF)

Thermal stability of proteins was determined using nanoscale differential scanning fluorimetry (nanoDSF), with runs performed on a Prometheus NT.48 (NanoTemper Technologies). A two-fold serial dilution series of C26 was prepared in DMSO. C26 was added to HilD or HilC (5 μM) in SEC buffer, giving a final DMSO concentration of 1% (v/v). Samples were incubated for > 20 min at room temperature and centrifuged for 2 min prior to loading of standard capillaries (#PR-C002). Samples were heated from 20 to 80 °C with a temperature gradient of 0.5 °C min^−1^. Melting temperatures were calculated from changes in the fluorescence ratio (350/330 nm), using PR.Stability Analysis v1.0.3 and a temperature range of 40-70 °C for curve fitting. Data analysis was performed using Prism 8.4 (GraphPad). The change in HilD melting temperature, T_m_, was fitted as a function of ligand concentration, using equations (1) and (2) to yield apparent affinity (K_d,app_) values.

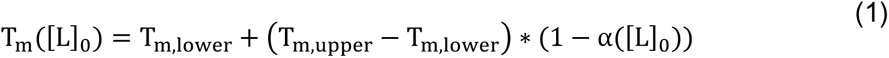

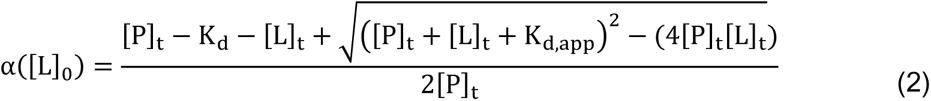

where [P]_t_ and [L]_t_ are the total protein and ligand concentrations, respectively.

### SEC-MALS

SEC-MALS experiments were performed using a Superdex™ 75 Increase 10/300 GL column (Cytiva) coupled to a miniDAWN Tristar Laser photometer (Wyatt) and a RI-2031 differential refractometer (JASCO). HilD (100 μM) was incubated with 1% (v/v) of either DMSO or C26 (10 mM dissolved in DMSO) for 20 minutes at room temperature. 50 μL of HilD samples were loaded onto the SEC column, equilibrated with SEC buffer, and separated using a flow rate of 0.5 ml min^−1^. Data analysis was carried out with ASTRA v7.3.0.18 software (Wyatt).

### BS^3^ cross-linking

BS^3^ crosslinking of HilD was performed as previously described^39^. In summary, HilD (10 μM) was first incubated with C26 or oleic acid in SEC buffer for 20 min at room temperature, with a final DMSO concentration of 1% (v/v). HilD was then cross-linked by incubation with 0.2 mM BS^3^ (Thermo Fisher Scientific Pierce, A39266) at room temperature for 1 hour, before the reaction was quenched by the addition of 50 mM Tris pH 7.5. Samples were analyzed using SDS-PAGE and visualized by silver staining.

### NMR Spectroscopy

Assignment of the C26 spectrum was readily available from considerations of chemical shifts and ^3^J couplings in a 1D spectrum. Saturation transfer difference experiments^80^ were acquired on the HilD C26 complex using a ligand concentration of 60 µM and a protein concentration of 16 µM. Measurements were carried out at 800 MHz on a Bruker AVIII Spectrometer. Spectra were acquired at 298 K with 4096 scans and 16384 acquired data points, with saturation at 0.8 ppm targeting protein methyl groups. An STD build-up series was acquired using saturation times of 400, 800, 1200, 2000 and 3000 µs.

The expected STD intensities were back calculated for frames of the MD trajectories using the CORCEMA algorithm^40–42^ implemented within the SHINE NOESY back calculation suite (in house software). This implementation uses ligand chemical shift and coupling data (Extended data Fig. 8), plus estimated of line width to simulate the STD spectrum. This allows direct comparison of experimental and back-calculated spectra with an R-factor based on the RMSD^81^. Parameters provided to the program include the protein and ligand concentrations, ligand affinity (35 µM) an estimate of the protein correlation time (12 ns), ligand chemical shifts and couplings. The non-instantaneous saturation model was used with saturated protons selected on the basis of chemical shifts predicted for the HilD AlphaFold model using SHIFTX2^82^. Quantitative comparison was carried out on the aromatic region of the spectrum (5.5-7.2 ppm), as these signals are well separated from residual protein and buffer signals and their intensities are expected to be most sensitive to the orientation of the ligand ring systems.

### Hydrogen/deuterium exchange mass spectrometry (HDX-MS)

HDX-MS experiments on HilD were conducted similar as described previously^39^. HDX-MS was performed on two samples of HilD, i.e., without or with C26 present. To do so, HilD (25 µM) was supplemented with 1% (v/v) of either DMSO or C26 (10 mM dissolved in DMSO) yielding a final concentration of 100 μM C26 in the sample. Both samples were stored in a cooled tray (1 °C) until measurement.

HDX reactions were prepared by a two-arm robotic autosampler (LEAP technologies) by addition of 67.5 μL HDX buffer (50 mM sodium phosphate pH 7.0, 200 mM NaCl, 1% (v/v) DMSO) prepared with 99.9% D_2_O to 7.5 μL of HilD sample without C26. HDX reactions of HilD in presence of C26 were prepared similarly but the HDX buffer was supplemented with 100 µM C26 to prevent dilution of the compound upon addition of HDX buffer. After incubation at 25 °C for 10, 30, 100, 1,000 or 10,000 seconds, 55 µL of the HDX reaction was withdrawn and added to 55 µL of pre-dispensed quench buffer (400 mM KH_2_PO_4_/H_3_PO_4_, pH 2.2, 2 M guanidine HCl) kept at 1 °C. 95 µL of the resulting mixture was injected into an ACQUITY UPLC M-Class System with HDX Technology (Waters)^83^. Non-deuterated protein samples were prepared similarly (incubation for approximately 10 s at 25 °C) by 10 fold dilution of HilD samples with H_2_O containing HDX buffer. The injected samples were flushed out of the loop (50 µL) with H_2_O + 0.1% (v/v) formic acid (100 µL min^−1^) and guided to a protease column (2 mm × 2 cm) containing the below specified proteases immobilized to the bead material, which was kept at 12 °C. For each protein state and timepoint, three replicates (individual HDX reactions) were digested with porcine pepsin, while another three replicates were digested with a column filled with a 1:1 mixture of protease type XVIII from *Rhizopus spp.* and protease type XIII from *Aspergillus saitoi*. In both cases, the resulting peptides were trapped on an AQUITY UPLC BEH C18 1.7 µm 2.1 × 5 mm VanGuard Pre-column (Waters) kept at 0.5 °C. After 3 min of digestion and trapping, the trap column was placed in line with an ACQUITY UPLC BEH C18 1.7 μm 1.0 × 100 mm column (Waters), and the peptides eluted at 0.5 °C using a gradient of eluents A (H_2_O + 0.1% (v/v) formic acid) and B (acetonitrile + 0.1% (v/v) formic acid) at a flow rate of 30 μl min^−1^ as follows: 0 7 min: 95 65% A; 7 8 min: 65 15% A; 8 10 min: 15% A; 10 11 min: 5% A; 11 16 min: 95% A. The eluted proteins were guided to a G2-Si HDMS mass spectrometer with ion mobility separation (Waters), and peptides ionized with an electrospray ionization source (250 °C capillary temperature, spray voltage 3.0 kV) and mass spectra acquired in positive ion mode over a range of 50 to 2,000 m/z in enhanced high definition MS (HDMS^E^) or high definition MS (HDMS) mode for non-deuterated and deuterated samples, respectively^84,85^. [Glu1]-Fibrinopeptide B standard (Waters) was employed for lock-mass correction. During separation of the peptide mixtures on the ACQUITY UPLC BEH C18 column, the protease column was washed three times with 80 µl of wash solution (0.5 M guanidine hydrochloride in 4% (v/v) acetonitrile), and blank injections performed between each sample to reduce peptide carry-over.

Peptide identification and analysis of deuterium incorporation were carried out with ProteinLynx Global SERVER (PLGS, Waters) and DynamX 3.0 softwares (Waters) as described previously^39^. In summary, peptides were identified with PLGS from the non-deuterated samples acquired with HDMS^E^ by employing low energy, elevated energy, and intensity thresholds of 300, 100 and 1,000 counts, respectively. Identified ions were matched to peptides with a database containing the amino acid sequence of HilD, porcine pepsin, and their reversed sequences with the following search parameters: peptide tolerance = automatic; fragment tolerance = automatic; min fragment ion matches per peptide = 1; min fragment ion matches per protein = 7; min peptide matches per protein = 3; maximum hits to return = 20; maximum protein mass = 250,000; primary digest reagent = non-specific; missed cleavages = 0; false discovery rate = 100. Only peptides that were identified in three out of six (for each protease digestion regime) non-deuterated samples and with a minimum intensity of 25,000 counts, a maximum length of 30 amino acids, a minimum number of three products with at least 0.1 product per amino acid, a maximum mass error of 25 ppm and retention time tolerance of 0.5 minutes were considered for further analysis. Deuterium incorporation into peptides was quantified with DynamX 3.0 software (Waters). Hereby, the datasets generated with pepsin digestion or after digestions with proteases type XIII and XVIII were pooled. All spectra were manually inspected and, if necessary, peptides omitted (e.g., in case of low signal-to-noise ratio or presence of overlapping peptides).

The observable maximal deuterium uptake of a peptide was calculated by the number of residues minus one (for the N-terminal residue) minus the number of proline residues contained in the peptide. For the calculation of HDX in per cent the absolute HDX was divided by the theoretical maximal deuterium uptake multiplied by 100. To render the residue specific HDX differences from overlapping peptides for any given residue of HilD, the shortest peptide covering this residue was employed. Where multiple peptides were of the shortest length, the peptide with the residue closest to the peptide’s C-terminus was utilized.

### Microscale Thermophoresis

MST measurements for the binding of HilD to HilE were performed as previously described^39^. In summary, EYFP-HilD (100 nM) was incubated with 100 μM of either oleic acid or C26 (with a final concentration of 1% (v/v) DMSO) for 10 minutes at room temperature (22–25 °C), and subsequently mixed 1:1 with varying concentrations of HilE. Samples were incubated together for 10 min at room temperature, centrifuged for 5 min and loaded to standard capillaries (NanoTemper Technologies GmbH, #MO-K022). MST runs were performed at 25 °C on a NanoTemper Monolith NT.115, with an excitation power of 60% and medium MST power. Data were analyzed using the MO.Affinity Analysis v2.3 software, and affinity constants were calculated using the *K*d model.

### Subcellular Quantification of Uptake

*S*. Typhimurium was grown in Mueller-Hinton-2 medium to an OD_600_ of 0.8 and incubated with the inhibitors (100 ng/mL, ∼250 nM) for 10 min. Cells were then subjected to a fractionation protocol as previously described^43^. The obtained fractions were protein-depleted via precipitation using a mixture of H_2_O/ACN/MeOH (40/30/30) and a centrifugation at 3000 rpm in cold environment (4°C). The supernatant was evaporated in a CentriVap (Labconco, Kansas, MO, USA) device over night at 30°C before resuspending in 50 µL of appropriate LC/MS/MS buffer, containing 10 ng/mL caffeine as internal standard. Results were generated on a triple quadrupole mass spectrometer (AB Sciex 6500, Darmstadt, Germany) connected to an Agilent 1290 Infinty II UHPLC (Agilent Technologies, Santa Clara, CA, USA). Separation was done via reverse phase with an RP-18 column (Phenomenex Gemini, 3µm NX-C18 110A, 50 × 2 mm) with a respective column guard (5 × 2 mm, Phenomenex, Torrance, CA, USA) at a flowrate of 700 µL/min and an elution gradient from 5% to 95% B within 4 min (A: H_2_O+0.1% HCOOH; B: ACN+0.1% HCOOH). Source parameters of the mass spectrometer and mass transitions are given in Supplementary Tab. 7. Calibration curves were recorded with the compounds in the respective matrices. Data was quantified with Multiquant 3.03 (AB Sciex, Darmstadt, Germany).

### Statistics

Statistical analyses were conducted using GraphPad Prism 10.1.1. Data are presented as mean ± s.d. Comparisons with *p* > 0.05 were not considered significant.

## Data availability

Supplementary figures, data collection, and further supporting information are available free of charge at the publisheŕs website. All molecular dynamics trajectories and raw data related to the protein-ligand interactions within the simulations will be available in the repository: DOI: 10.5281/zenodo.8129269, 10.5281/zenodo.8139104 and 10.5281/zenodo.10993310 upon publication.

## Acknowledgments

A.B. and S.W. acknowledge funding from the German Center for Infection Research (DZIF, TTU06.912) and the Baden-Württemberg Stiftung (BWST_WSF-018).

T.K. acknowledges funding by the Clusters of Excellence EXC2180 iFIT (project ID 390900677), and EXC2124 CMFI (project ID 390838134), the Faculty of Medicine of the University of Tübingen’s Fortüne program (NR.2613-0), the Federal Ministry of Education and Research (BMBF), the Baden-Württemberg Ministry of Science as part of the Excellence Strategy of the German Federal and State Governments, by the means of the program TüCAD2, as well as the German Center for Infection Research (DZIF, TTU06.716).

V.S.K. and M. B. acknowledge funding from the Baden-Württemberg Stiftung (BWST_WSF-018) and from the German Center for Infection Research (DZIF, TTU 09.722, TTU06.801).

W.S. and G.B. acknowledge support by the German Research Council (DFG) through the core facility for HDX-MS (project 324652314 to Gert Bange, Marburg).

A.K. and J-C. S. acknowledge DZIF funding TI07.003.

M.P and A.F. acknowledge funding by the Federal Ministry of Health of Germany for integrated genomic surveillance of enteric pathogens (grant D81959). We thank the Sequencing Core Facility of the Genome Competence Centre, Robert Koch Institute, for providing excellent sequencing services.

A.P. and T.K. acknowledge CSC-Finland for generous computational resources.

M.D.H. acknowledges support from the Baden-Württemberg Stiftung (BWST_WSF-018) and from institutional funds of the Max Planck Society.

S. W. acknowledges funding from the and from the German Center for Infection Research (DZIF, TTU06.801, TTU06.808, TTU06.819 and TTU06.829). Work in the laboratory of S.W. was also supported by infrastructural measures of the Cluster of Excellence EXC2124 Controlling Microbes to Fight Infections (CMFI), project ID 390838134.

We thank Andrea Eipper, Melanie Nowak, Antje Ritter, and Nick Mozer for technical assistance. We would like to thank Dr. Libera Lo Presti for her substantial contribution in reviewing and improving the manuscript. We are grateful to the diagnostic team of the Institute of Medical Microbiology and Hygiene of Tübingen for providing clinical isolates. We thank Dr. Thomas Hesterkamp for his valuable advice on the drug development process.

## Author contributions

A.B. designed and performed experiments on mode of action, spectrum of activity, wrote the manuscript with input from all authors, acquired funding

J.D.J. designed and performed protein expression and purification, and *in vitro* experiments on mode of action

I.G. designed and performed experiments on the phenotypic screen, transcriptome analysis, and performed *in-silico* analysis of point mutants

T.K. performed the *in silico* structural analysis of the HilD-C26 complex.

V.S.K. performed the synthesis of C26 and analogs

W.S. performed HDX-MS experiments

A.K. performed experiments on T1SS

S.S. performed experiments on T3SS-2

J-C.S. performed experiments on alanine scan

M.P. designed, supervised and performed subtyping, sequence analysis and selection of representative strains of German clinical isolates

A.N. performed the synthesis of compound analogs

S.K. performed the synthesis of compound analogs

S-K.H. performed the subcellular quantification of uptake

T.C. performed the synthesis of compound analogs

C.P. performed protein expression and purification, and *in vitro* experiments

M.C. designed, performed and evaluated NMR experiments

K.R. developed formulation for the animal studies for C26

M.M. selected clinical isolates from the university hospital of Tübingen

G.B. supervised the HDX-MS experiments, acquired funding

A.F. designed, supervised selection of representative strains of German clinical isolates

A.P. conceptualized and executed the initial virtual screening and supervised the *in-silico* data analyses, acquired resources

M.B. conceptualized and supervised the chemistry part, acquired funding

M.D.H. conceptualized and supervised *in vitro* experiments, acquired funding

S.W. conceptualized and supervised the project, acquired funding

All authors provided critical feedback and helped shape the research, analysis, and manuscript.

## Extended data figures

**Figure 1.**
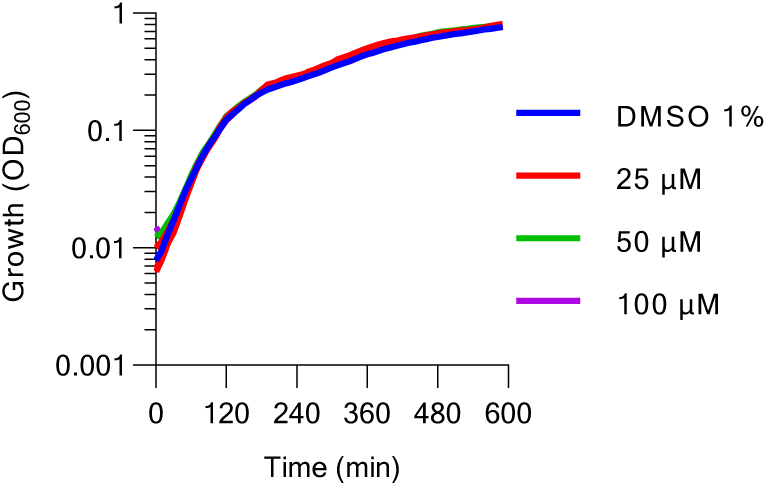
Growth of *S.* Typhimurium SL1344 in LB medium supplemented with C26 at different concentrations or 1% (v/v) DMSO. Experiment performed in a 96-well plate. Growth assessed by measuring the OD at 600 nm (n = 3 biological replicates).

**Figure 2.**
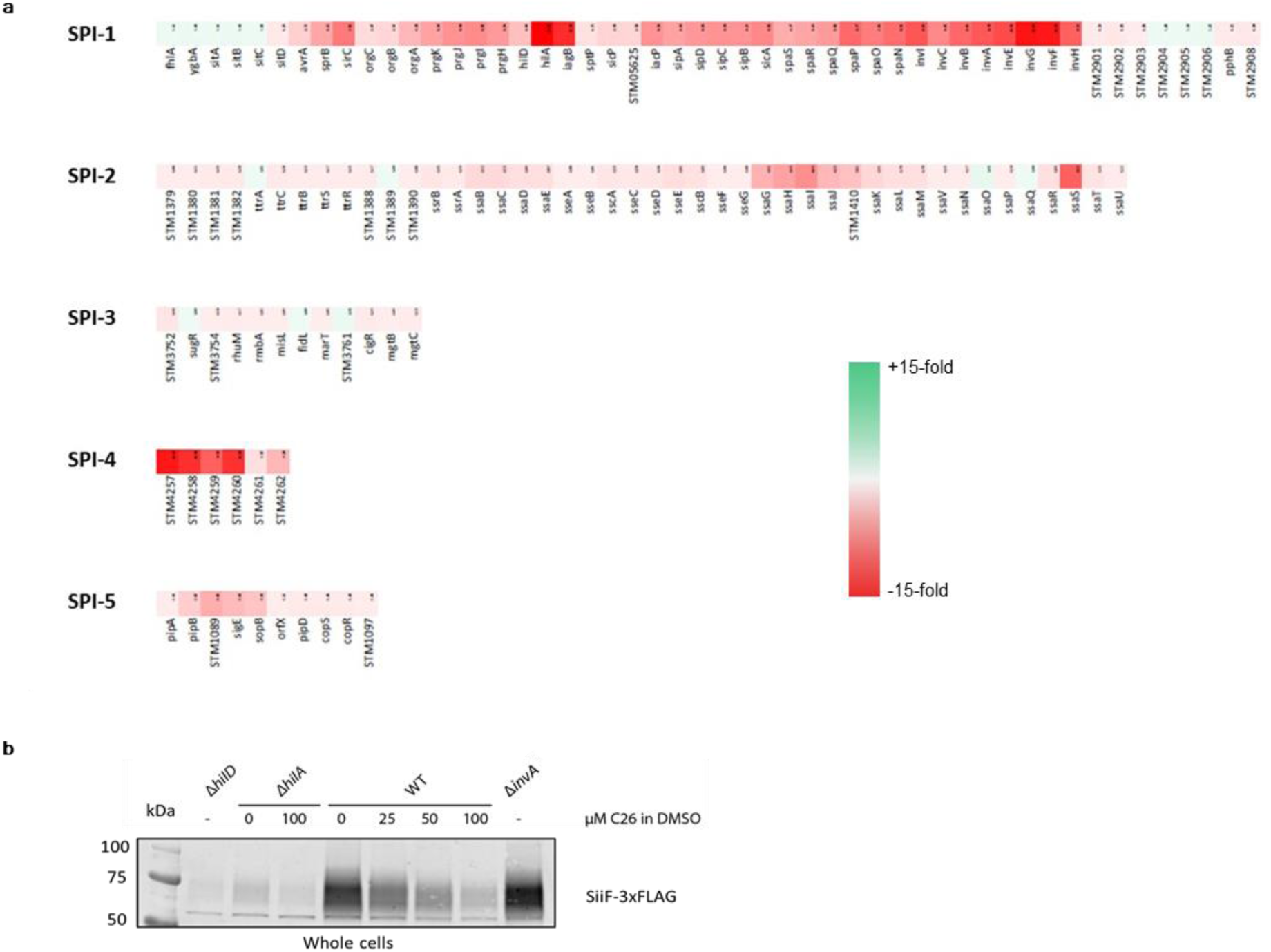
Effect of C26 on the expression of invasion genes. **a)** Heat map of mRNA fold-changes from genes encoded in SPI-1-5. **b)** Monitoring the expression of FLAG-tagged T1SS structure protein SiiF by Western blotting using mouse anti-FLAG (1:10000) antibody in the indicated *Salmonella* strains.

**Figure 3.**
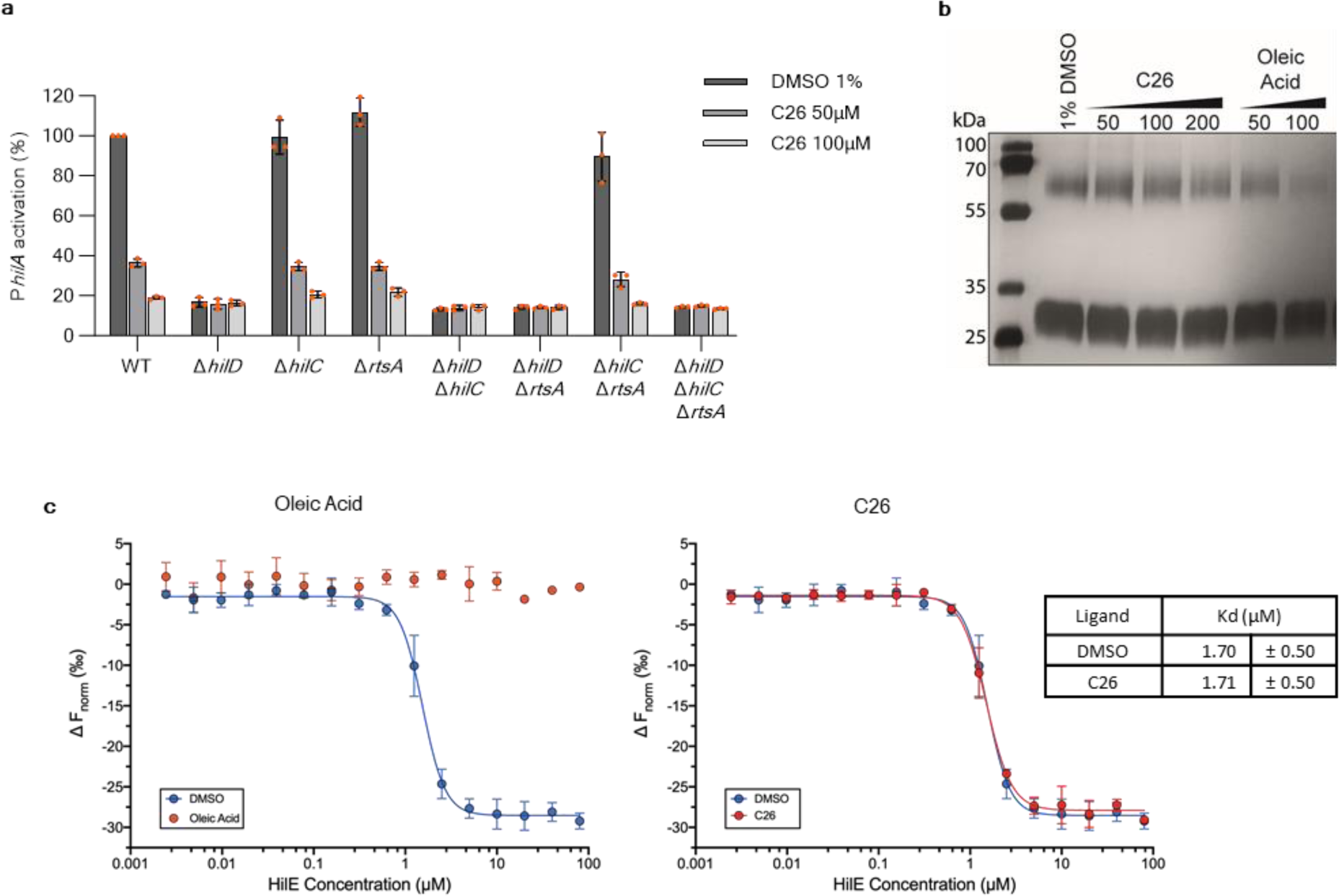
Effect of C26 on HilD dimerization and activity. **a)** Cell-based P*hilA* activation assay to assess the effect of C26 on HilD, HilC, and RtsA transcriptional activity levels (n = 3 biological replicates). **b)** HilD dimerization investigated by BS^3^ cross-linking of HilD monomers (10 µM) in the presence of C26 or oleic acid. **c)** MST assay to investigate the effect of C26 and oleic acid on the *in vitro* formation of the HilD-HilE heterodimer. EYFP-HilD was incubated with either 1% DMSO (blue), 100 μM oleic acid (orange) or 100 μM C26 (red) and increasing concentrations of HilE. Data show changes in thermophoresis at an MST on-time of 1.5 s and represents the mean ± SD of four replicates.

**Figure 4.**
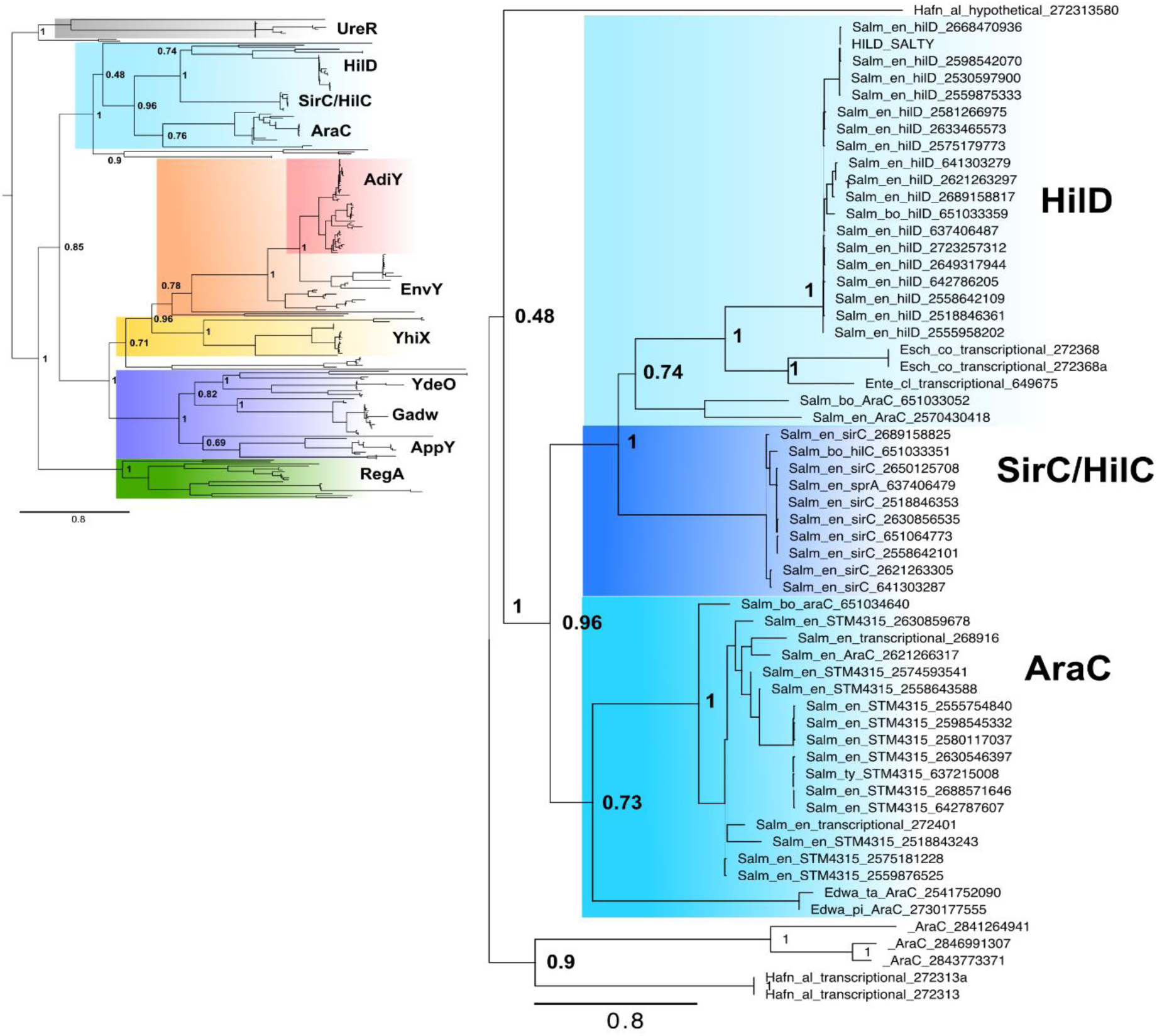
Dendrogram showing homologs of HilD. The tree was generated with FigTree (v1.4.4). Left, overview of all members of AraC/XylS family protein grouped. Right: subclade containing HilD, HilC and AraC from *Salmonella* spp. highlighted in blue. Only clades with branch statistical support of posterior probability values (calculated using aBayes) with values >0.7 or relevant for discussion are displayed.

**Figure 5.**
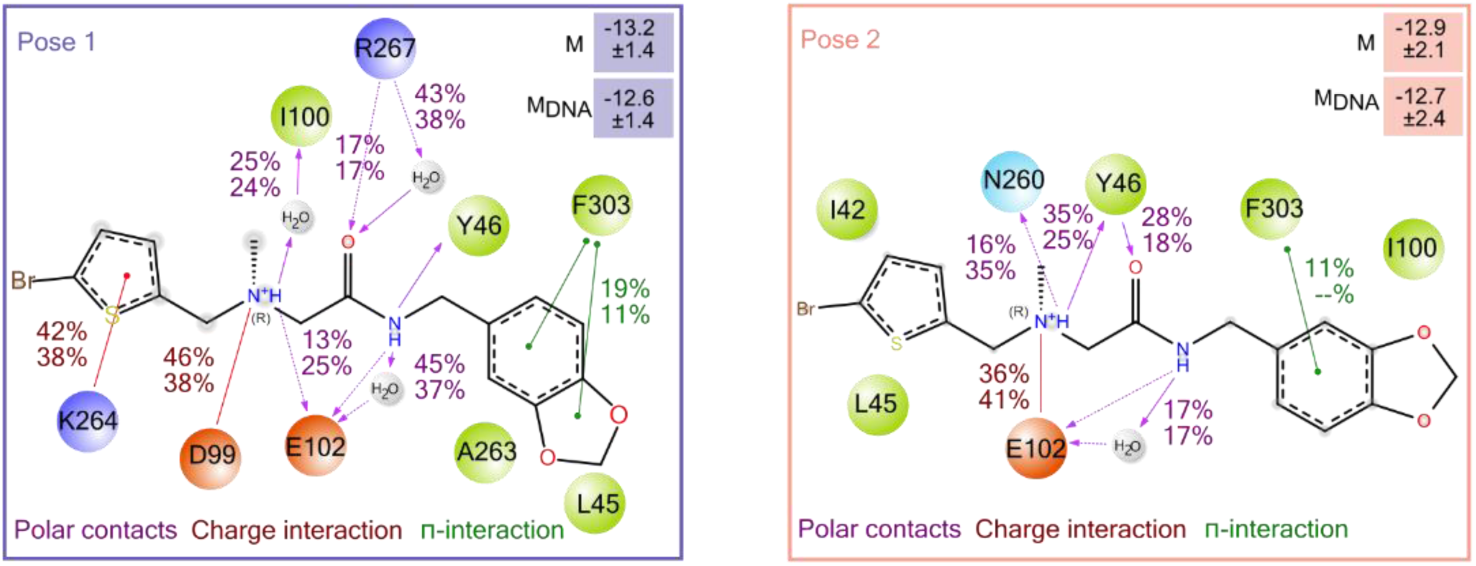
2D schematic representation of C26 in pose 1 and pose 2 potential binding modes summarizing their interaction frequency along the analyzed trajectory of MD simulations (∼10 µs per ligand). Interaction frequency (%) in the upper labels are derived from monomers + DNA while the below numbers derive from simple monomeric simulations. Polar interactions are depicted in purple, charged interactions in red and π-mediated interactions as green lines. Quantification of the predicted binding energy for each ligand along the simulated trajectory, using MM/GBSA (see extended methods for calculation). The median of the calculated energies is displayed as colored boxes, together with its standard deviation, and free energy binding calculation (kcal/mol normalized by the Heavy Atoms Count, HAC).

**Figure 6.**
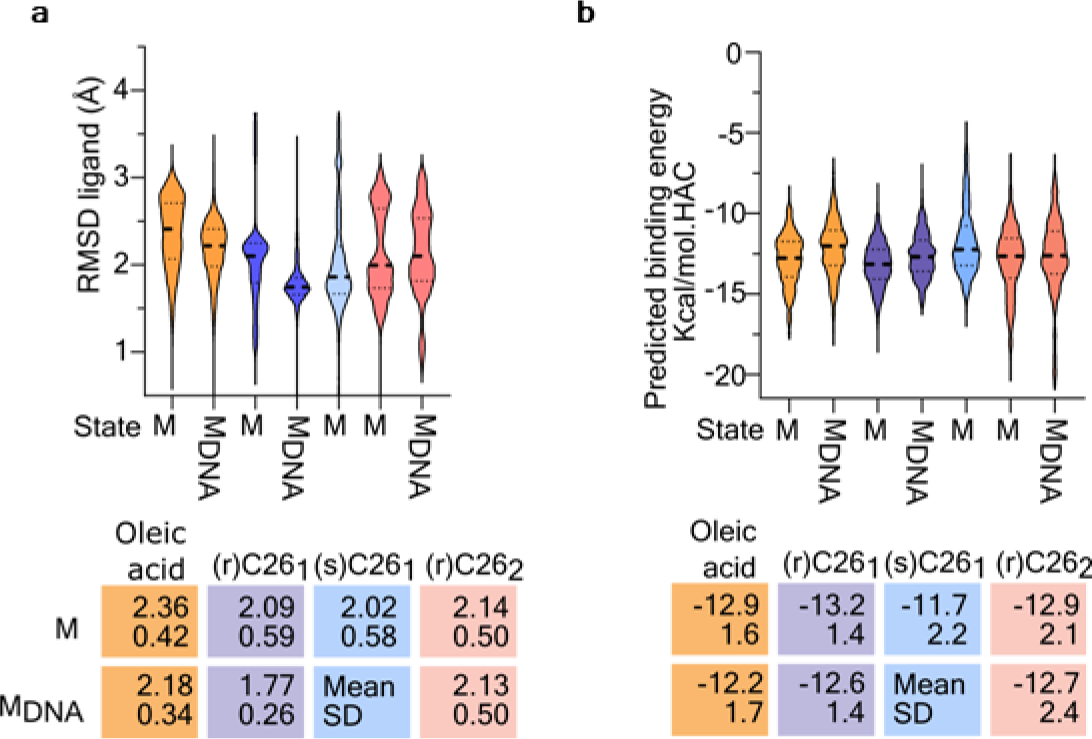
Violin plots depicting the **a)** Root mean square deviation of the ligand’s heavy atoms along the trajectory time (∼10 µs per ligand). **b)** Quantification of the predicted binding energy for each ligand along the simulated trajectory, using MM/GBSA (see extended methods for calculation), in the free energy binding calculations (kcal/mol normalized by the Heavy Atoms Count, HAC).

**Figure 7.**
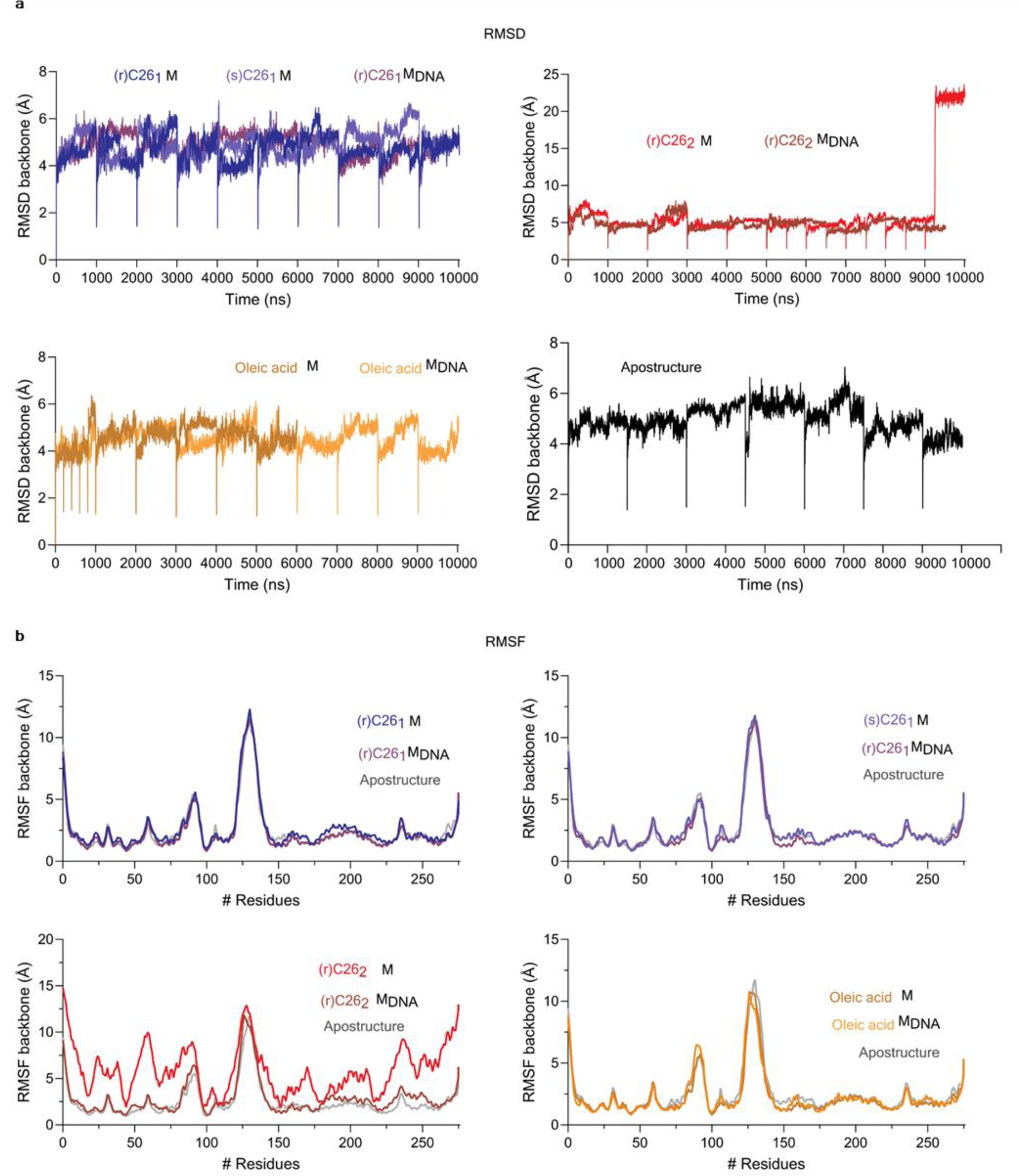
Protein’s backbone RMSD **(a)** and RMSF **(b)** were used to evaluate the equilibration of the system.

**Figure 8.**
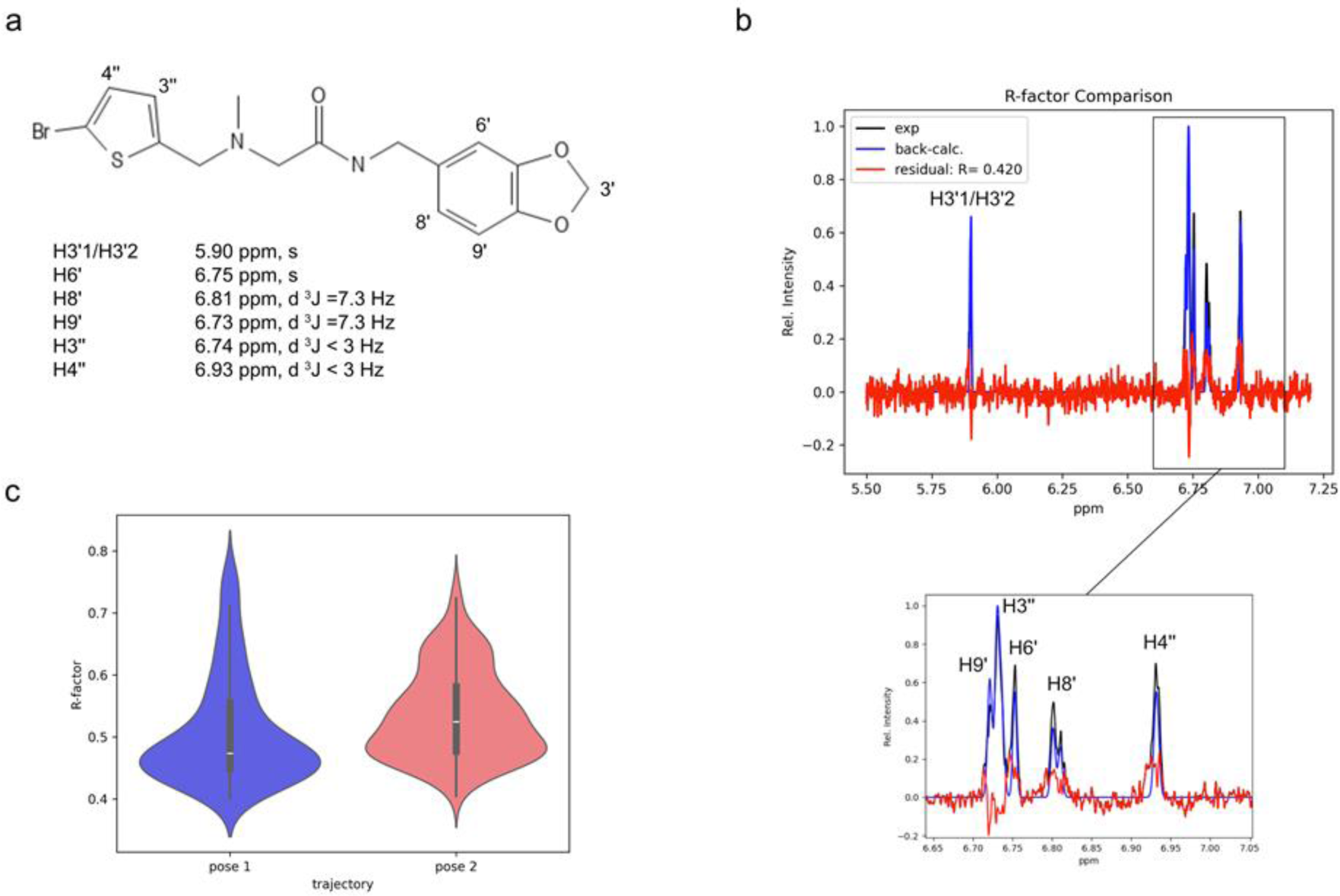
NMR saturation transfer difference (STD) experiments favour pose 1 over pose 2. **a)** NMR assignment of C26 protons considered in the STD analysis (s; singlet, d; doublet). **b)** Example of back-calculation of expected STD intensities using the CORCEMA algorithm. The program takes ligand affinity, protein and ligand concentrations and parameters of the NMR experiment into account to predict the relative intensities of STD peaks. In the current implementation, a simulated spectrum is also calculated based on ligand chemical shifts and couplings and an R-factor calculated on basis of RMSD to the experimental spectrum. Only the region shown was included in quantitative analysis. The experimental data is shown in black, while the back-calculated and residual are in blue and red, respectively. The inset shows the aromatic signals; note that H9’ and H3’’ are partially overlapped. The comparison shown is for the best frame from the two MD trajectories; frame 8 of the pose 1 trajectory (R-factor 0.420). **c)** Violin plots for the distribution of R-factors in MD trajectories starting from pose 1 and pose 2. The two distributions are significantly different (Mann-Whitney p-value < 1e-06), with the pose 1 trajectory enriched in frames with lower R-factors. All data shown is for a saturation time of 800 ms, representing a compromise between STD intensity and sensitivity to the ligand pose.

**Figure 9.**
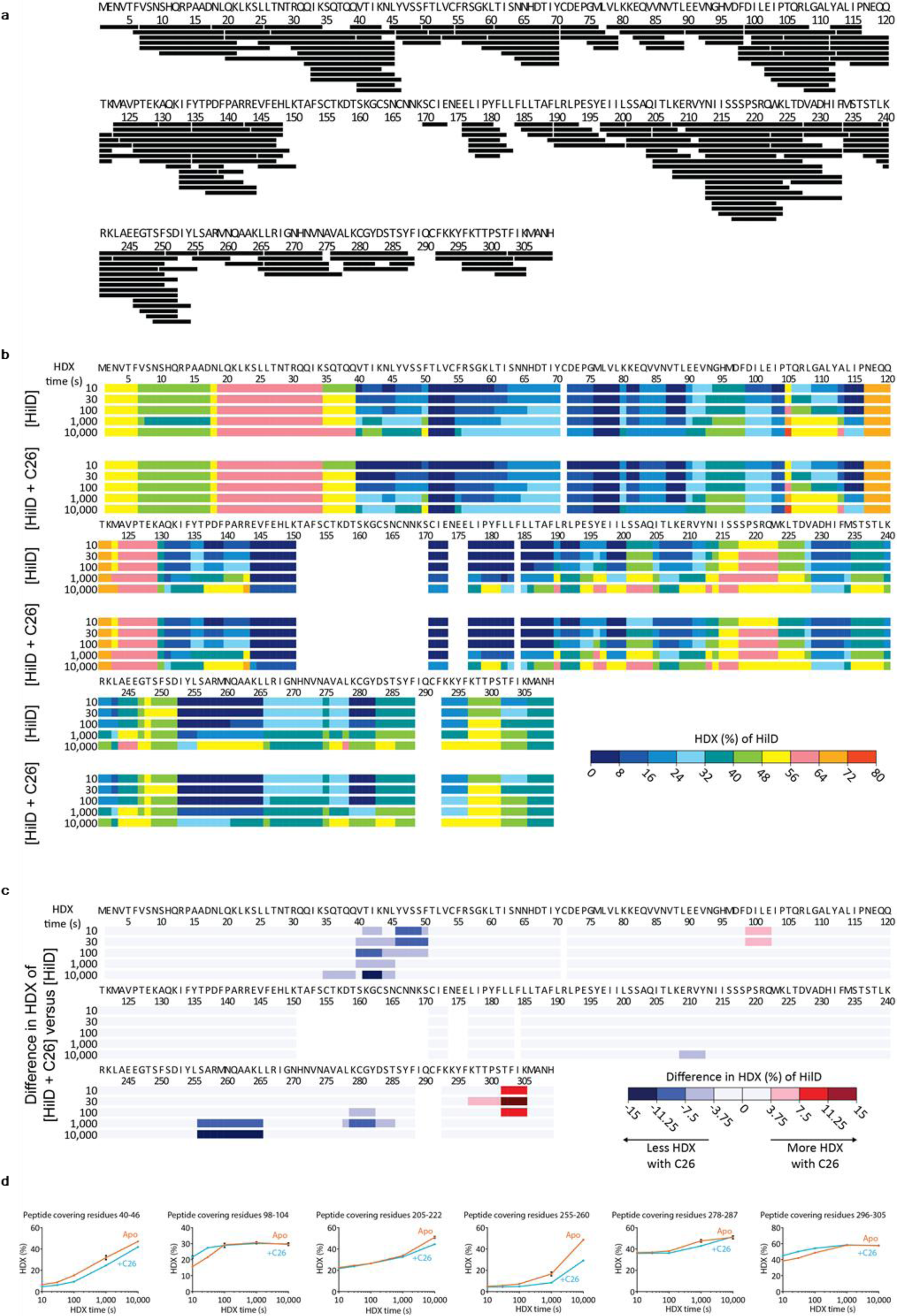
HDX-MS of HilD. **a)** Each black bar denotes a peptide of HilD identified during HDX-MS experiments. **b)** Residue-specific HDX for HilD, either in isolation or in presence of 100 µM C26, was obtained from peptides by employing the shortest peptide covering any residue. No HDX could be obtained for amino acid sequences in the gaps, which indicate regions not covered by any peptides. **c)** C26-dependent changes in HDX of HilD, expressed as the difference in residue-specific HDX between HilD in isolation (HilD) and in the presence of 100 µM C26 (HilD + C26). **d)** HDX of selected representative HilD peptides. Data represent mean ± s.d. of n = 3 technical replicates (individual HDX reactions).

**Figure 10.**
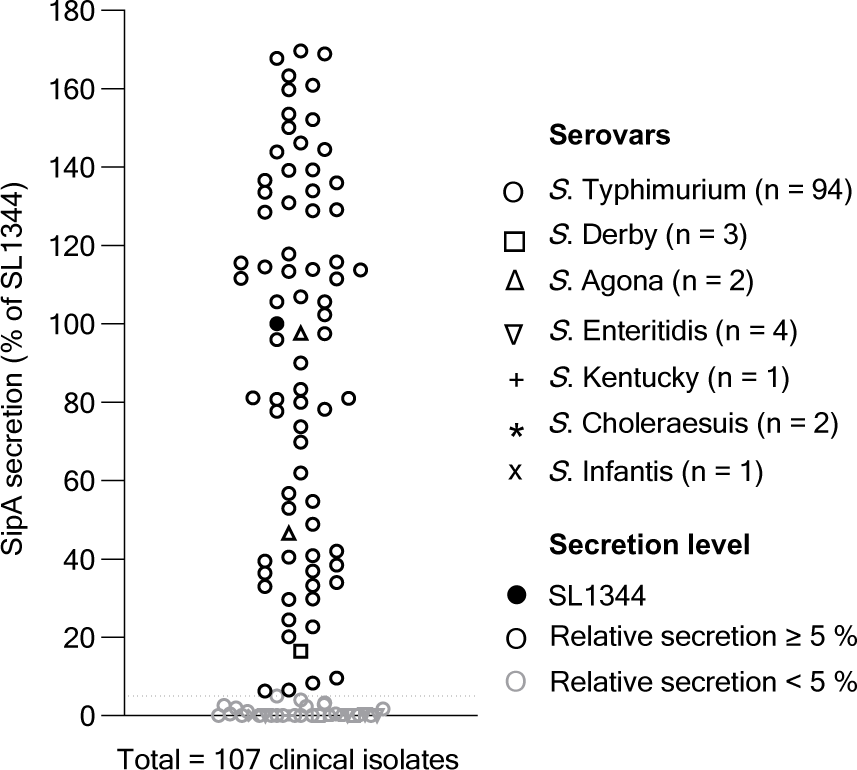
Secretion levels of SipA in clinical isolates of *S. enterica* relative to SL1344.

## Supplementary information

**Figure 1.**
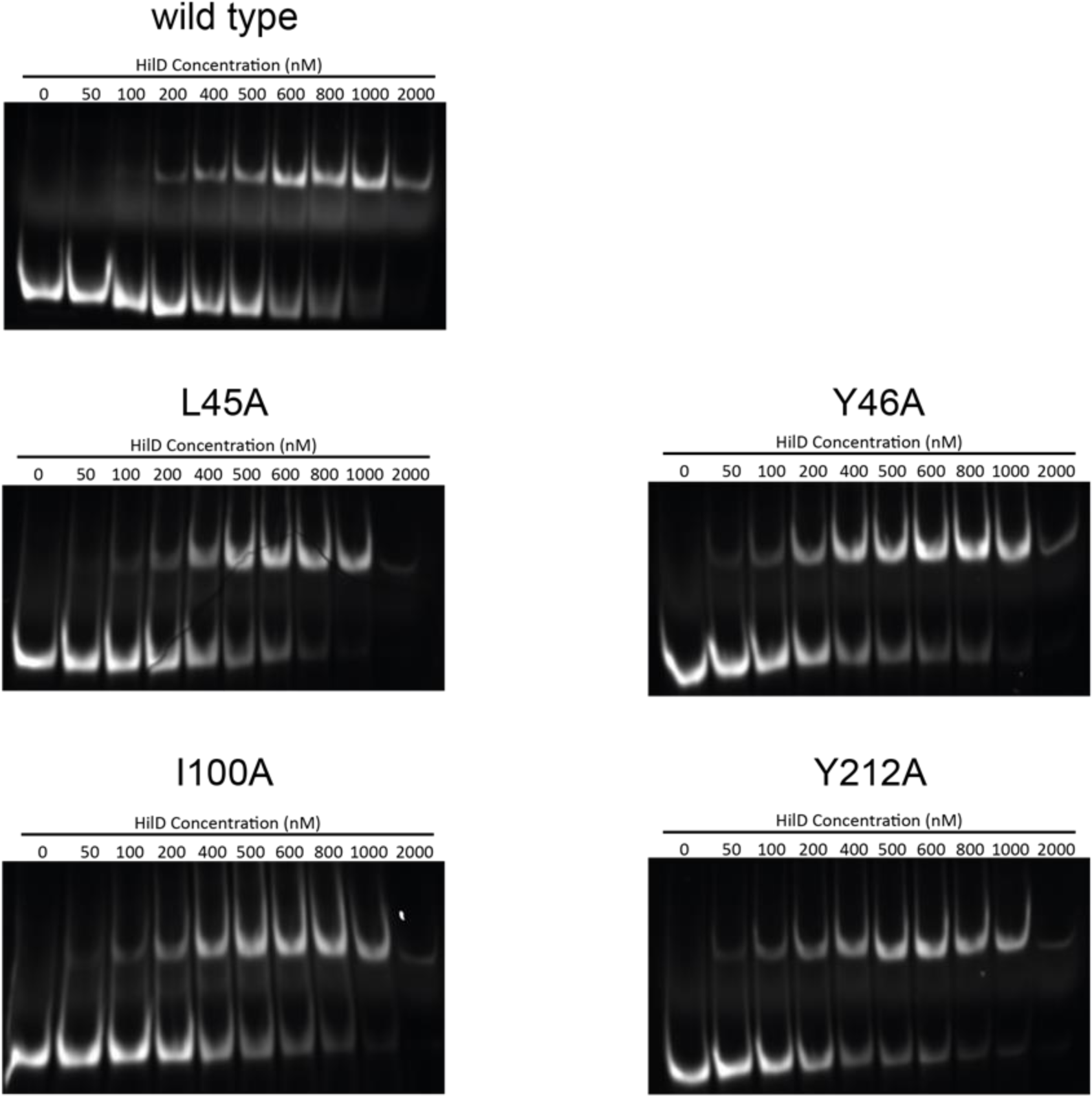
Electrophoretic mobility shift assay (EMSA) showing the binding of purified HilD mutants to the promoter of *hilA*.

**Figure 2.**
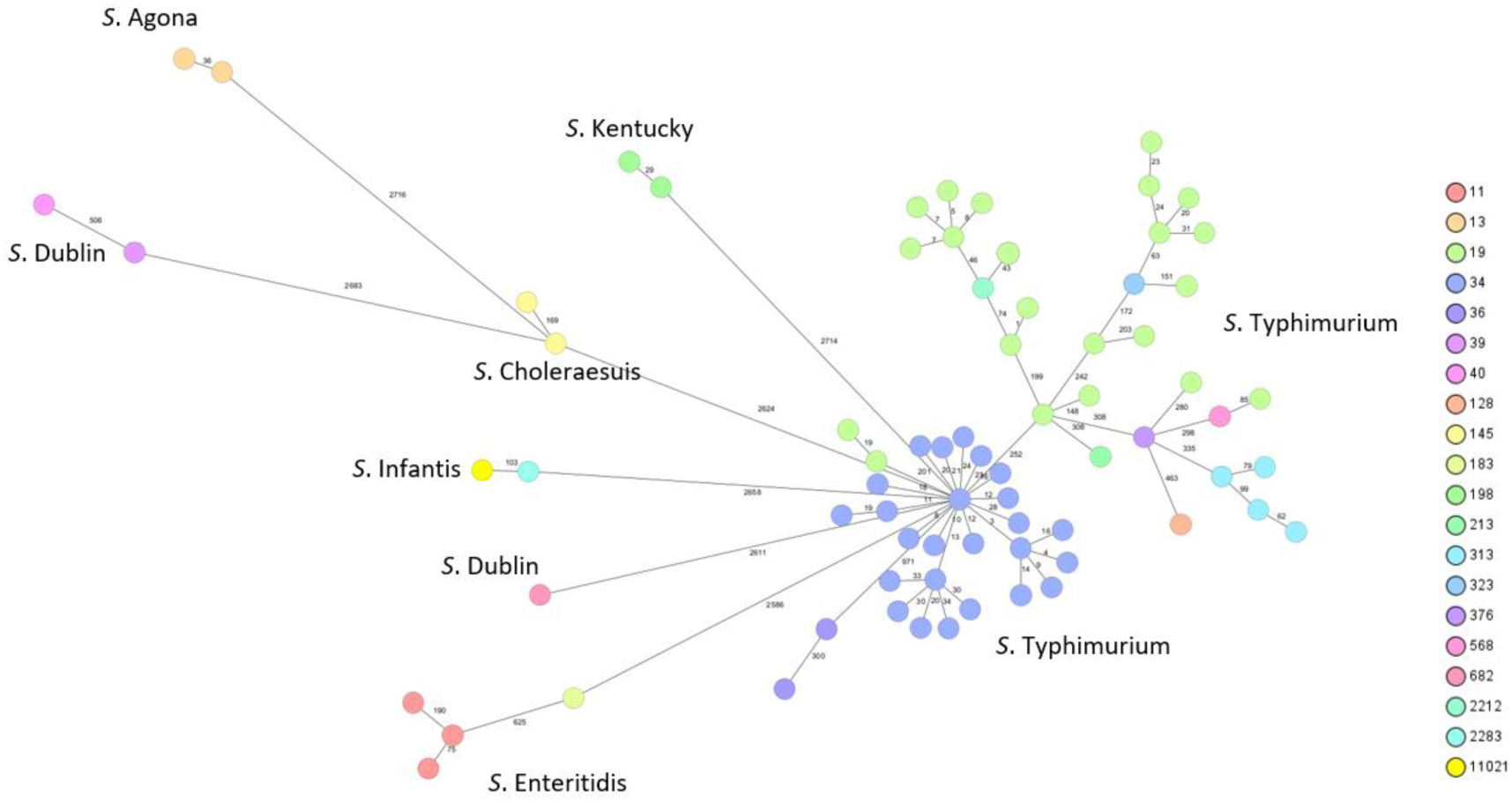
Minimum spanning tree of 75 selected S. enterica strains from Germany based on cgMLST (Ridom SeqSphere+, 3.002 alleles Enterobase cgMLST scheme). Color-coding based on sequence type.

**Table 1.** Chemical properties of compound C26. Values were generated with ChemAxon and DataWarrior.

| MW (g/mol) | clogP | clogS | HBD | HBA | pI | Rotatable bonds | Druglikeness |
| --- | --- | --- | --- | --- | --- | --- | --- |
| 397 | 2.69 | -3.79 | 1 | 5 | 10.4 | 6 | 2.68 |

Molecular weight (MW), H-bond donor (HBD) and acceptor (HBA), isoelectric point (pI).

**Table 2.** Recorded death events of mice during the first 72 hours after treatment.

| Compound | Route | Dose (mg/kg) | Toxicity (death / test) |  |  |  |  |
| --- | --- | --- | --- | --- | --- | --- | --- |
|  |  |  | 1 h | 2 h | 24 h | 48 h | 72 h |
| Vehicle | PO | 5 ml/kg | 0/3 | 0/3 | 0/3 | 0/3 | 0/3 |
| C26 | PO | 3 | 0/3 | 0/3 | 0/3 | 0/3 | 0/3 |
|  |  | 10 | 0/3 | 0/3 | 0/3 | 0/3 | 0/3 |
|  |  | 30 | 0/3 | 0/3 | 0/3 | 0/3 | 0/3 |

PO, *per os*. Vehicle: 5% Tween 20, lecithin (from soy bean: 40 mg in 800 μL distilled water), 45 % PBS.

**Table 3.**
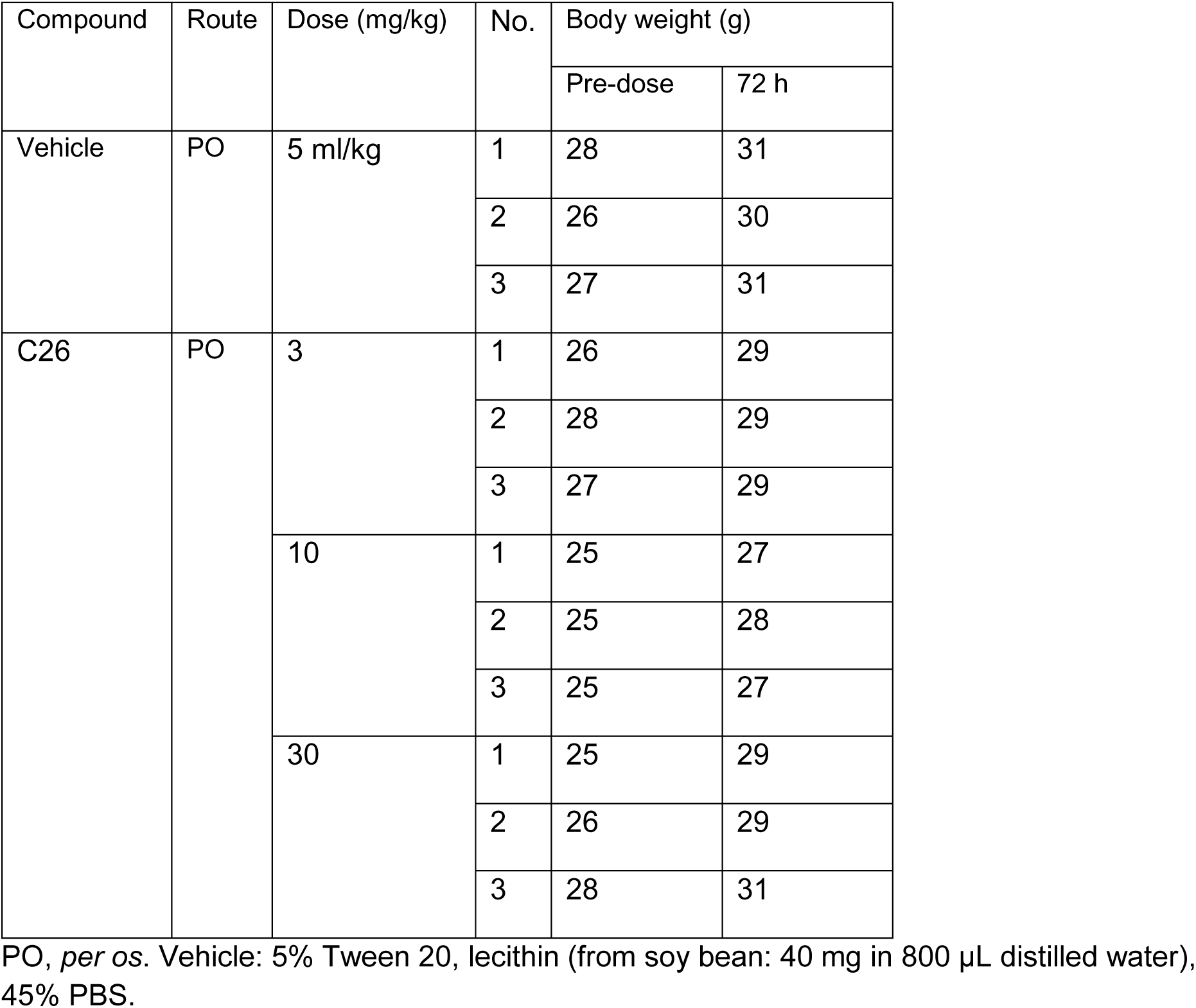
Recorded body weights of mice before treatment and at 72 hours after treatment. PO, *per os*. Vehicle: 5% Tween 20, lecithin (from soy bean: 40 mg in 800 μL distilled water), 45% PBS.

**Table 4.** Molecular mass determination by SEC-MALS analysis of HilD in the presence of DMSO (1%) or C26 (100 µM). Values correspond to the curves shown in Fig. 3h.

| Construct | Molecular Mass (kDa) |  |  |
| --- | --- | --- | --- |
| | Theoretical (Dimer) | Measured (± SD) | |
| HiID | 70.4 | 70.4 | ± 2.0 |
| HiID + DMSO | 70.4 | 71.4 | ± 2.8 |
| HiID + C26 | 70.4 | 69.8 | ± 1.8 |

**Table 5.** List of the 50 most frequent substitutions found among 2,351 sequences of *S. enterica*.

| Substitution | # | Frequency (10<sup>-3</sup>) |
| --- | --- | --- |
| N86S | 608 | 258,6134 |
| R30Q | 259 | 110,1659 |
| A124S | 257 | 109,3152 |
| S220G | 238 | 101,2335 |
| V40I | 227 | 96,55466 |
| A275S | 219 | 93,15185 |
| A110D | 217 | 92,30115 |
| D72E | 182 | 77,41387 |
| K169Q | 142 | 60,39983 |
| A130V | 126 | 53,59422 |
| V228I | 122 | 51,89281 |
| T226P | 115 | 48,91536 |
| V125I | 108 | 45,9379 |
| T127V | 108 | 45,9379 |
| S164N | 103 | 43,81114 |
| N165S | 102 | 43,38579 |
| G162S | 100 | 42,53509 |
| V125K | 97 | 41,25904 |
| P137S | 96 | 40,83369 |
| K157M | 92 | 39,13228 |
| D230E | 90 | 38,28158 |
| M123R | 80 | 34,02807 |
| K169R | 80 | 34,02807 |
| T121I | 79 | 33,60272 |
| I215L | 77 | 32,75202 |
| F134I | 71 | 30,19991 |
| R221K | 68 | 28,92386 |
| E209K | 58 | 24,67035 |
| E73A | 55 | 23,3943 |
| M306I | 54 | 22,96895 |
| T37A | 49 | 20,84219 |
| K305F | 48 | 20,41684 |
| A307K | 47 | 19,99149 |
| N308M | 45 | 19,14079 |
| S249N | 44 | 18,71544 |
| H309A | 43 | 18,29009 |
| K305R | 35 | 14,88728 |
| A124V | 34 | 14,46193 |
| K279T | 33 | 14,03658 |
| H231N | 30 | 12,76053 |
| T127A | 29 | 12,33518 |
| F134L | 25 | 10,63377 |
| M96L | 24 | 10,20842 |
| S237P | 22 | 9,35772 |
| T41S | 21 | 8,932369 |
| K43R | 21 | 8,932369 |
| K150R | 19 | 8,081667 |
| E128D | 18 | 7,656316 |
| S216Y | 18 | 7,656316 |
| I62V | 16 | 6,805615 |

**Table 6.** Clinical isolates of *S. enterica*. UKT, University hospital of Tübingen. RKI, Robert Koch Institute. Antibiotic susceptibilities were investigated by broth microdilution according to EUCAST criteria (The European Committee on Antimicrobial Susceptibility Testing. Breakpoint tables for interpretation of MICs and zone diameters. Version 10.0, 2020. http://www.eucast.org) AMP, Ampicillin. MEZ, Mezlocillin. OTE, Oxytetracycline. SMZ, Sulfamethoxazole. STR, Streptomycin. CMP, Chloramphenicol. CTM, Cefotiam. CTX, Cefotaxime. MSU, Mezlocillin/Sulbactam. NAL, Nalidixic acid. SXT, Trimethoprim / Sulfamethoxazole. TCY, Tetracycline. AZM, Azithromycin. CAZ, Ceftazidime. CIP, Ciprofloxacin. FOX, Cefoxitin. GEN, Gentamicin. Table cells are left blank when the corresponding characteristic is unknown. *Identified using NCBI AMRFinderPlus^86^.

| Name | Source | Resistance phenotype | Resistance genes* | ST | Provider |
| --- | --- | --- | --- | --- | --- |
| **S. Typhimurium** |  |  |  |  |  |
| 67708850 | Blood |  |  |  | UKT |
| 67529692 | Stool |  |  |  | UKT |
| 67962944 | Stool |  |  |  | UKT |
| 67775928 | Stool |  |  |  | UKT |
| 67774011 | Stool |  |  |  | UKT |
| 67738336 | Stool |  |  |  | UKT |
| 67713042 | Urine |  |  |  | UKT |
| 67708976 | Stool |  |  |  | UKT |
| 67681882 | Stool |  |  |  | UKT |
| 67625108 | Stool |  |  |  | UKT |
| 67564932 | Stool |  |  |  | UKT |
| 67547889 | Stool |  |  |  | UKT |
| 67463497 | Stool |  |  |  | UKT |
| 67459438 | Stool |  |  |  | UKT |
| 67453336 | Urine |  |  |  | UKT |
| 67447090 | Stool |  |  |  | UKT |
| 67444809 | Stool |  |  |  | UKT |
| 67411073 | Stool |  |  |  | UKT |
| 67189086 | Stool |  |  |  | UKT |
| 67187434 | Stool |  |  |  | UKT |
| 67178294 | Stool |  |  |  | UKT |
| 67065118 | Stool |  |  |  | UKT |
| 67064696 | Stool |  |  |  | UKT |
| 63957635 | Blood |  |  |  | UKT |
| 63852331 | Blood |  |  |  | UKT |
| 63804211 | Blood |  |  |  | UKT |
| SL3 | Stool |  |  |  | UKT |
| SL17 | Stool |  |  |  | UKT |
| SL18 | Stool |  |  |  | UKT |
| SL60 | Stool |  |  |  | UKT |
| SL131 | Stool |  |  |  | UKT |
| SL120 | Stool |  |  |  | UKT |
| SL256 | Stool |  |  |  | UKT |
| SL175 | Stool |  |  |  | UKT |
| SL145 | Stool |  |  |  | UKT |
| 06-01900 | Stool | AMP, MEZ, OTE, SMZ, STR | *bla*<sub>TEM-1</sub>, *sul2*, *tet*(B) | 34 | RKI |
| 18-02653 |  | AMP, CMP, CTM, CTX, MEZ, MSU, NAL, OTE, STR | *bla*<sub>CARB-2</sub>, *bla*<sub>CTX-M-1</sub>, *floR*, *tet*(G) | 19 | RKI |
| 19-00422 | Stool | AMP, CTM, CTX, MEZ, MSU, OTE | *bla*<sub>TEM-1</sub>, *bla*<sub>CTX-M-1</sub>, *qnrS1*, *sul2*, *tet*(A) / *tet*(M) | 19 | RKI |
| 19-02162 | Blood | AMP, CAZ, FOX, CTM, CTX, MEZ, MSU, OTE, STR | *bla*<sub>TEM-1</sub>, *bla*<sub>CMY-2</sub>, *qnrB19*, *sul2*, *tet*(B) | 34 | RKI |
| 19-04442 | Stool | AMP, CTM, CTX, MEZ, MSU, STR, SXT | *bla*<sub>CTX-M-1</sub>, *sul2*, *dfrA17* | 34 | RKI |
| 19-05051 | Stool | AMP, CTM, CTX, MEZ, OTE, STR, SXT | *bla*<sub>CTX-M-1</sub>, *sul1*, *tet*(A), *dfrA1* | 19 | RKI |
| 20-00098 | Stool | AMP, CMP, MEZ, NAL, OTE, SXT | *bla*<sub>TEM-1</sub>, *qnrB19*, *cmlA1*, *sul3*, *tet*(B), *dfrA12* | 34 | RKI |
| 20-00760 |  | AMP, CTM, CTX, MEZ, MSU, STR, SXT | *bla*<sub>TEM-1</sub>, *bla*<sub>CTX-M-1</sub>, *qnrS1*, *sul1* / *sul2*, *dfrA12* | 34 | RKI |
| 20-01187 | Stool | AMP, MEZ, OTE, STR | *bla*<sub>TEM-1</sub>, *sul2*, *tet*(B) | 34 | RKI |
| 20-01384 | Stool | AMP, CMP, MEZ, MSU, STR | *bla*<sub>CARB-2</sub>, *floR*, *tet*(G) | 19 | RKI |
| 20-02017 | Stool | AMP, MEZ, OTE, STR | *bla*<sub>TEM-1</sub>, *sul2*, *tet*(B) | 34 | RKI |
| 20-02167 | Stool | AMP, MEZ, MSU, OTE, STR, SXT | *bla*<sub>TEM-1</sub>, *sul1* / *sul2*, *tet*(A), *dfrA1* | 34 | RKI |
| 20-02297 | Stool | Sensitive | None | 313 | RKI |
| 20-02498 | Stool | AMP, MEZ, STR | *bla*<sub>TEM-1</sub>, *sul2*, *tet*(B) | 34 | RKI |
| 20-02749 | Stool | AMP, CMP, MEZ, MSU, OTE, STR | *bla*<sub>TEM-1</sub>, *floR*, *sul2*, *tet*(B) | 34 | RKI |
| 20-03136 |  | Sensitive | None | 323 | RKI |
| 20-03498 |  | AMP, CMP, MEZ, OTE, SXT | *bla*<sub>TEM-1</sub>, *qnrS1*, *cmlA1* / *floR*, *sul2* / *sul3*, *tet*(A) / *tet*(M), *dfrA12* | 19 | RKI |
| 20-03595 |  | AMP, MEZ, MSU | *bla*<sub>TEM-1</sub> | 19 | RKI |
| 20-06170 |  | AMP, TCY | *bla*<sub>TEM-1</sub>, *sul2*, *tet*(B) | 34 | RKI |
| 20-06645 |  | Sensitive | None | 19 | RKI |
| 21-00373 | Stool | Sensitive | None | 19 | RKI |
| 21-00797 | Stool | AMP, AZM, CAZ, CTX, SXT, TCY, TMP | *bla*<sub>TEM-1</sub>, *bla*<sub>CTX-M-1</sub>, *tet*(B), *dfrA17* | 34 | RKI |
| 21-01183 | Stool | AMP, CIP, NAL, TCY | *bla*<sub>TEM-1</sub>, *qnrB19*, *sul2*, *tet(B)* | 34 | RKI |
| 21-01586 | Stool | AMP, CIP, CMP, NAL, TCY | *bla*<sub>CARB-2</sub>, *gyrA_D87N*, *floR*, *tet(G)* | 19 | RKI |
| 21-02490 | Stool | AMP, TCY | *bla*<sub>TEM-1</sub>, *sul2*, *tet(B)* | 34 | RKI |
| 21-02635 | Stool | AMP | *bla*<sub>TEM-1</sub> | 2212 | RKI |
| 21-03310 | Stool | AMP, CAZ, CTX | *bla*<sub>CTX-M-1</sub> | 34 | RKI |
| 21-03504 | Stool | AMP, CIP, NAL, SXT, TCY, TMP | *bla*<sub>TEM-1</sub>, *qnrB19*, *sul2*, *tet(A)*, *dfrA5* | 19 | RKI |
| 21-03926 | Stool | AMP, TCY | *bla*<sub>TEM-1</sub>, *sul2*, *tet(B)* | 34 | RKI |
| 21-03980 | Stool | Sensitive | None | 19 | RKI |
| 21-06117 | Stool | CIP, NAL | *qnrB19*, *sul1* | 19 | RKI |
| 21-06637 | Stool | Sensitive | None | 213 | RKI |
| 22-00115 | Stool | AMP, TCY | *bla*<sub>TEM-1</sub>, *sul2*, *tet(B)* | 34 | RKI |
| 22-00733 | Stool | Sensitive | None | 128 | RKI |
| 22-00857 | Stool | Sensitive | None | 568 | RKI |
| 22-01508 | Stool | AMP, TCY | *bla*<sub>TEM-1</sub>, *sul2*, *tet(B)* | 34 | RKI |
| 22-02255 | Stool | Sensitive | None | 313 | RKI |
| 22-02256 | Stool | AMP | *bla*<sub>TEM-1</sub> | 19 | RKI |
| 22-02294 | Stool | Sensitive | None | 376 | RKI |
| 22-03517 | Stool | AMP, CAZ, CIP, CTX, FOX, TCY | *bla*<sub>CMY-2</sub>, *qnrS1*, *tet(A)* | 36 | RKI |
| 22-03880 | Stool | AMP | *bla*<sub>TEM-1</sub> | 19 | RKI |
| 22-03956 |  | AMP, CTX | *bla*<sub>TEM-1</sub>, *bla*<sub>CTX-M-1</sub> | 19 | RKI |
| 22-04189 | Stool | AMP, CAZ, CIP, CTX | *bla*<sub>TEM-1</sub>, *bla*<sub>CTX-M-1</sub>, *qnrS1* | 19 | RKI |
| 22-04406 | Stool | Sensitive | None | 19 | RKI |
| 22-05027 | Stool | Sensitive | None | 36 | RKI |
| 22-05885 | Urine | AMP, CHL, TCY | *bla*<sub>CARB-2</sub>, *floR*, *tet(G)* | 19 | RKI |
| 22-06301 | Stool | SXT, TMP | *sul1* / *sul2*, *dfrA1* | 34 | RKI |
| 22-06509 | Stool | Sensitive | None | 19 | RKI |
| 22-07856 |  | AMP, CIP, CTX, SXT, TCY, TMP | *bla*<sub>CTX-M-1</sub>, *qnrS1*, *sul1*, *tet(A)*, *dfrA12* | 34 | RKI |
| 22-07859 | Blood | AMP | *bla*<sub>TEM-1</sub> | 19 | RKI |
| 22-07954 | Stool | AMP, CIP, NAL, TCY | *bla*<sub>TEM-1</sub>, *qnrB19*, *sul2*, *tet(B)* | 34 | RKI |
| 23-00671 |  | AMP, TCY | *bla*<sub>TEM-1</sub>, *sul2*, *tet(B)* | 34 | RKI |
| 23-00837 | Blood | Sensitive | None | 19 | RKI |
| 23-00954 | Stool | Sensitive | None | 19 | RKI |
| 23-01121 | Blood | Sensitive | None | 313 | RKI |
| 23-01218 | Blood | AMP, SXT, TMP | *bla*<sub>TEM-1</sub>, *sul1* / *sul2*, *dfrA1* | 313 | RKI |
| 23-04527 | Blood | AMP | *bla*<sub>TEM-1</sub> | 19 | RKI |
| **Other *S. enterica* serovars** |  |  |  |  |  |
| 18-00670,<br>Serovar<br>Choleraesuis | blood | STR | None | 145 | RKI |
| 18-04107,<br>Serovar<br>Enteritidis |  | Sensitive | None | 11 | RKI |
| 18-04810,<br>Serovar<br>Agona | stool | Sensitive | None | 13 | RKI |
| 19-01006,<br>Serovar<br>Choleraesuis |  | Sensitive | None | 145 | RKI |
| 19-01481,<br>Serovar<br>Enteritidis | stool | Sensitive | None | 11 | RKI |
| 19-02948,<br>Serovar<br>Kentucky | stool | AMP, CIP, GEN,<br>MEZ, MSU,<br>NAL, OTE, STR | *bla*<sub>TEM-1</sub>,<br>*gyrA*_D87Y /<br>*gyrA*_S83F /<br>*parC*_S80I, *sul1*,<br>*tet(A)* | 198 | RKI |
| 19-03178,<br>Serovar<br>Agona | stool | Sensitive | None | 13 | RKI |
| 20-05235,<br>Serovar<br>Derby | stool | Sensitive | None | 39 | RKI |
| 20-06129,<br>Serovar<br>Enteritidis | stool | Sensitive | None | 11 | RKI |
| 21-01720,<br>Serovar<br>Infantis | stool | AMP, CIP,<br>CMP, NAL,<br>SXT, TCY, TMP | *bla*<sub>TEM-1</sub>,<br>*gyrA*_S83Y, *cmlA1*,<br>*sul1* / *sul3*, *tet(A)*,<br>*dfrA8* | 2283 | RKI |
| 21-02525,<br>Serovar<br>Derby | stool | Sensitive | None | 682 | RKI |
| 21-02979,<br>Serovar<br>Enteritidis | stool | Sensitive | None | 183 | RKI |
| 21-06414,<br>Serovar<br>Derby | stool | Sensitive | None | 40 | RKI |

**Table 7.** Mass spectrometric parameters used for the quantification of C26 and SW-C182.

|  | Q1<br>(mass)<br>[g/mol] | Q3<br>(mass)<br>[g/mol] | Declustering<br>potential [V] | Collision<br>energy<br>[V] | Collision<br>cell exit<br>potential<br>[V] |
| --- | --- | --- | --- | --- | --- |
| **Caffeine (IS)** |  |  |  |  |  |
| quantifier | 195.116 | 138.1 | 81 | 27 | 10 |
| qualifier | 195.116 | 110.1 | 81 | 31 | 6 |
| **C26** |  |  |  |  |  |
| quantifier | 397.52 | 174.7 | 11 | 25 | 10 |
| qualifier | 397.52 | 134.9 | 11 | 21 | 10 |
| **SW-C182** |  |  |  |  |  |
| quantifier | 432,96 | 175.0 | 1 | 23 | 12 |
| qualifier | 432,96 | 96.1 | 1 | 81 | 12 |

**Table 8.**
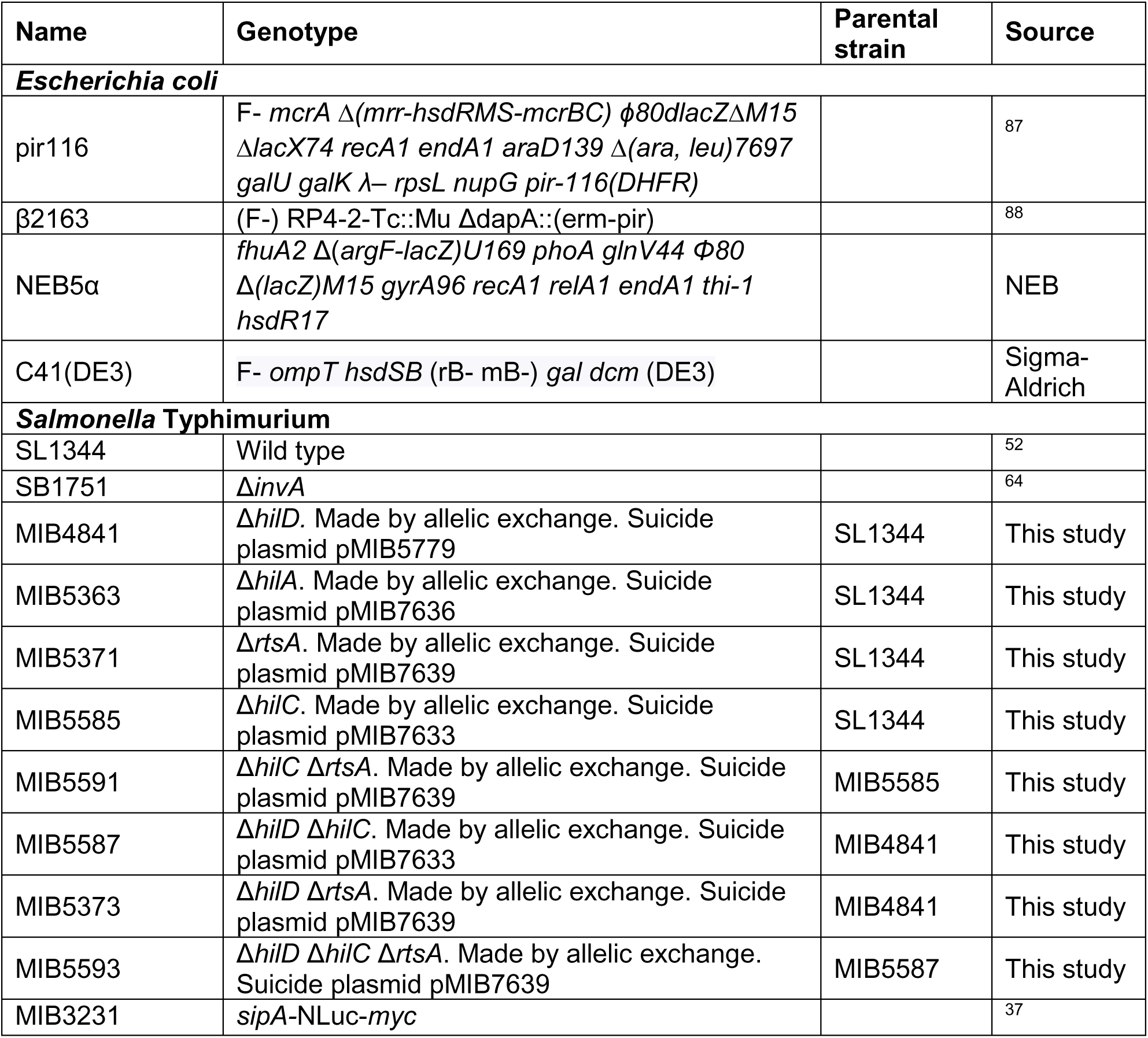
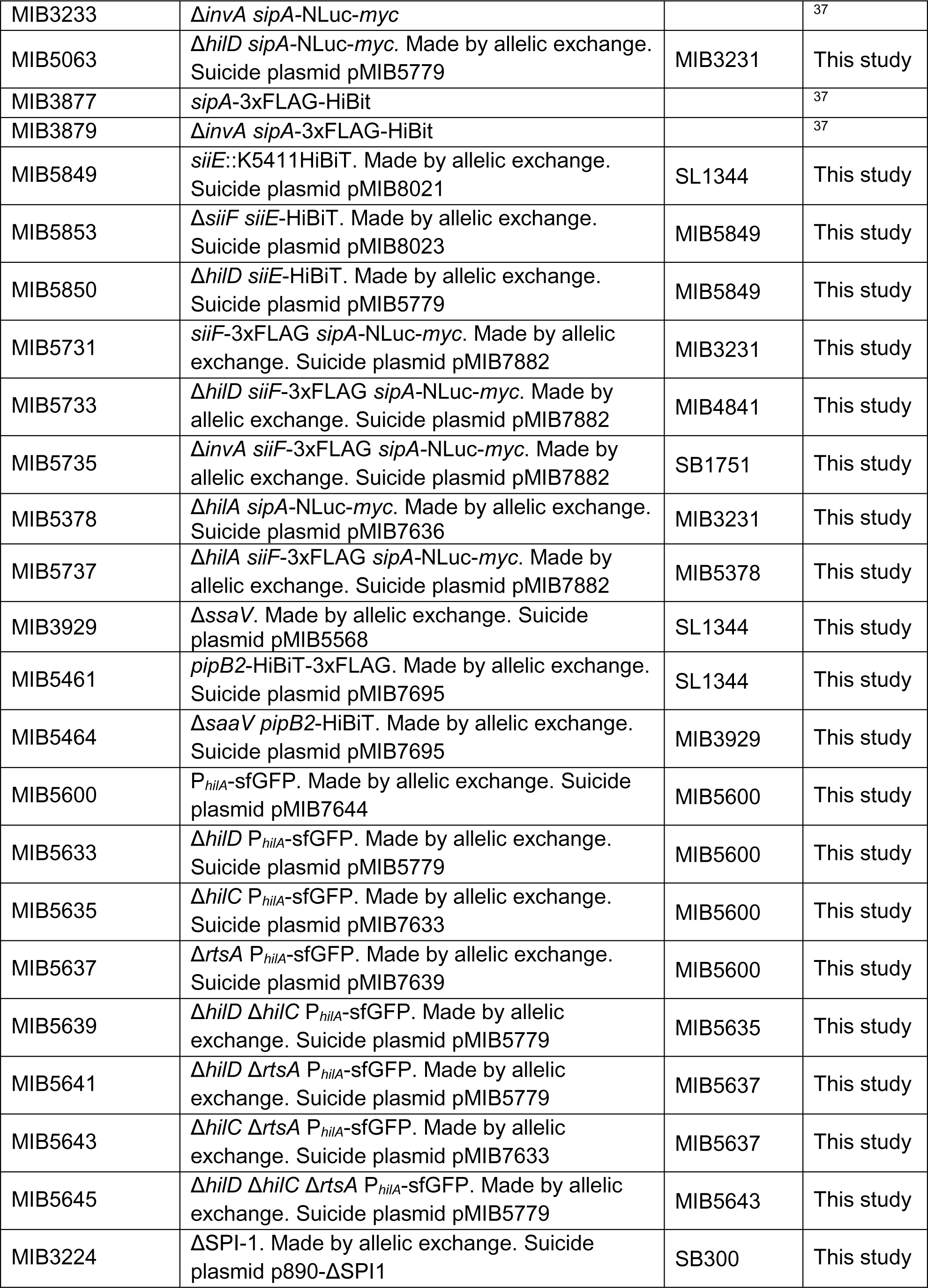
List of strains.

**Table 9.**
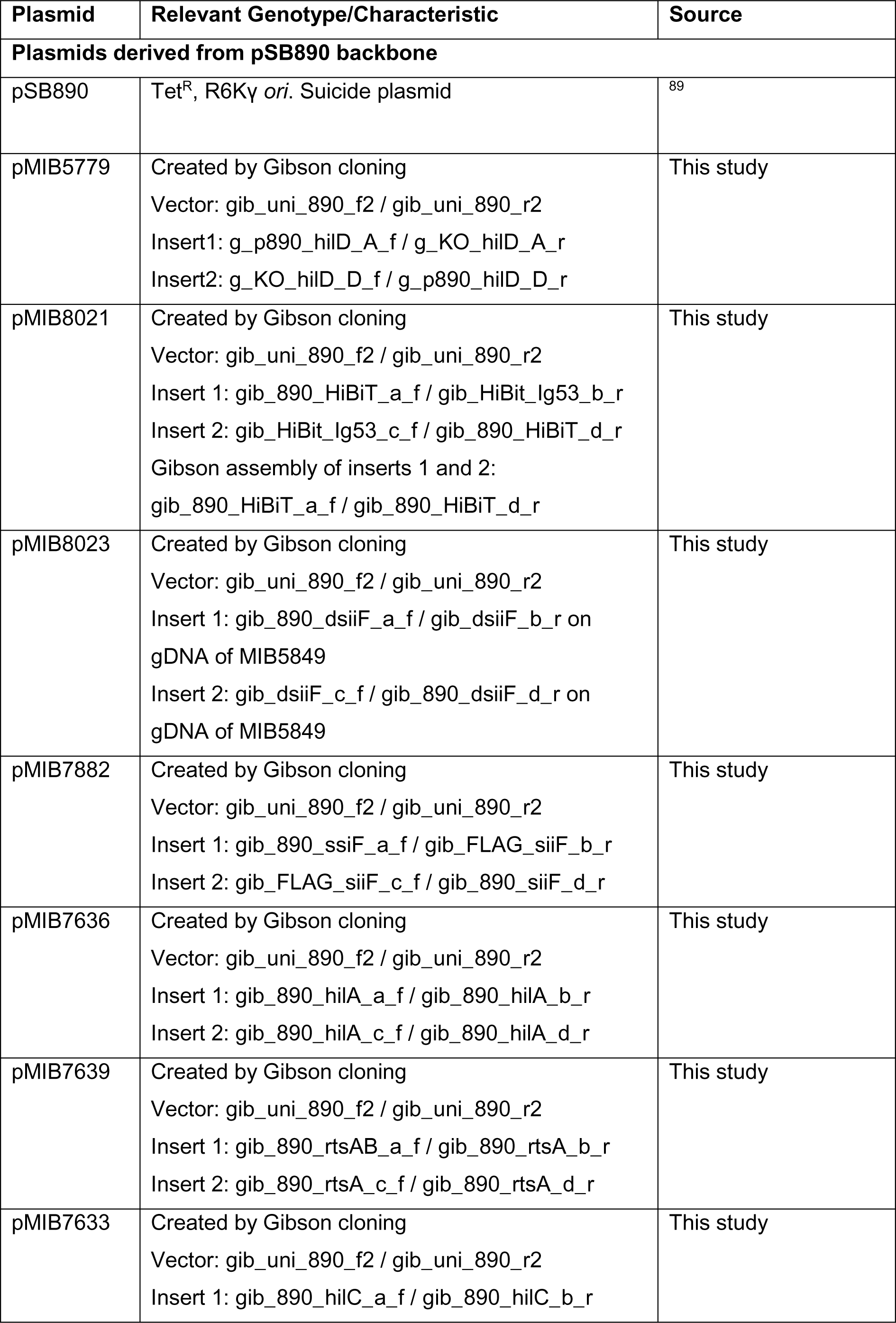
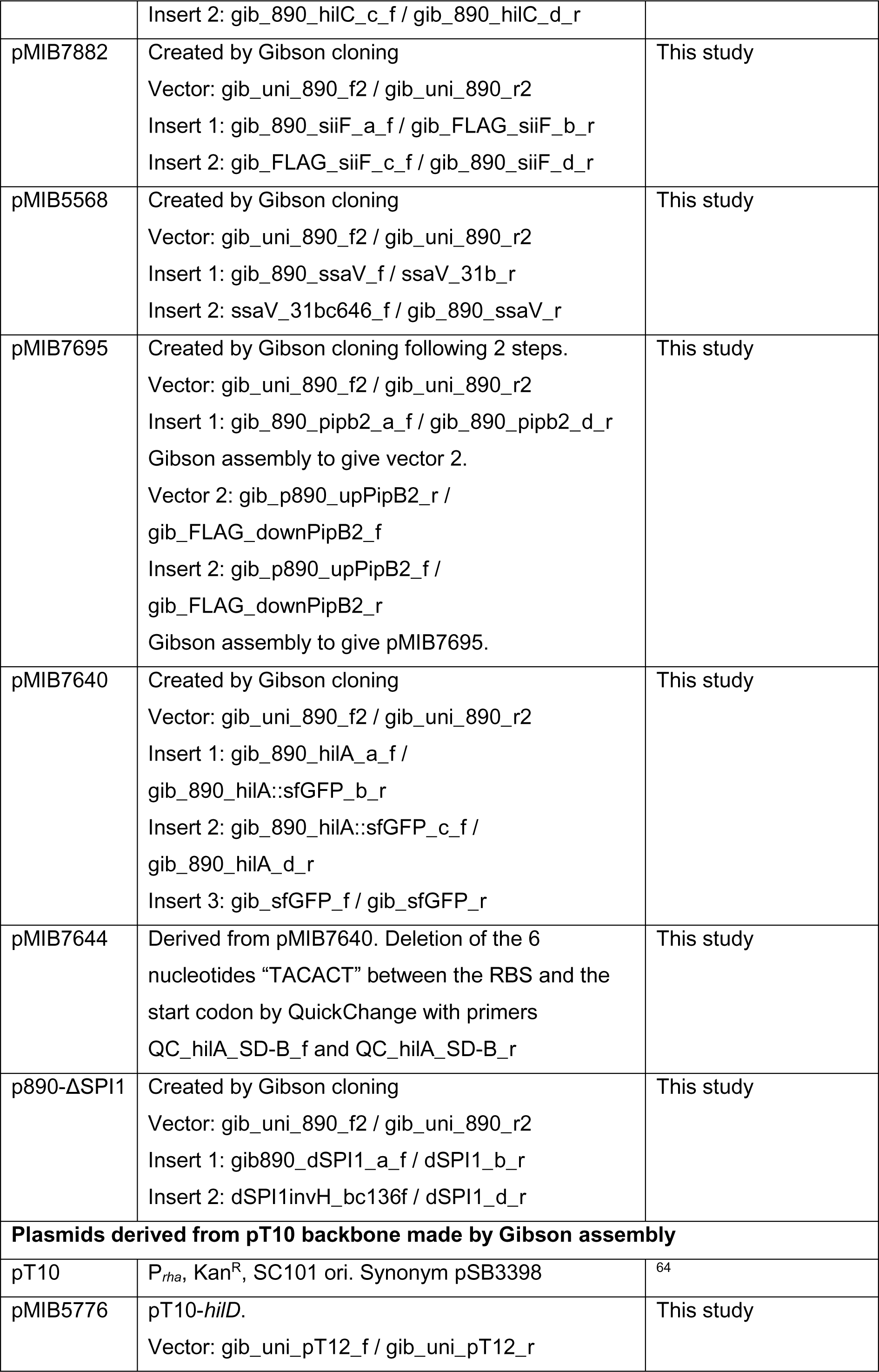
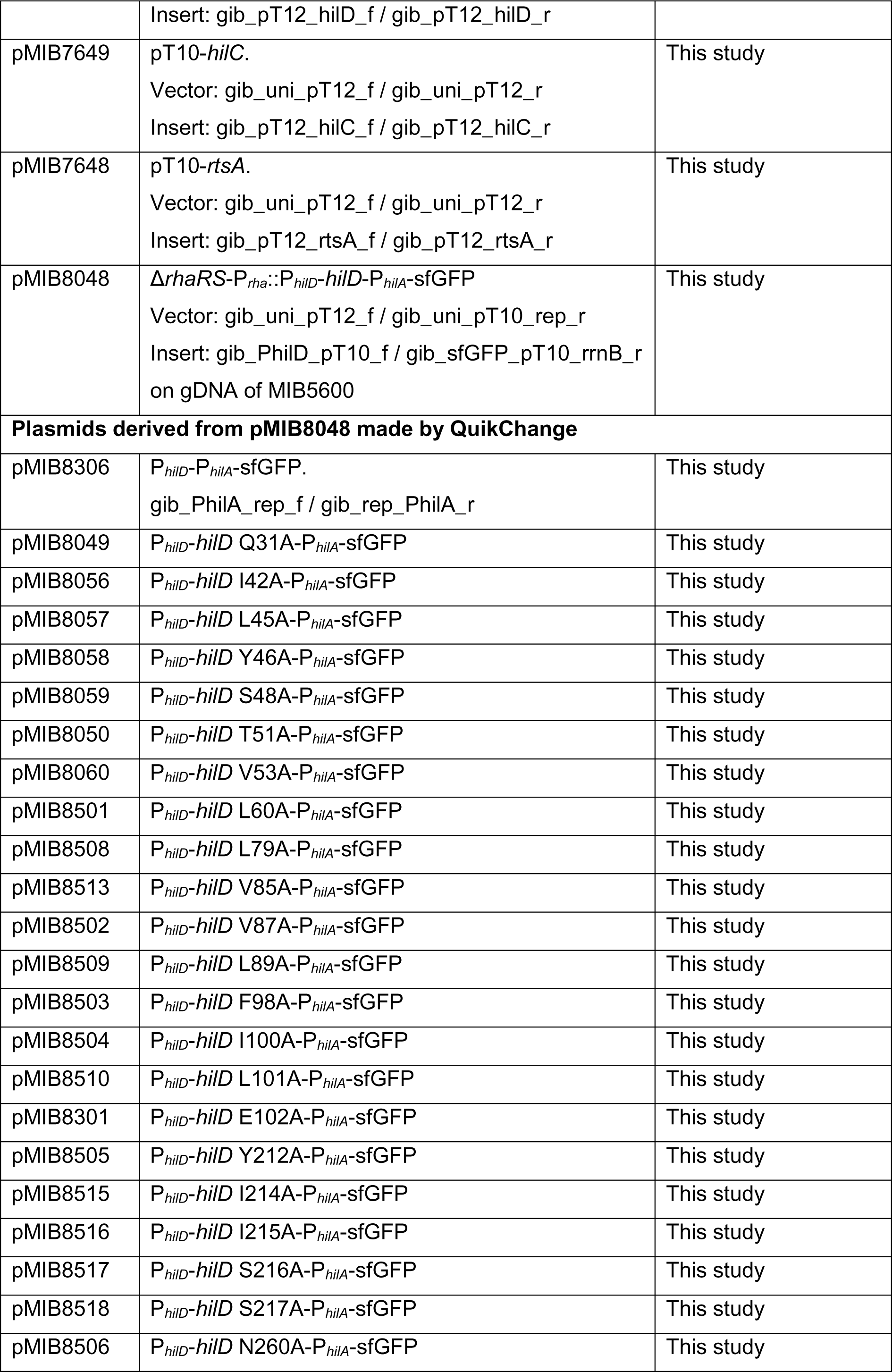
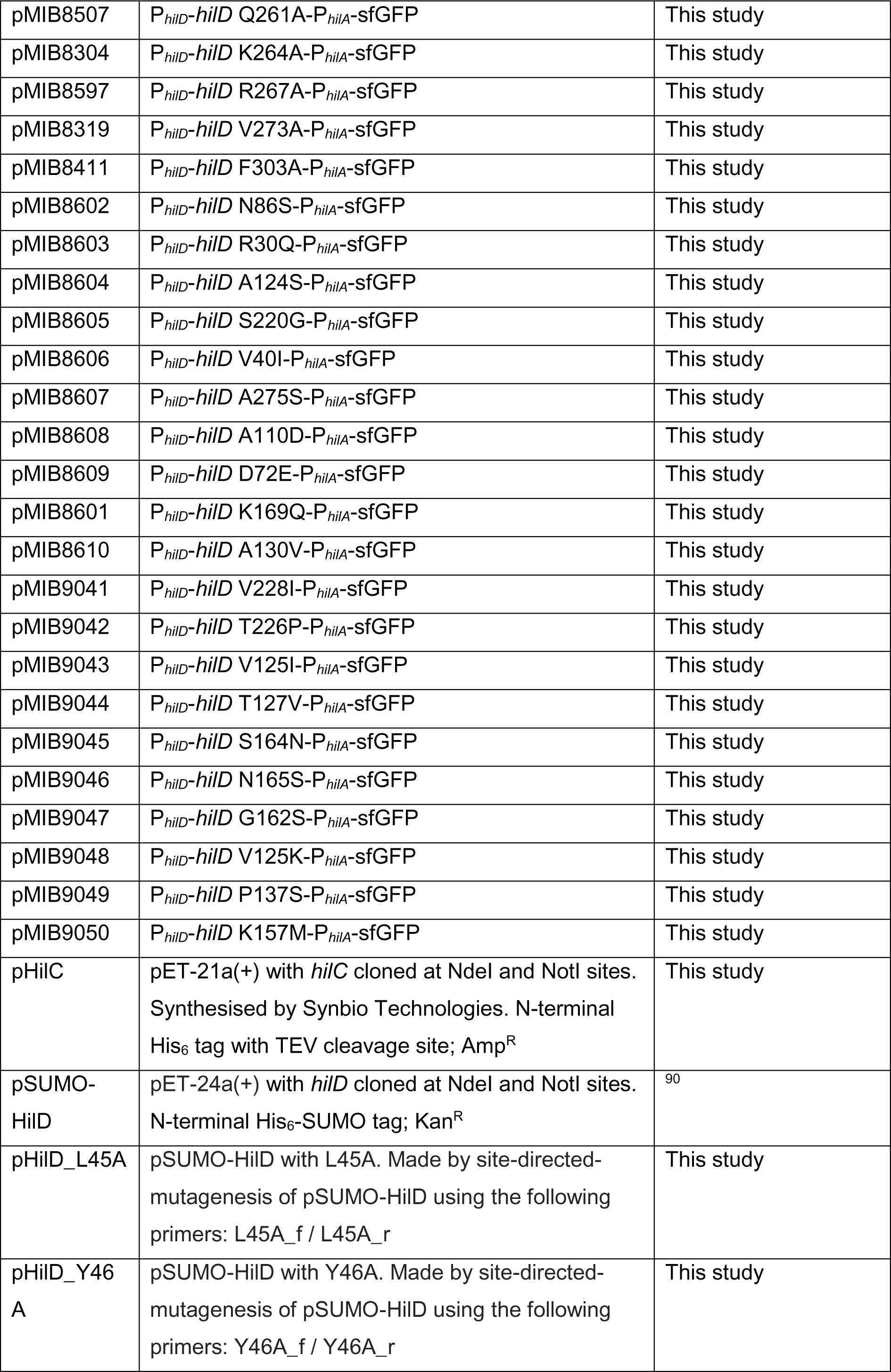
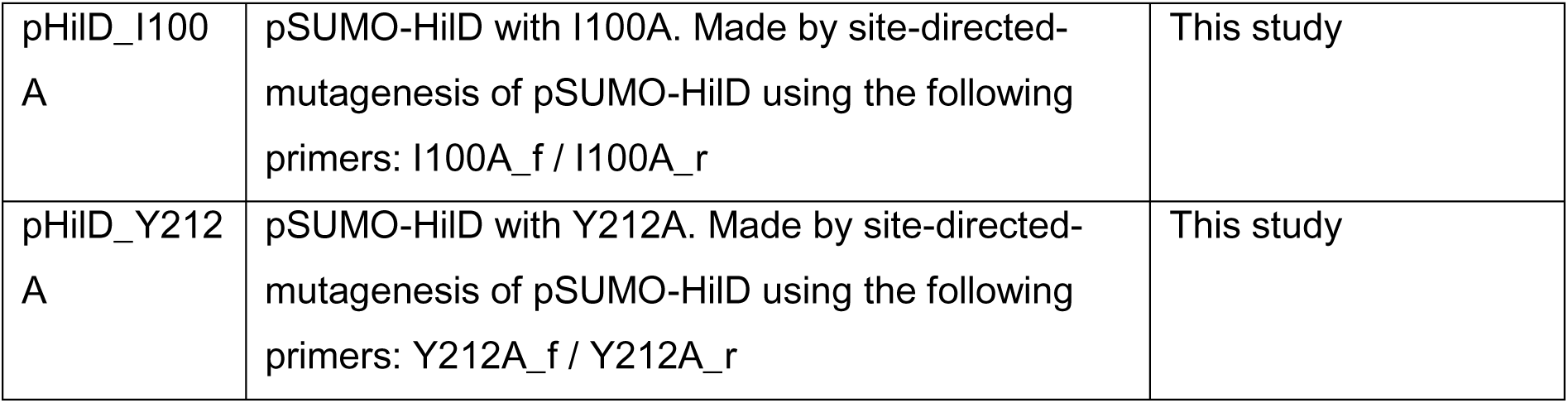
Plasmids. If not otherwise specified, inserts were amplified from genomic DNA of *S.* Typhimurium SL1344.

**Table 10.**
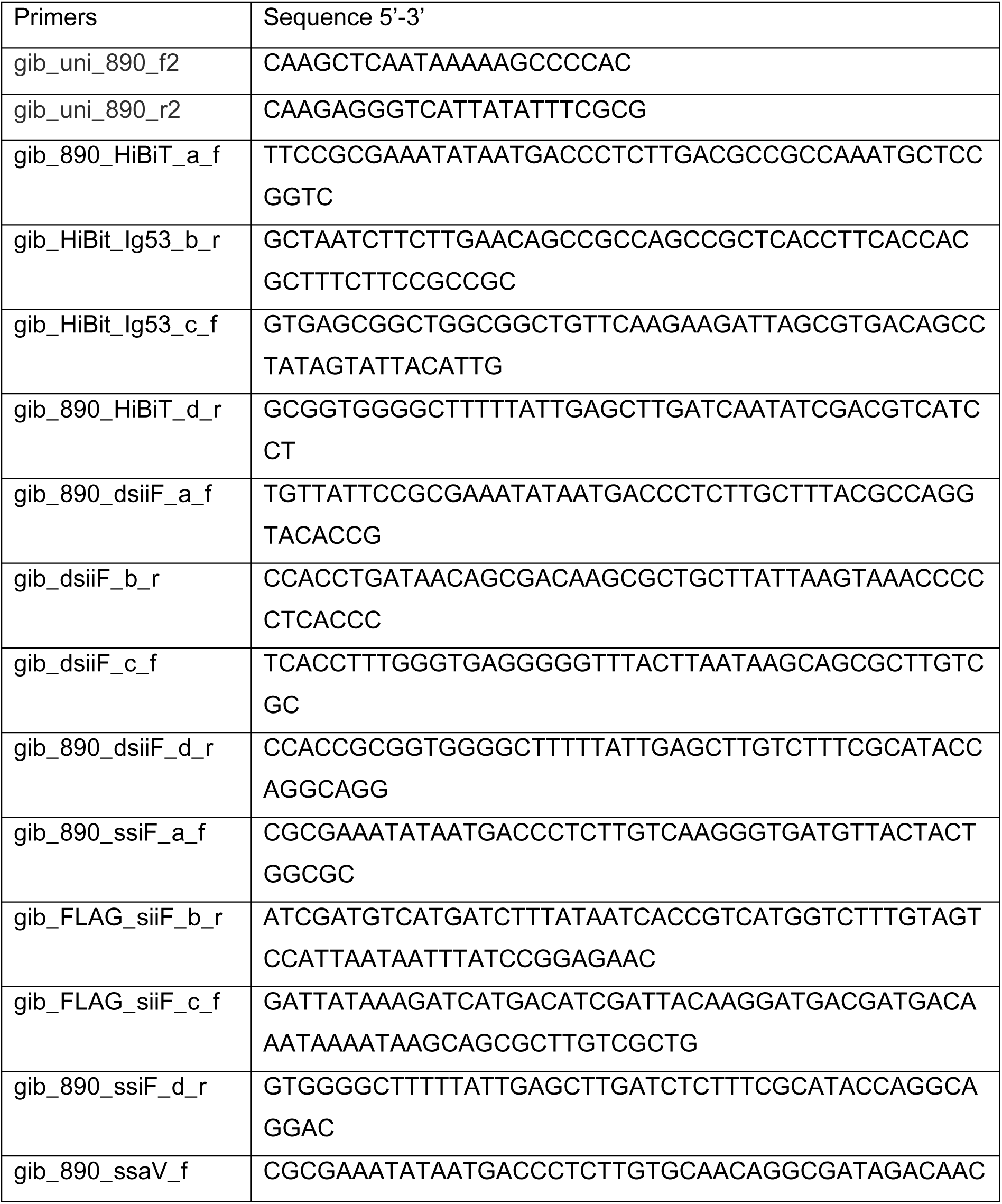
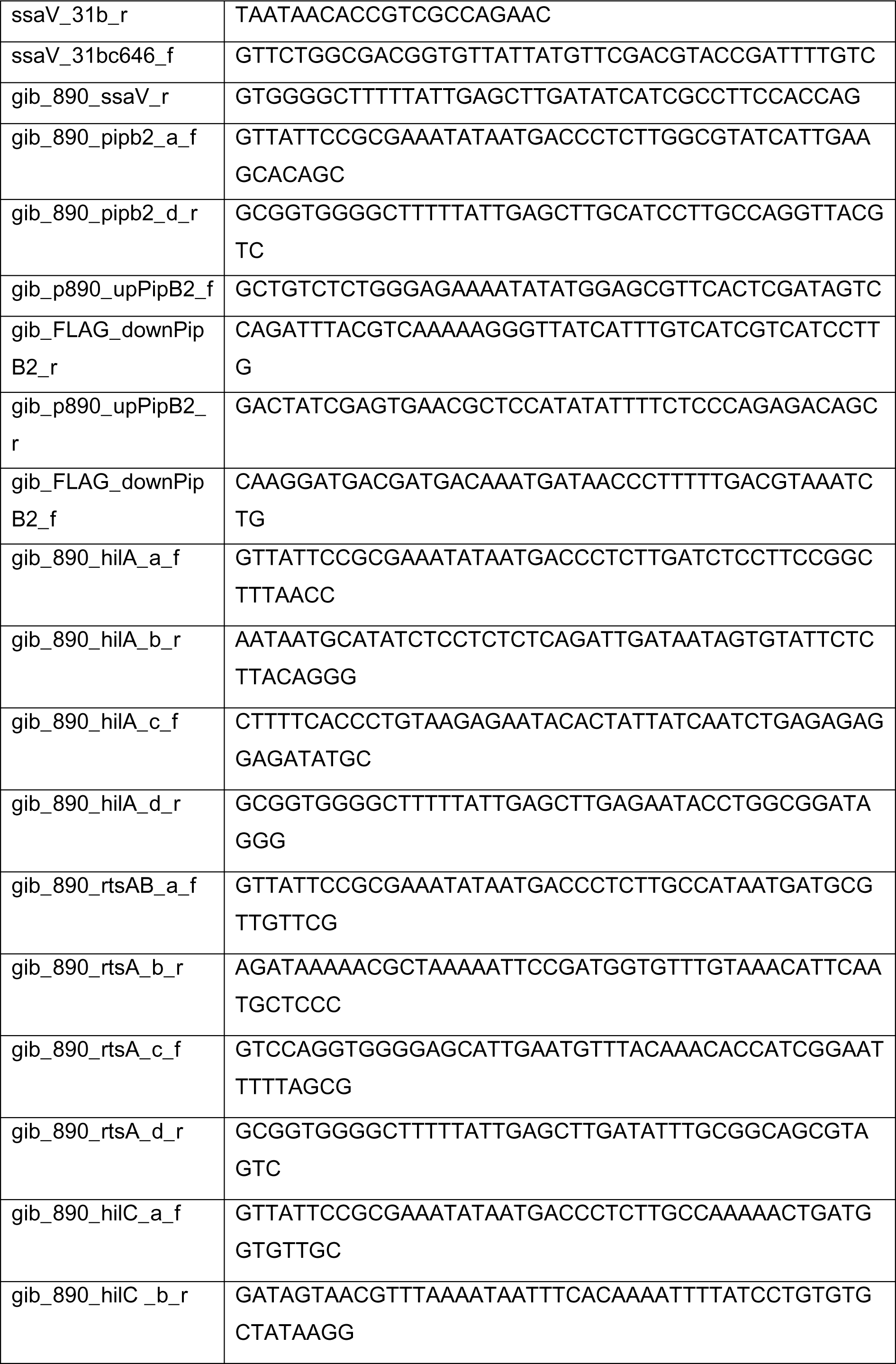
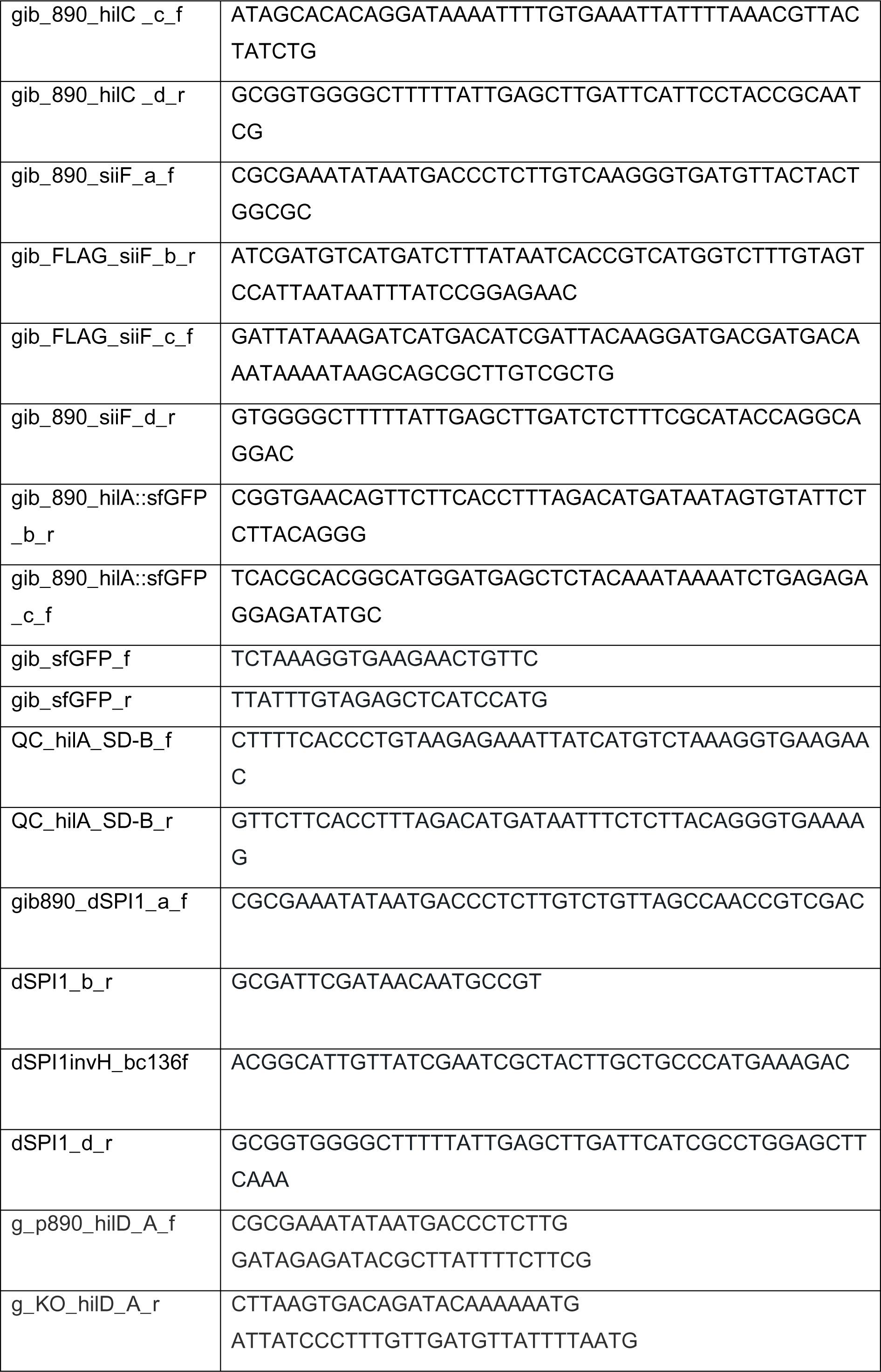
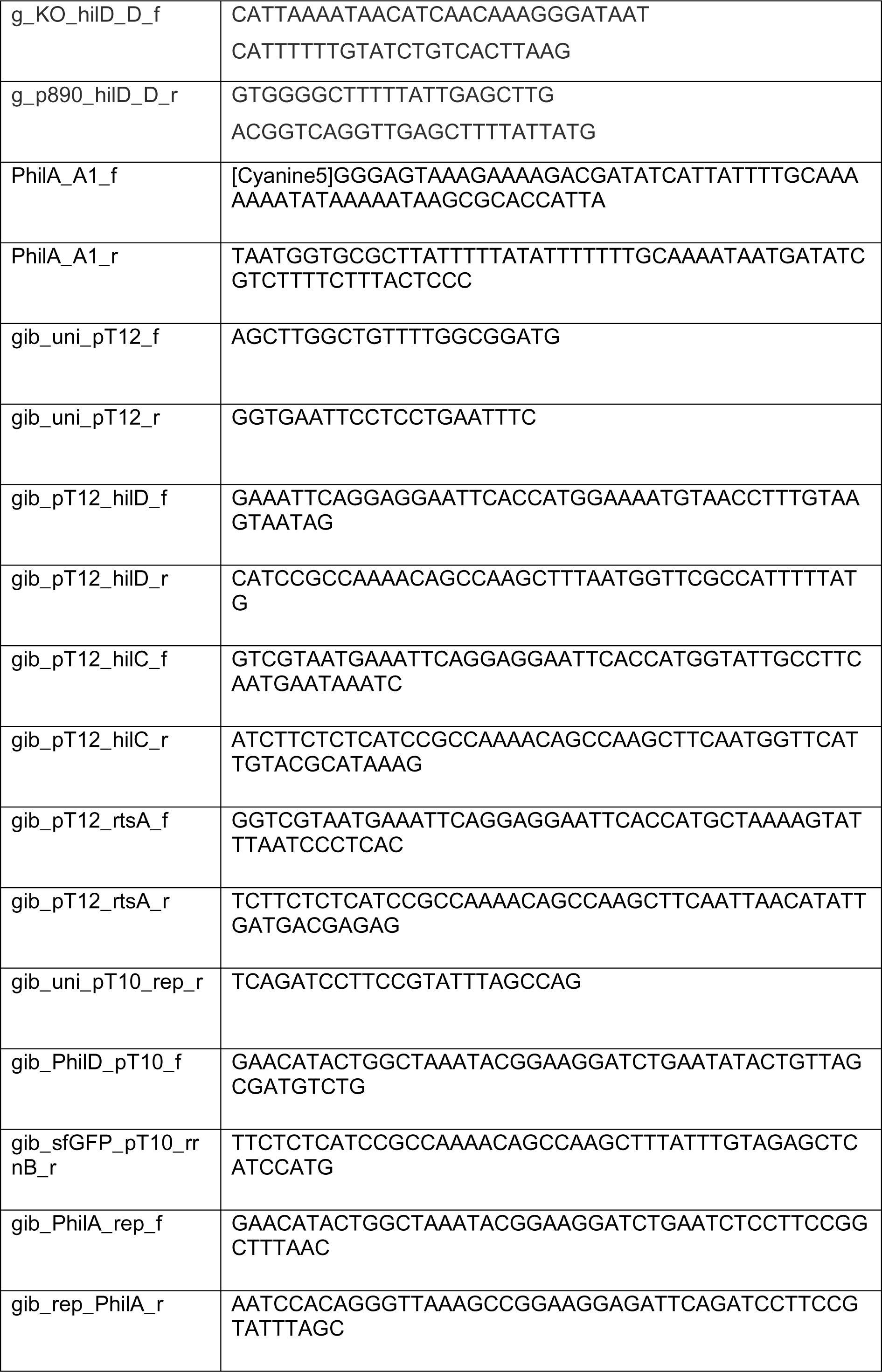
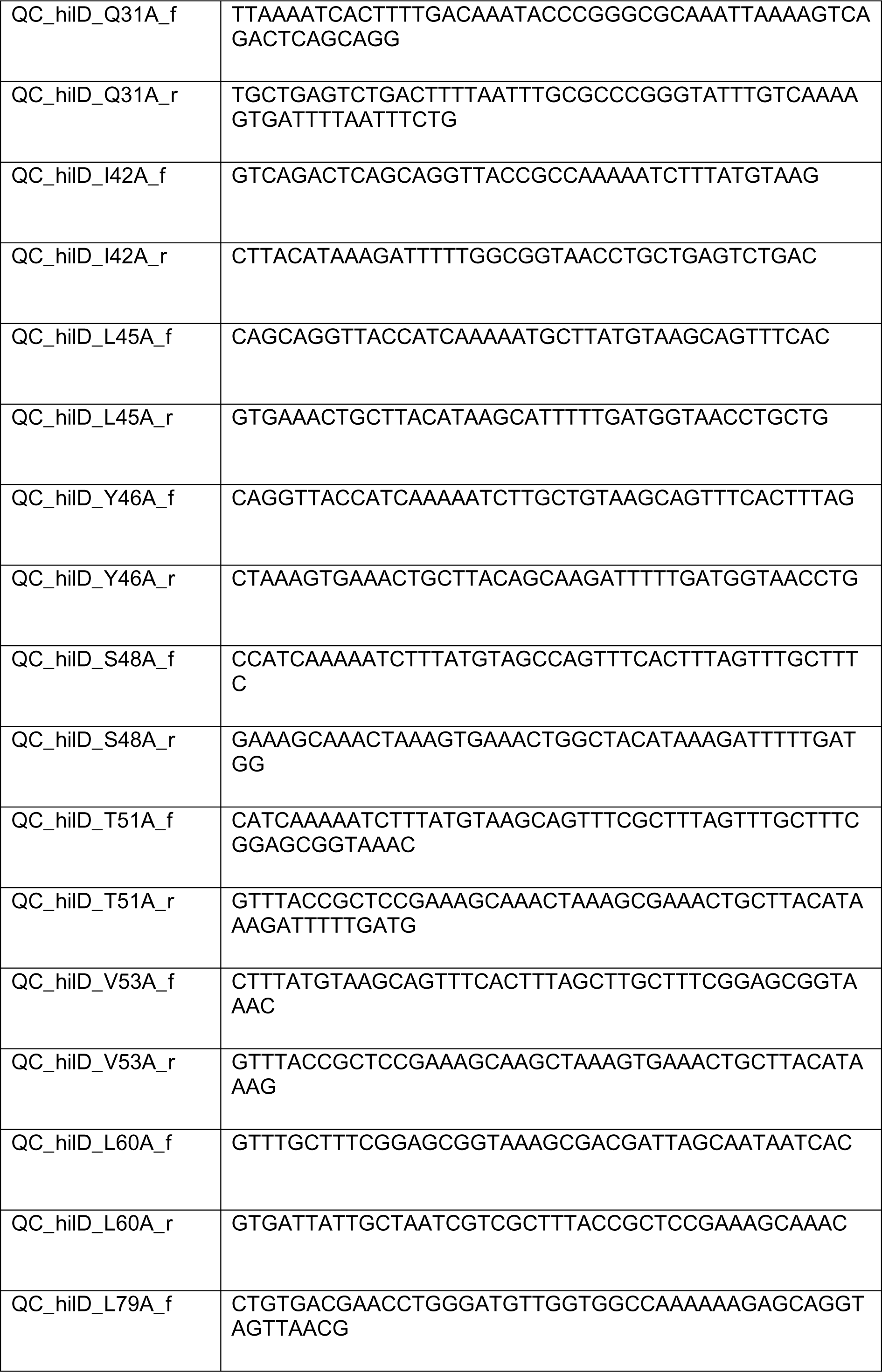
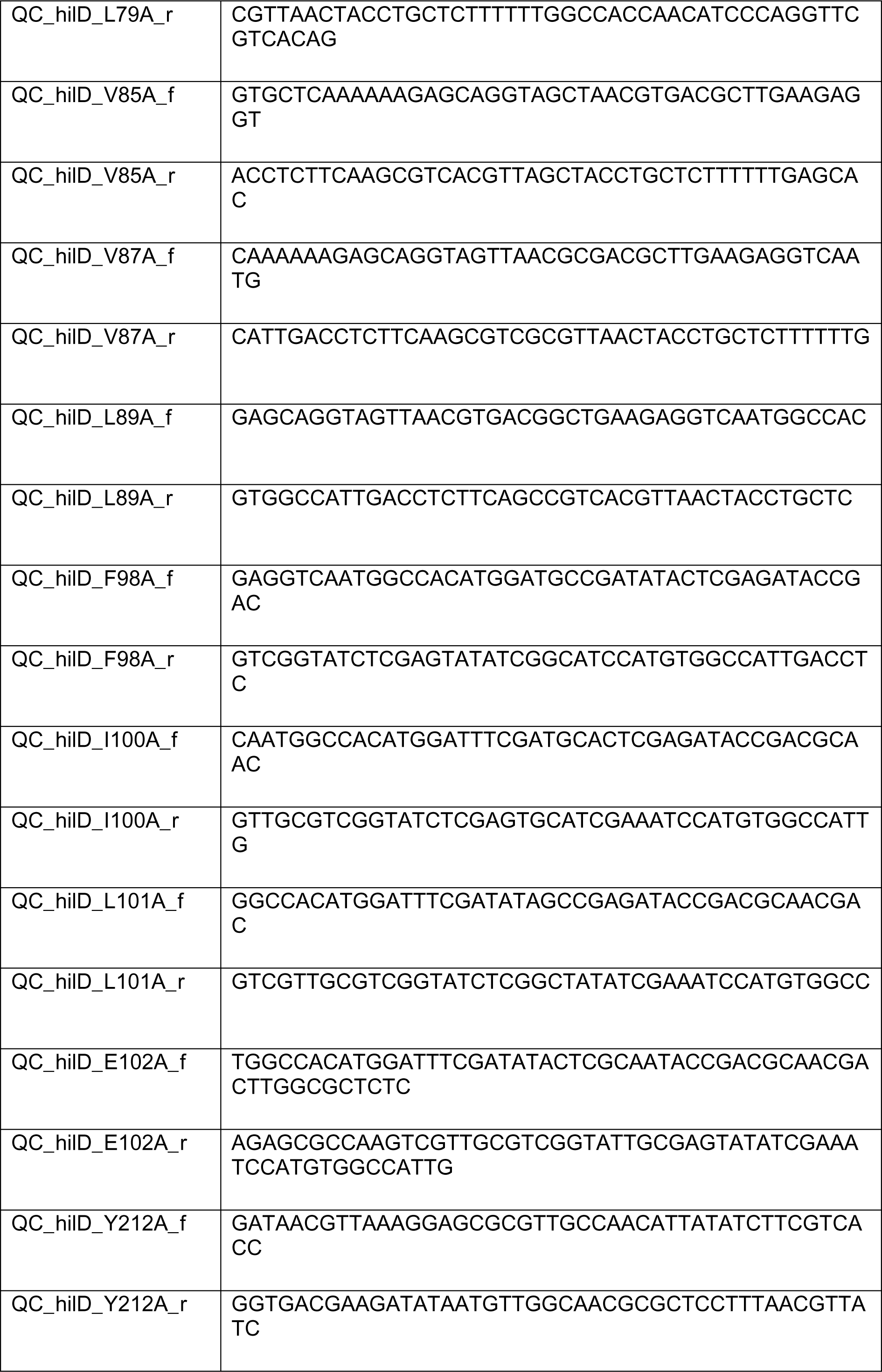
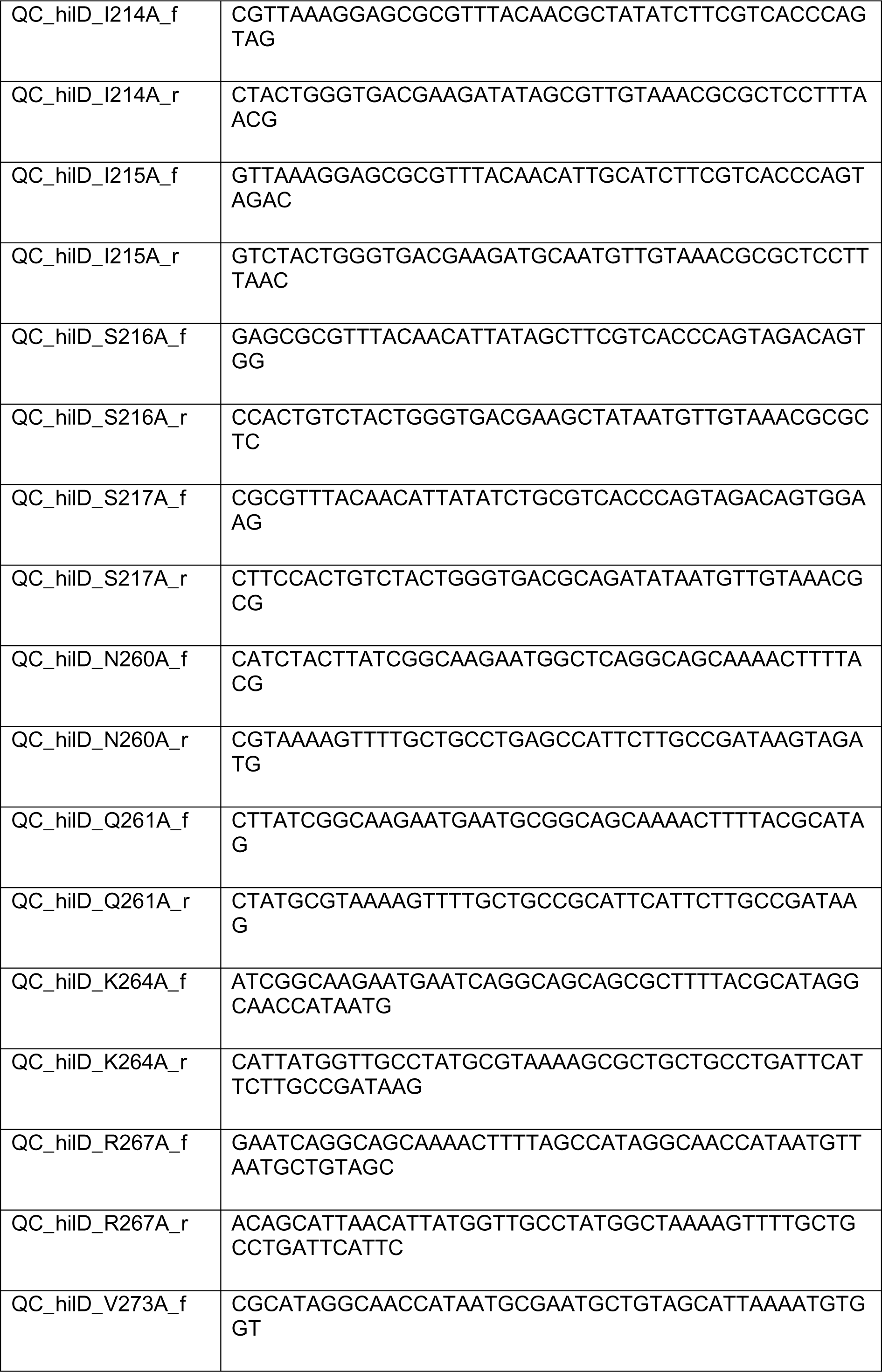
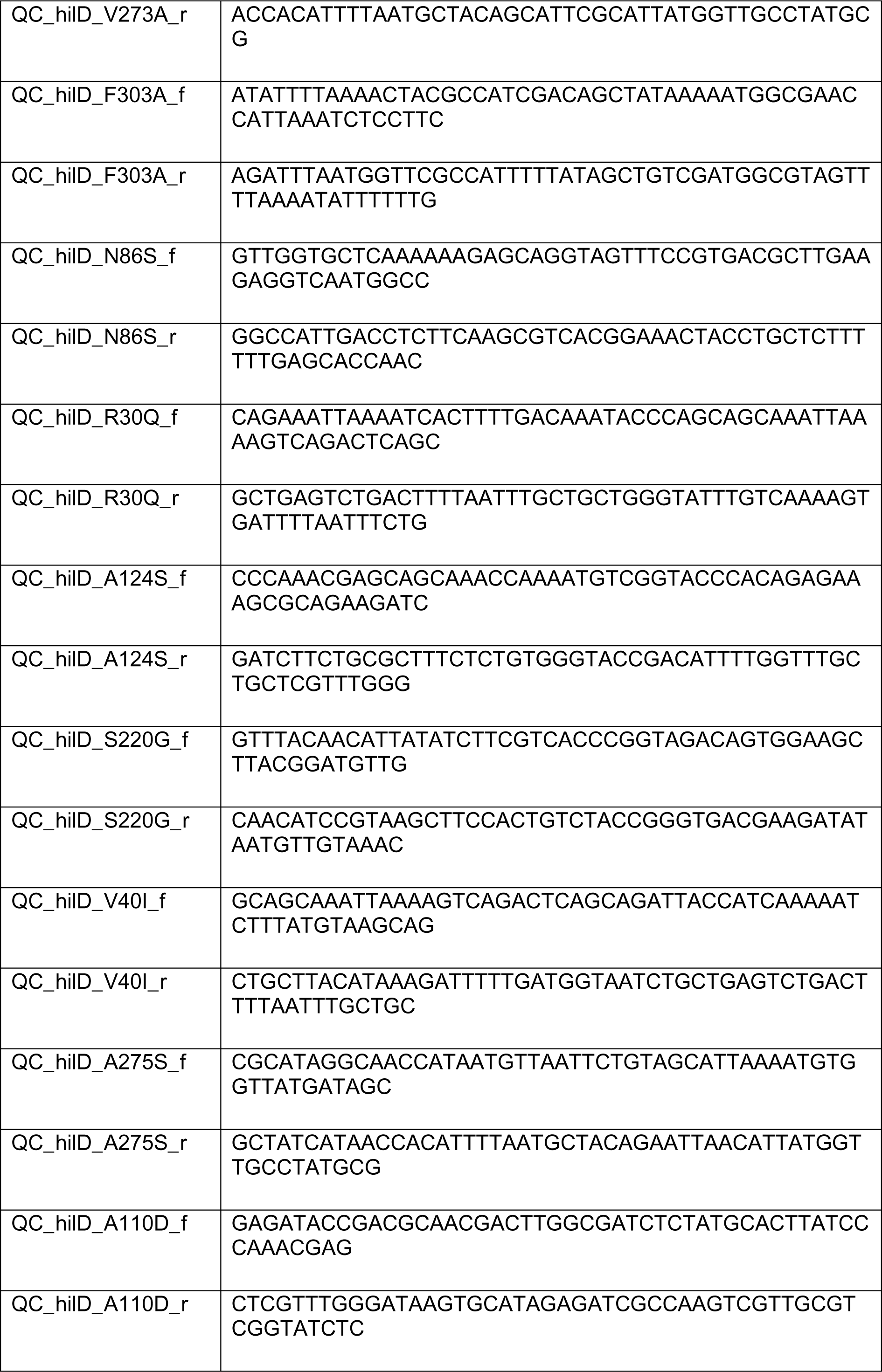
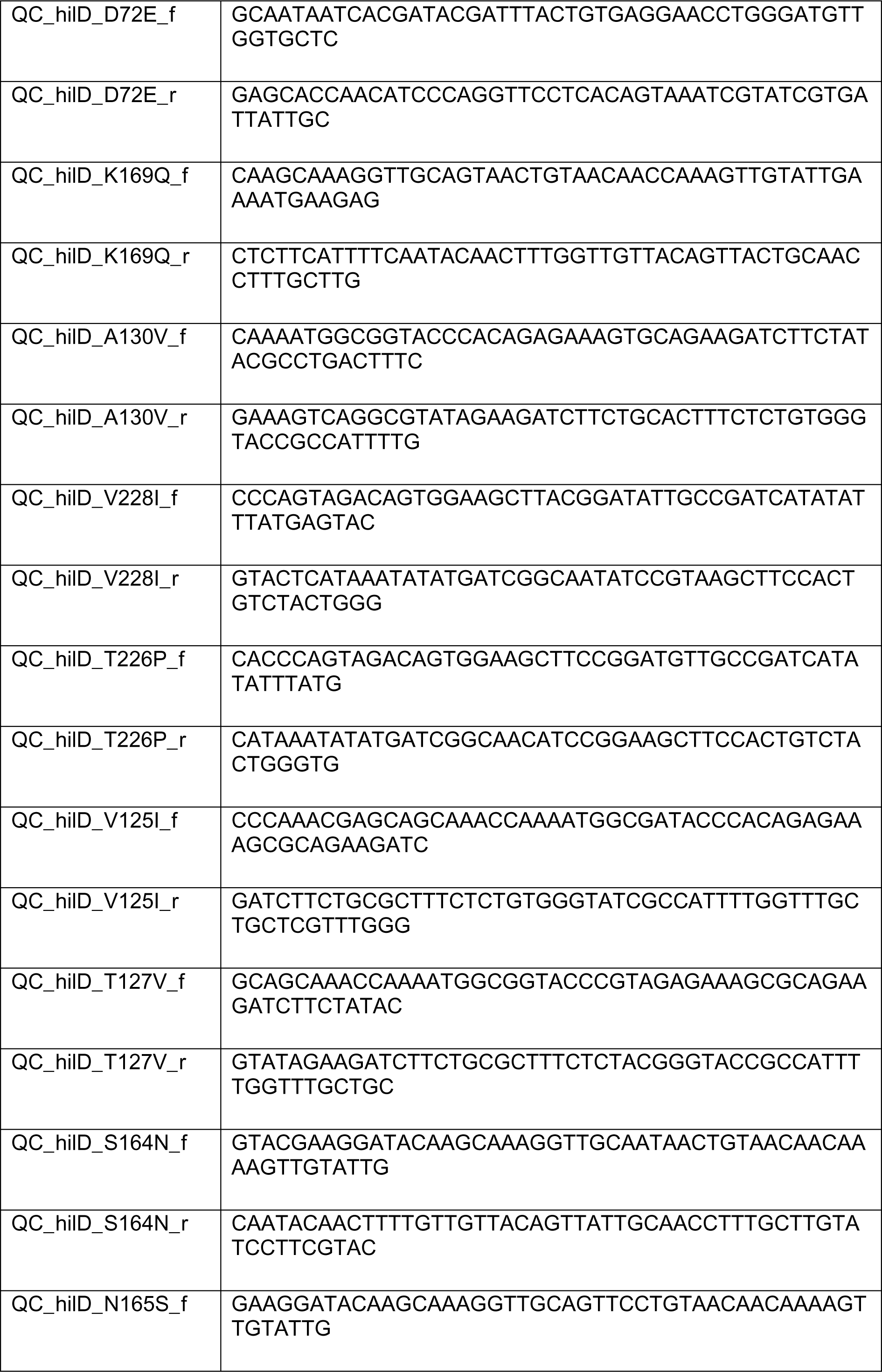
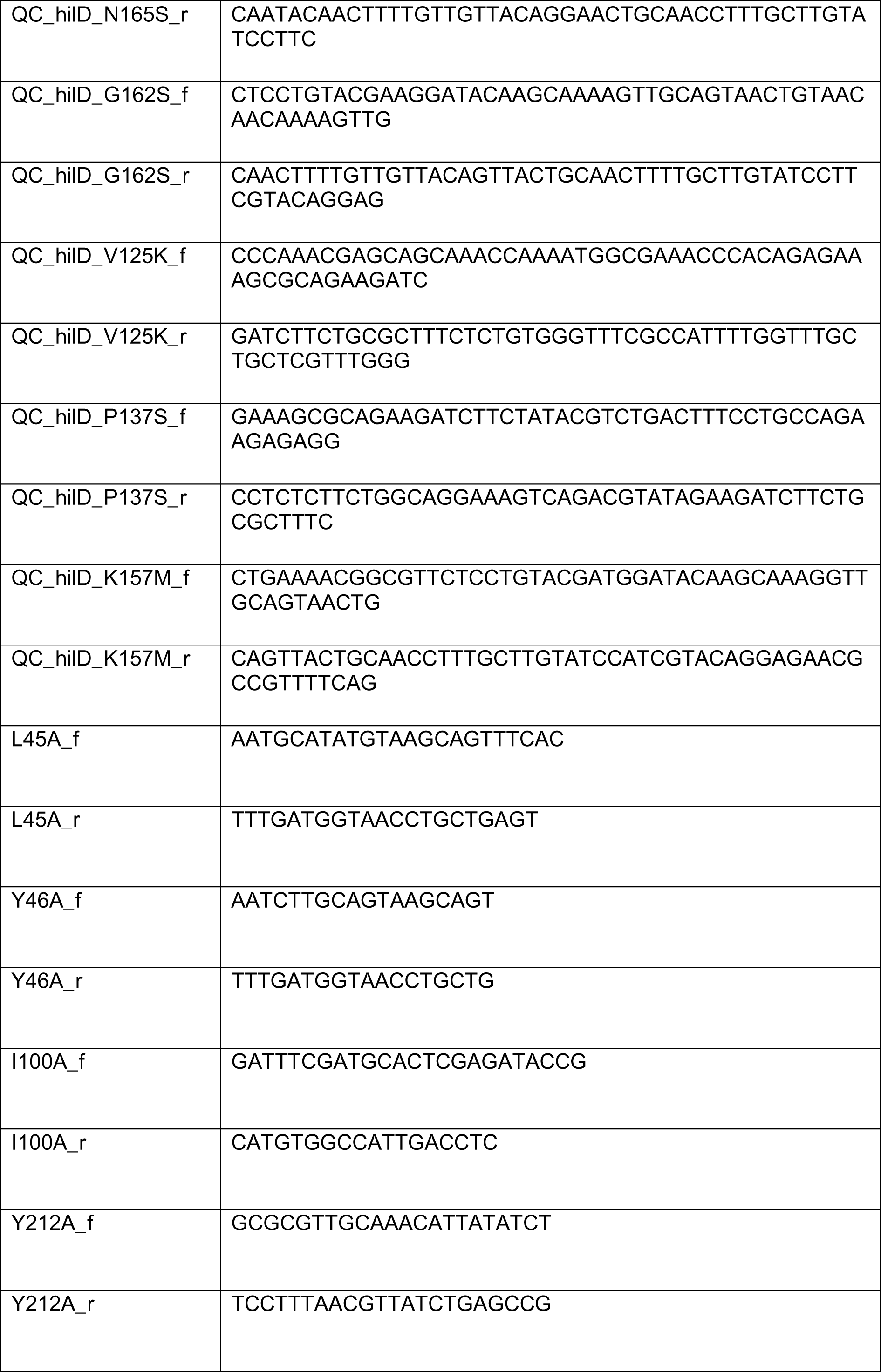
List of primers.

## NMR Spectra

### *N*-(benzo[d][1,3]dioxol-5-ylmethyl)-2-(((5-bromothiophen-2-yl)methyl)(methyl)amino)acetamide (C26)

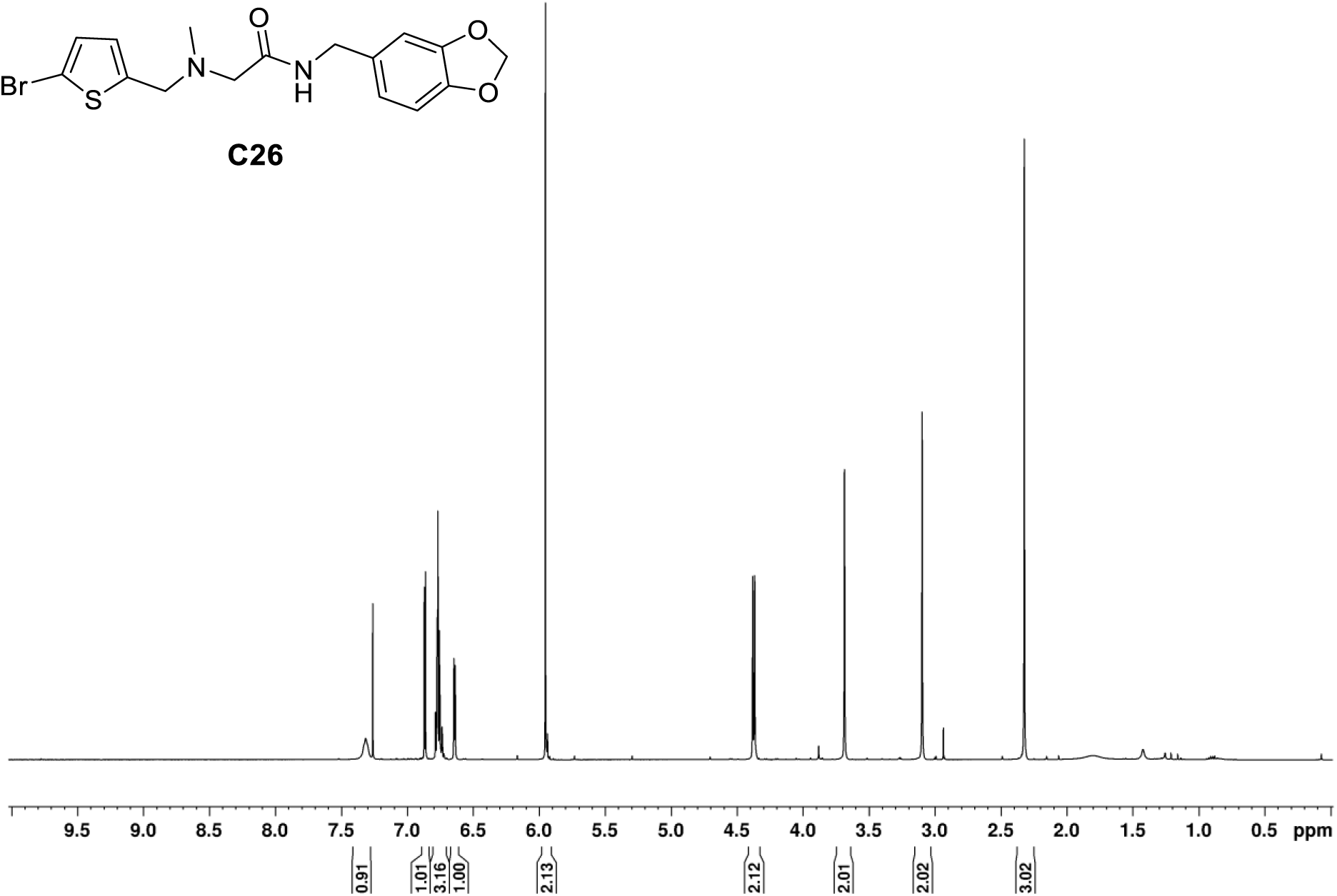

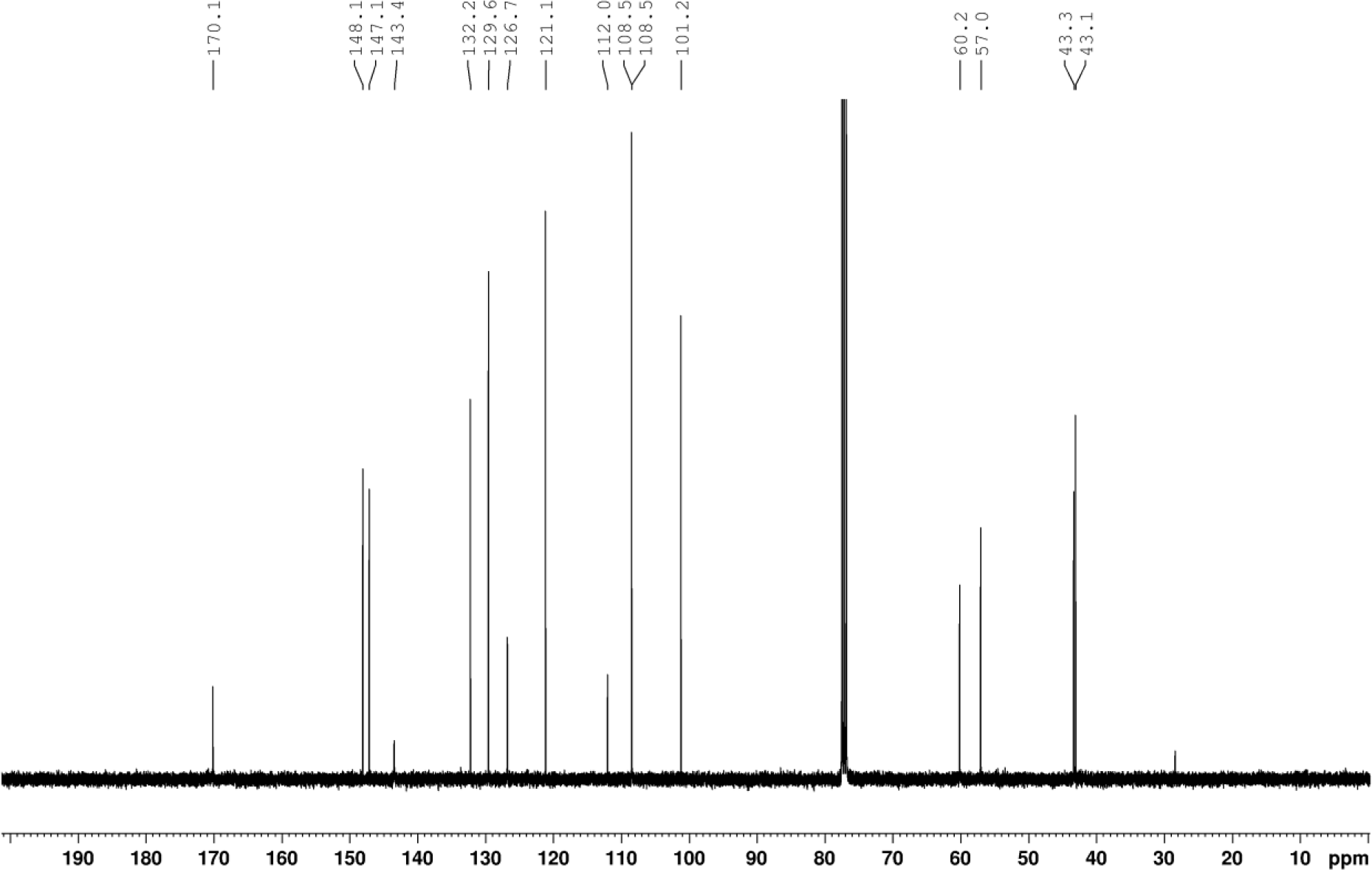

### N-(benzo[d][1,3]dioxol-5-ylmethyl)-2-((2-bromobenzyl)(methyl)amino)acetamide (SW-C165)

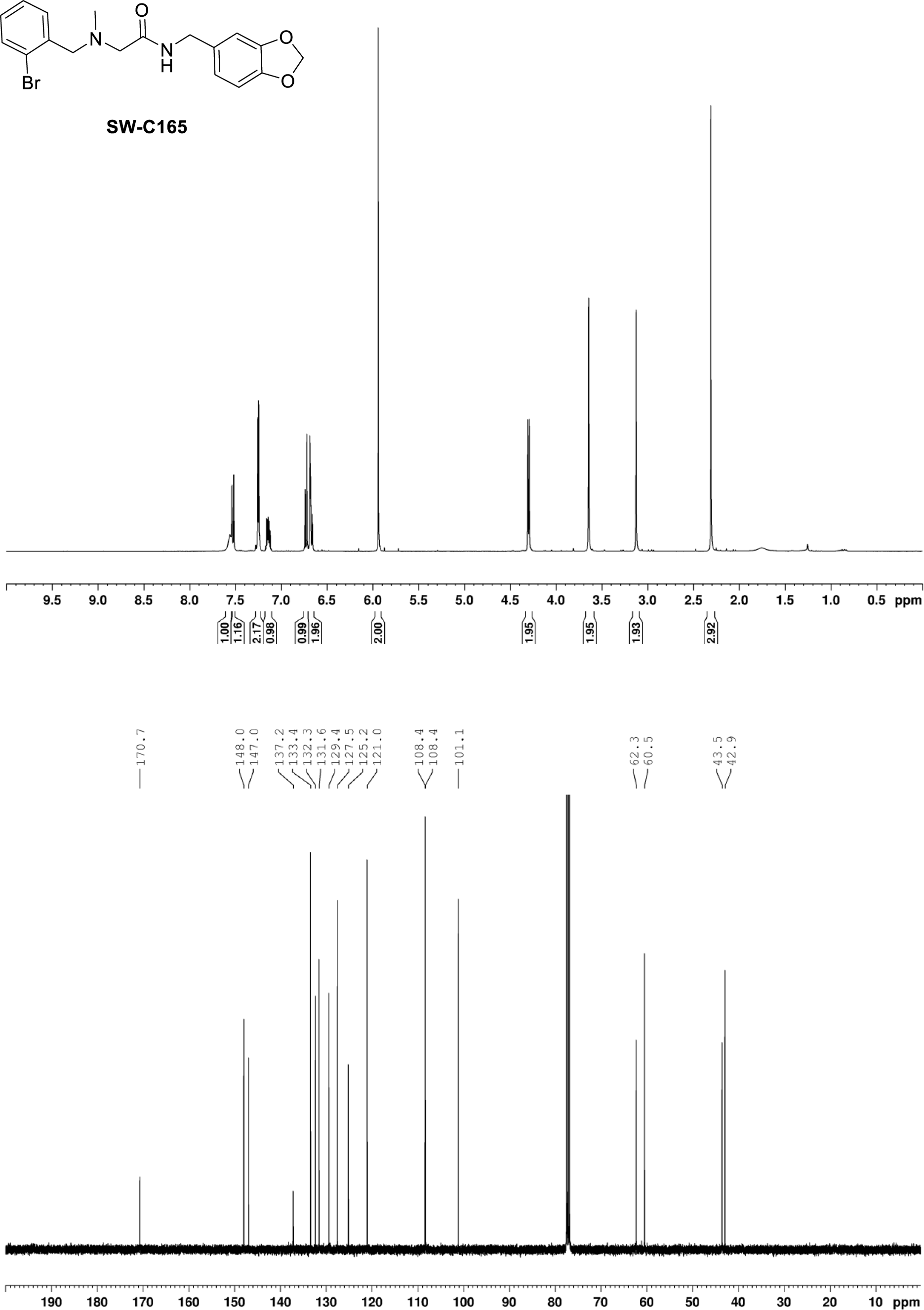

### *N*-(benzo[d][1,3]dioxol-5-ylmethyl)-2-(((5-bromofuran-2-yl)methyl)(methyl)amino)acetamide (SW-C210)

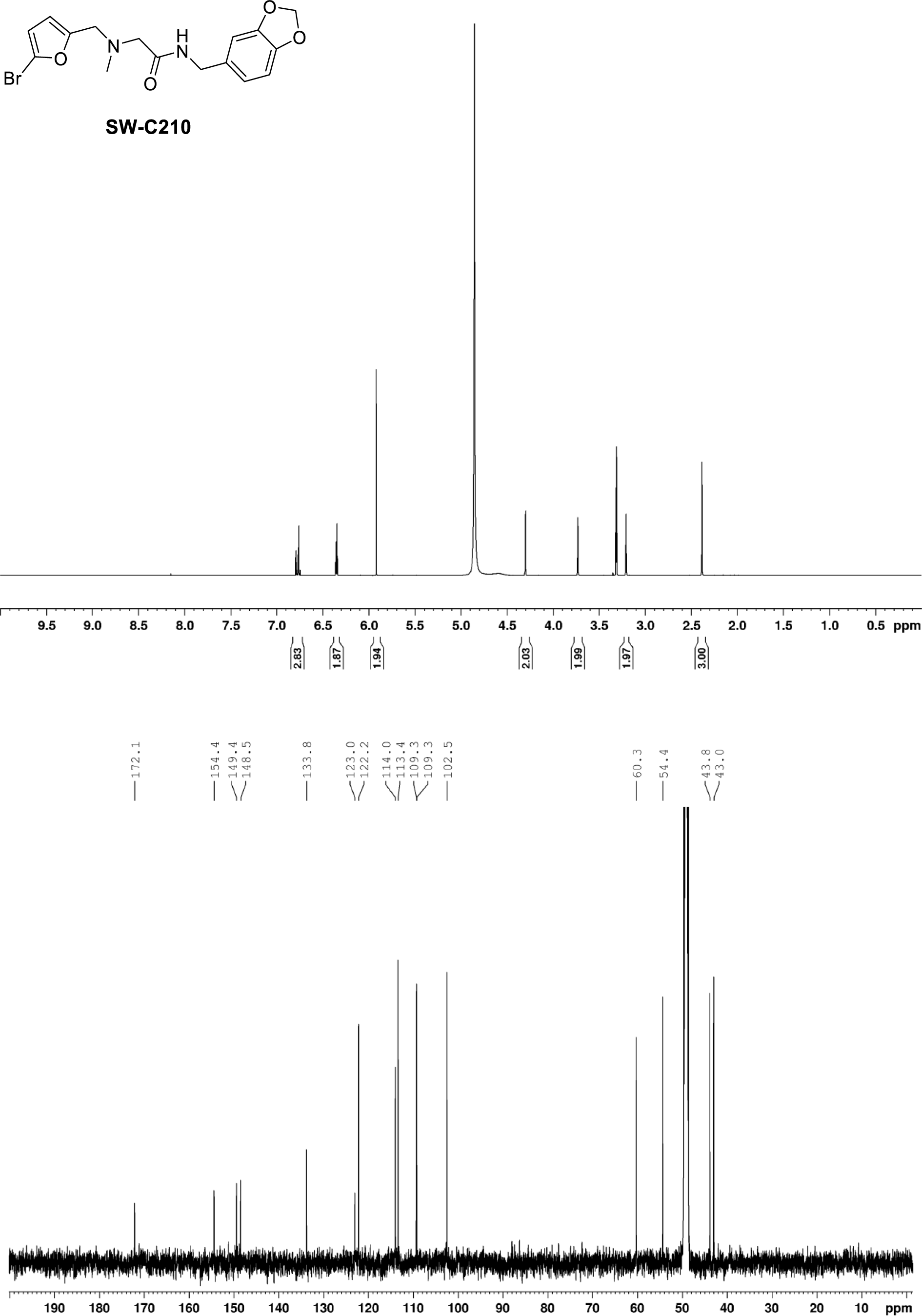

### 2-(((5-Bromothiophen-2-yl)methyl)(methyl)amino)-*N*-(3,5-dichlorobenzyl)acetamide (SW-C250)

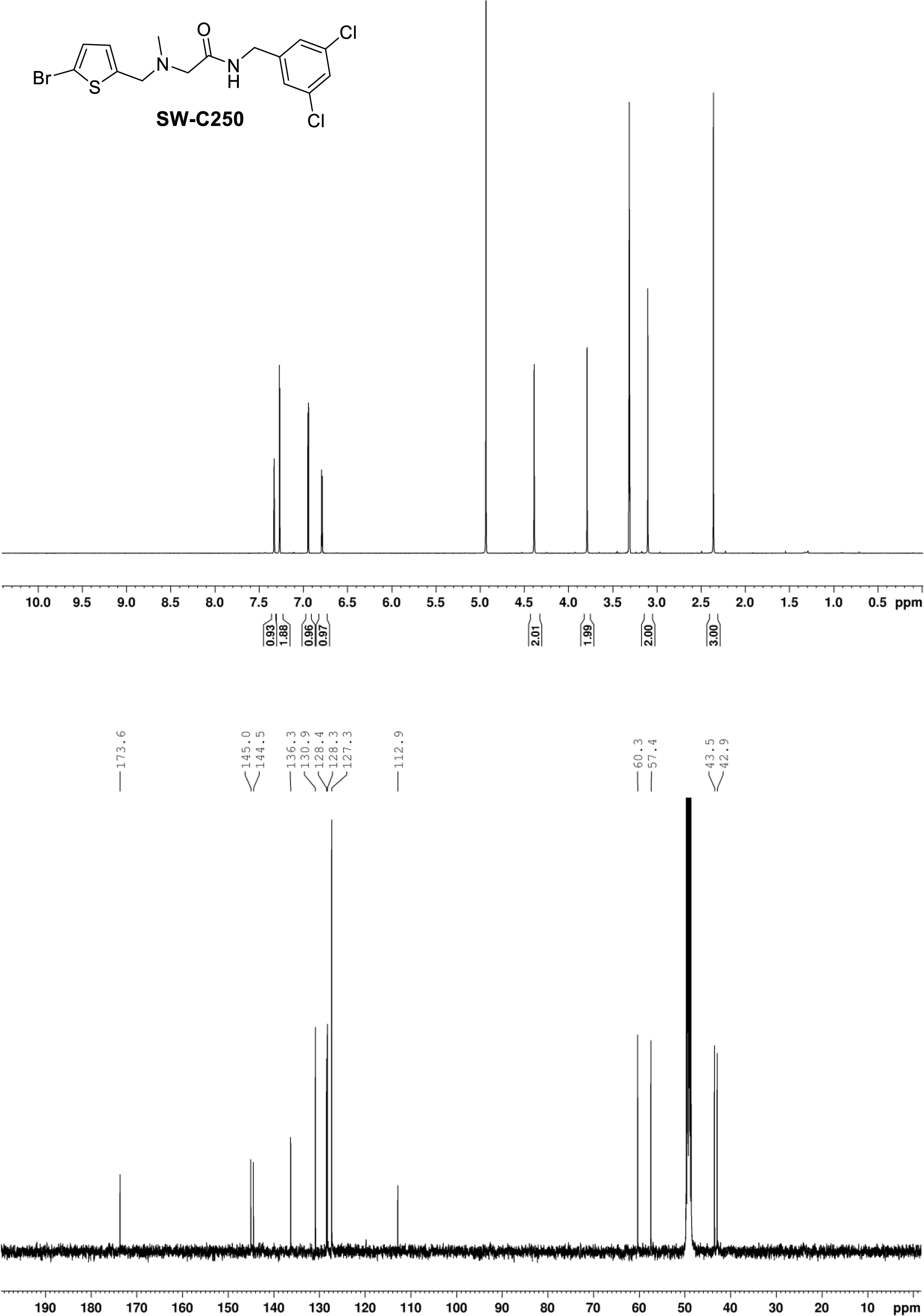

### 2-(((5-Bromothiophen-2-yl)methyl)(methyl)amino)-*N*-(4-chlorobenzyl)acetamide (SW-C202)

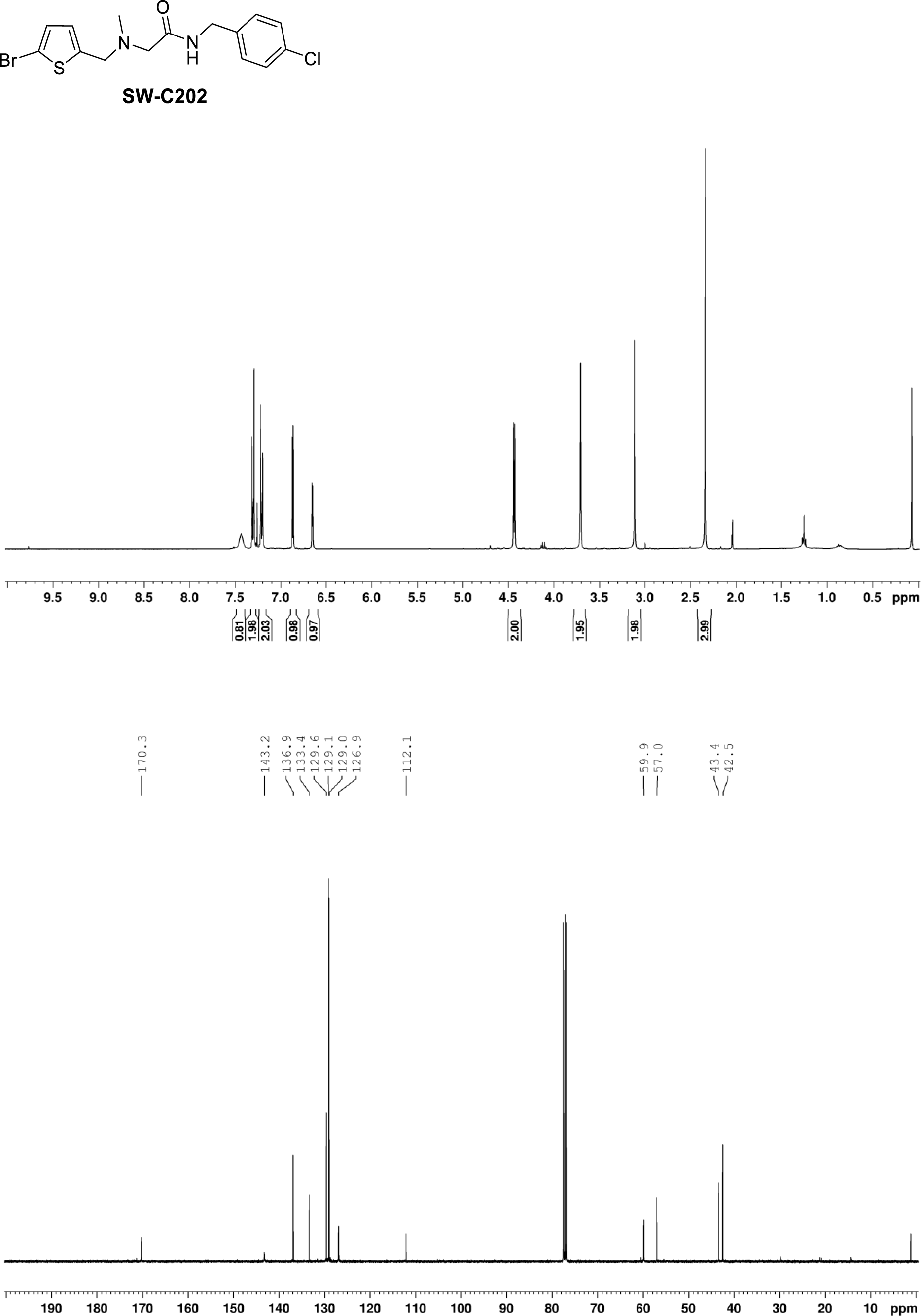

### 2-(((5-Bromothiophen-2-yl)methyl)(methyl)amino)-*N*-(3,4-dihydroxybenzyl)acetamide (SW-C170)

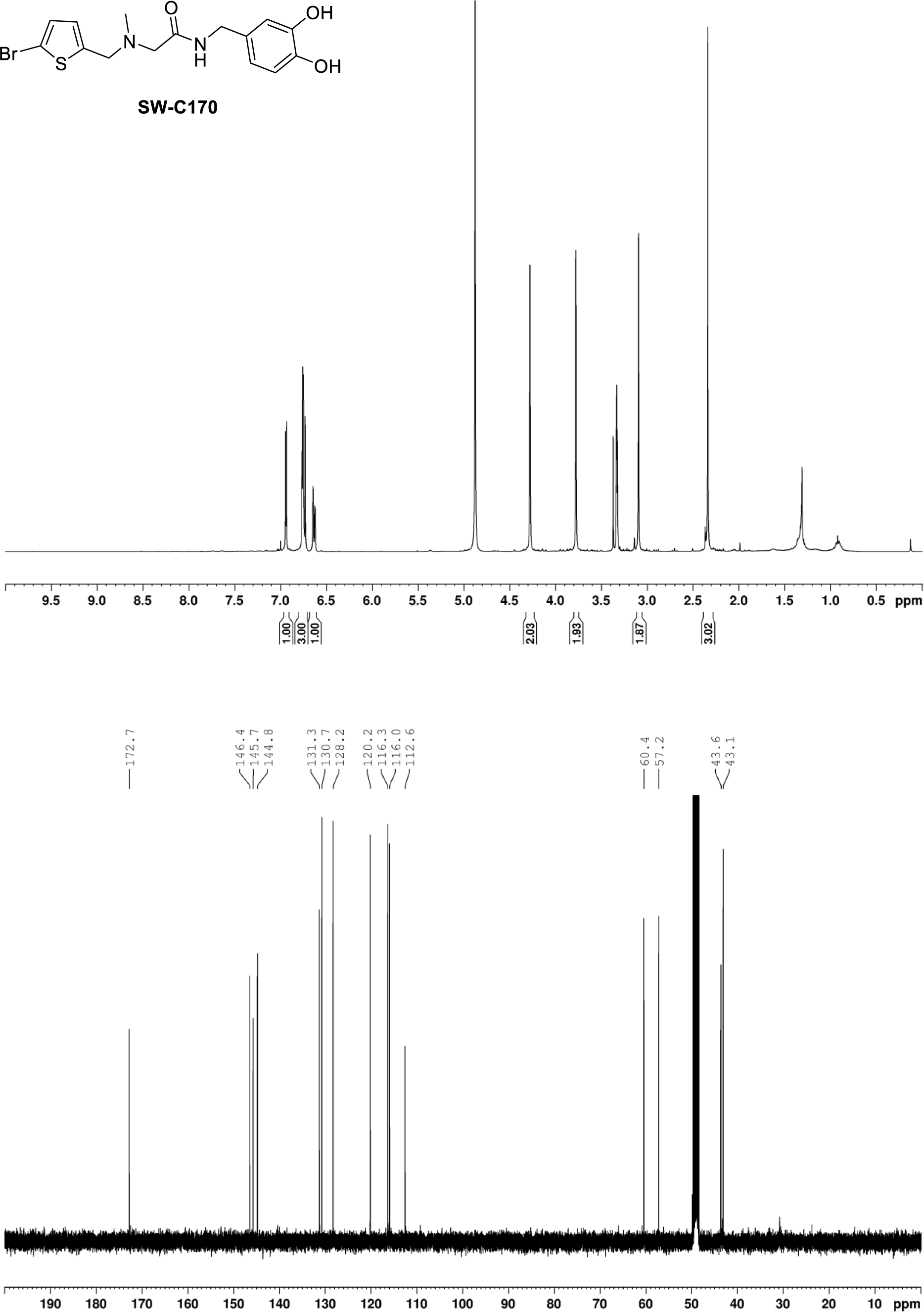

